# Complex Genomic Structural Variation Underlies Climate Adaptation across *Eucalyptus* species

**DOI:** 10.64898/2025.12.18.695066

**Authors:** Zixiong Zhuang, Scott Ferguson, Margaret Mackinnon, John Burley, Kevin D Murray, Justin O Borevitz, Ashley Jones

## Abstract

Climate change threatens global natural forest ecosystems, however, our limited understanding of adaptive genomic variation in wild forest trees restricts our ability to address this threat. *Eucalyptus* species likely harbour many climate adaptive alleles considering their long-standing evolutionary success across the Australian continent, yet their complex genomic variation and adaptive implications remain undercharacterised. Here, we present a comprehensive pangenome analysis of *Eucalyptus viminalis* using haplotype-resolved genomes and population-level long-read sequencing spanning the species’ natural distribution. We discovered complex and simple structural variants (SVs) that doubled the haploid genome length (530 Mbp haploid to 1.1 Gbp pan-genome). Landscape genomic analyses with SVs identified *CHILL1*, a 400-kb cold-adaptation locus underlined by complex structural variation. Validation across 434 accessions from three *Eucalyptus* species confirmed the large-effects of *CHILL1*, which outperforms species classification at predicting cold adaptation. These findings demonstrate that wild tree populations harbour critical adaptive alleles that can expedite climate-informed restoration.

## Main

Forests critically regulate global climate systems and are the foundation for global biodiversity ^1^. However, climate change poses a critical challenge, increasingly causing forests to degrade and release rather than sequester carbon ^2^. Understanding the adaptive capacity and genomic basis of climate adaptation in forest trees is crucial for their timely conservation, yet the fundamental understanding of genomic variation and adaptive alleles in wild tree populations remains limited ^3^. This lack of knowledge constrains our ability to identify climate-resilient genotypes and predict long-term success of natural species, which is critical for global forest restoration and management.

Among the genetic factors that may play a role in climate-resilience, genome structural variants (SVs) more than 50 bp encompassing insertions, deletions, inversions, and duplications are novel yet often complex genomic variations that could be used to identify climate-resilient genotypes. Recent pangenome studies highlighting SVs in humans ^4–7^, other primates ^8–10^, aves ^11,12^, model plants ^13,14^, crops ^15–17^, and some forestry trees with low genetic diversity ^3,18^ have revealed that SVs profoundly impact phenotypic variation and frequently underlie adaptive traits that cannot be explained by single nucleotide polymorphisms (SNPs). Therefore, further characterising SVs in naturally evolving trees represents a critical opportunity to enhance our ability at identifying climate adaptive alleles and forest resilience.

*Eucalyptus* is a highly successful genus with more than 840 species of native trees that have dominated almost all environmental niches across Australia, making it a compelling system for investigating the genomic basis of climate adaptation ^19^. *Eucalyptus* exhibit large morphological differences within species, which allowed them to be adapted across extreme environmental gradients ^20^. Among these, *Eucalyptus viminalis* is a key species to study, as it has one of the widest environmental ranges throughout the south east coast and hinterland of Australia (−10°C to 25°C; 500-1,700 mm annual rainfall, 0-2000m altitude). Moreover, it serves as a prominent habitat and food source for endemic fauna, notably koalas ^7,21^. We therefore chose *E. viminalis* as the focal study system to investigate genomic variation and natural adaptive capacity to wide environmental conditions across this broad species niche.

Previous genome-wide association studies (GWASs) have revealed remarkably high levels of genetic diversity of *Eucalyptus* species. However, these studies relied exclusively on incomplete genome sequences ^22–25^ which systematically excluded SVs. In contrast, comparative genomics across 33 *Eucalyptus* species demonstrated SVs have driven interspecific genome evolution ^26,27^. However, the prevalence, allele segregation, and adaptive significance of SVs within *Eucalyptus* populations remains unknown, but are required to further our understanding of climate adaptation^28^.

In this study, we characterise the complex genomic variation and their adaptive implications within *E. viminalis*. We performed pangenome analyses, discovered both simple SVs (sSVs) and complex SVs (cSVs), then explored their proximity and influence to known genes. By integrating range-wide long-read genomics and continental-scale environmental data, we identified a complex SV locus for which there was strong evidence of a role in climate adaptation in *E. viminalis* and two related species. Our findings highlight the importance of individual adaptive loci, rather than species or local provenance, in climate adaptation and establishes a genomic foundation for the smart design of forests in the face of rapid climate change.

## Results

### A naturally evolving *Eucalyptus* species has a diverse pangenome

To create a foundation of high-resolution resources for studying the genetic variation of *E. viminalis* across its geographic distribution, we first sought to obtain complete haplotype resolved genomes from representative individuals. We selected 10 haplotypes that we sequenced across the species range with ONT long-reads, which had coverage depths of approximately 20 to 122x and read-length N50s of 5 to 47 kbp per tree (Fig.1 a-c; Supplementary Tables 1-7). The individual with the most central geographic location was selected as the reference genome for later investigations (ACT haplotypes 1 and 2), and was therefore supplemented with additional PacBio HiFi sequencing (55× coverage, N50 20 kbp) to improve base-pair accuracy, and Hi-C chromatin conformation capture (89 million paired-end short-reads; Supplementary Note 2) for chromosome scaffolding. All *de novo* assemblies were generated with hifiasm, producing high quality haplotype-resolved genomes in 11 chromosome-scale scaffolds. Genomes were highly complete (Benchmarking Universal Single-Copy Orthologs; BUSCOs average of 97.50%), contiguous (average contig N50 24.60 Mbp, average scaffold N50 52.16 Mbp), and accurate (average genome quality value; QV of 55.75). Lastly, we generated ONT direct RNA sequencing data to support *ab initio*, homology-based annotation of the reference genome (18 million reads; N50: 1.62 kb; Supplementary Note 3).

**Figure 1.**
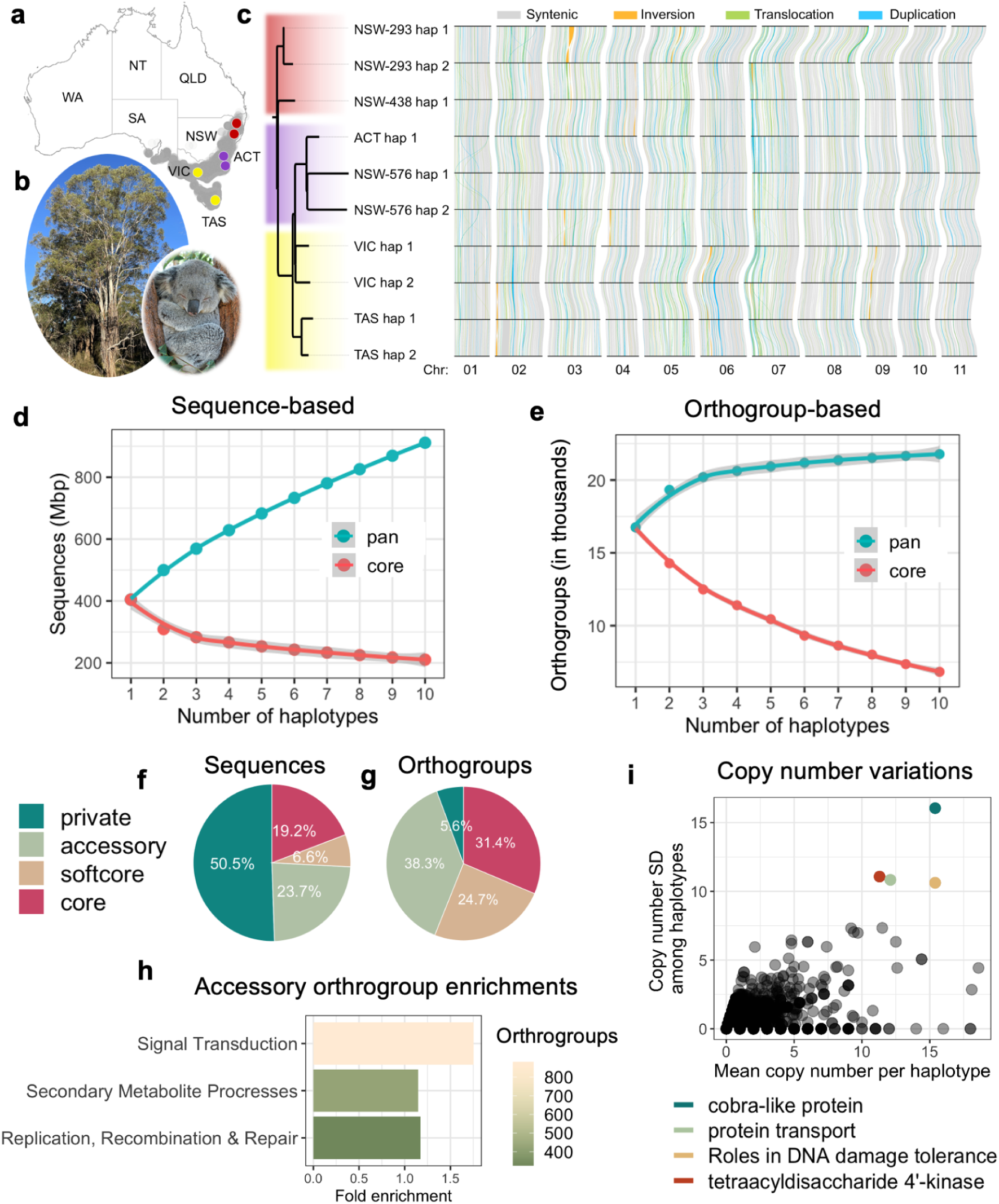
The *Eucalyptus viminalis* pangenome is undergoing rapid sequence expansion while retaining genome architecture. (a) Map of Australia showing the natural distribution of *E. viminalis* (grey shading) and the locations of trees used for genome assembly and pangenome analysis (red, purple and yellow). ACT: Australian Capital Territory, NSW: New South Wales, NT: Northern Territory, QLD: Queensland, SA: South Australia, TAS: Tasmania, VIC: Victoria, and WA: Western Australia. (b) *Eucalyptus viminalis* ssp. *viminalis* in its natural environment at Bendoura, NSW. Inset: A koala (*Phascolarctos cinereus*), the dependent arboreal marsupial. Photographs taken by Dean Nicolle and Gary Runn, respectively. (c) Phylogenetic relationships, synteny, and structural variation across 10 *E. viminalis* long-read genome assemblies. Intrachromosomal structural variants ≥20 kb are displayed. (d) Linearly increasing numbers of novel sequences in sequence-based

Using these representative *E. viminalis* genomes, we sought to understand their structural and functional genomic diversity. Through multi-genome alignments, we first revealed any large (>20 kbp) structural rearrangements that could potentially alter genome synteny. We thus mostly detected duplications and translocations that did not alter karyotype or macro-synteny (Fig. 1c). To further characterise genomic variations across assemblies, we used Pangenome Graph Builder (PGGB), sequentially investigating homologous haploblocks and then identifying variable sequences between individuals. PGGB revealed a rapidly expanding pangenome (Fig. 1d). In total, PGGB identified 1.1 Gbp of nonredundant pangenomic sequences, which is approximately 2x larger than the size of a standard single haploid genome (Fig. 1d). To further explore our findings, we additionally used Pannagram, a pangenome analysis method using different discovery principles to complement the PGGB results. This reaffirmed the PGGB results (Supplementary Note 6) and suggested 15% (or 164,999,912 bp) of these sequence variations are SVs. This level of genomic diversity is unprecedented in existing forestry and crop studies ^14,29,30^, highlighting the importance of studying naturally evolving species.

Large genomic variations such as SVs have the potential to influence the entire coding sequence of genes, causing presence/absence variations (PAVs) and copy number variations (CNVs), generating novel sources of functional diversity. Thus, we quantified gene orthogroup PAVs and CNVs across the *E. viminalis* pangenome using our *de novo* gene annotations. In contrast to the linearly increasing sequence space, orthogroups only increased logarithmically per genome input, suggesting neutrality ^31^ (Fig. 1e). Total orthogroups saturated at 21,780 compared with the 16,770 found in the reference genome, marking a 23% gain (Fig. 1e). Moreover, the orthogroup PAVs were more conserved than sequence-space variations; 56% of orthogroups were shared by almost all genomes (at least 9 out of 10), compared with the 26% of the basepairs (Fig. 1f-g). Orthogroups with variable presence/absence in the genomes could introduce sequence and functional diversification.

These accessory orthogroups were enriched in functionally critical pathways, including signal transduction, secondary metabolite processes, and replication, recombination & repair (Fig. 1h). In addition to orthogroup PAVs, CNVs within the same orthogroups also contribute to sequence and functional diversification. Among the 21,780 pangenomic orthogroups, most orthogroups (62%) had between one to five gene copies, although some reached an average of 15 copies per genome (Supplementary Note 5). Among orthogroups with highly variable copy numbers, four stood out as having the highest variances (SD >10) (Fig. 1i); these orthogroups were functionally related to DNA damage tolerance, kinase activity, and the transport and regulation of cell walls via Cobra-like proteins (Fig. 1i, Supplementary Note 5). CNVs in these orthogroups may therefore contribute to the diversification of gene function through gradual sequence divergence among gene copies. Our investigation of pangenomes and gene-level variations revealed the diversity within a naturally evolving *Eucalyptus* species, enhancing our knowledge of the genetic mechanisms driving its adaptability.

### Exploration of underlying complexity and potential source of structural variation

We sought to further investigate the complexities of our SVs and their potential source due to their adaptive and evolutionary significance. Thus, we distinguished between the often confounded simple (biallelic) structural variants (sSVs) and complex (multiallelic) structural variants (cSVs) using Pannagram, a low-bias variant calling method that allows the identification of cSVs through genome comparisons. In total, Pannagram identified 68,038 SVs from the 10 haploid genomes, including 59,428 sSVs and 26,688 cSVs. This corresponded to an average gain of 8,610 SVs per genome input (Fig. 2a). Across all 11 chromosomes, we found cSVs consistently constitute ∼20% total SVs (Fig. 2b; Supplementary Note 6). cSVs can be caused by nested events, we therefore hypothesised that larger variants would be more likely to be complex. Although we did not find evidence to support this claim in our data (Fig. 2b). SVs, especially cSVs, could hinder identification of functional trait loci, or otherwise hinder interpretations in functional studies, which merits further investigations.

**Figure 2.**
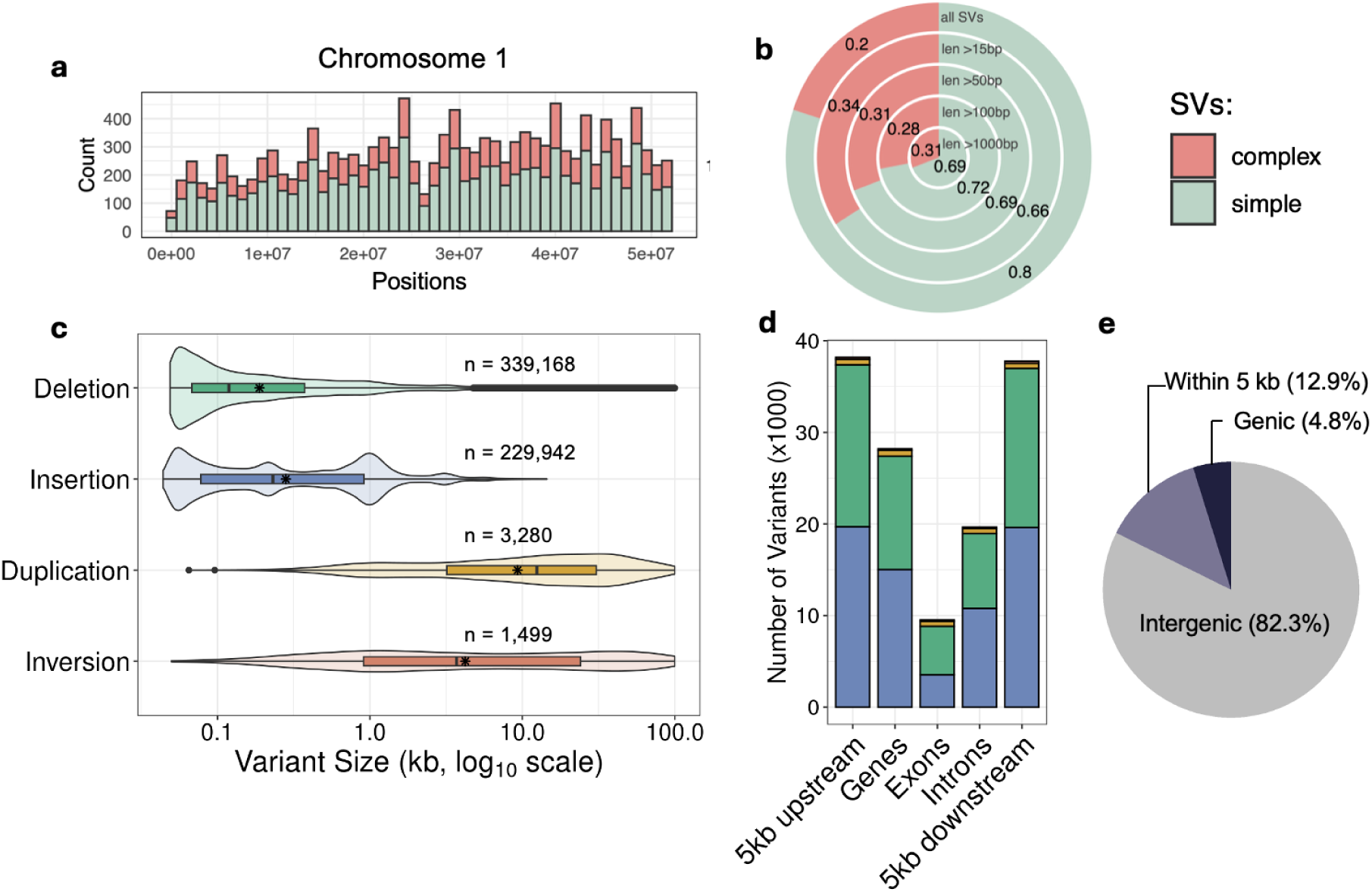
*Eucalyptus viminalis* harbours extensive structural variants influencing diverse genomic regions. (a) Simple (sSV) and complex (cSV) structural variants along chromosome 1 as an example of the results obtained for all chromosomes (see Supplementary Note 6 for the full genome). (b) Simple-to-complex SV ratio by size range. (c) Size distribution of nonredundant SVs discovered by Sniffles2. (d) SVs intersecting different genetic regions, colors represent SV types, consistent with panel (c). (e) Proportions of SVs intersecting intergenic and genic regions of the genome.

To explore the potential source of SVs, we tested the association between SVs and transposable elements (TEs), a common source of genomic variation ^32^. Across the genome, we found SVs were 15% more likely (p < 0.001, z = 141.27) to be near TE regions than by random chance. We next investigated the sequence homology between sSV sequences and the *de novo* annotated TE sequence library (Supplementary Note 7). In doing so, we found that 13% of sSVs have strong homology with TE sequences (>85% sequence similarity at >85% overlap). This suggested that some SVs may be derived directly from TE activities. Although the source of the majority of SVs remained unclear. Lastly, we investigated open reading frames within SV sequences that could transcribe intact peptide sequences. In doing so, we discovered a novel, 1,031-peptide Pol poly-protein, which was both associated with long tandem repeat (LTR) retrotransposons, and a cluster of 781 large SVs (6 - 12 kb) in our data (Supplementary Note 7). We confirmed active expression of this Pol protein in leaf tissues (TPM: 31; 566 transcripts), which suggested ongoing LTR expansions within *E. viminalis* (expanded in Supplementary Note 7). These results provided novel insight into how plant SVs arise, highlighting a key area for future studies.

### Structural variants influence diverse genomic loci

To increase the power of identifying functionally important structural polymorphisms, we sampled more deeply across the native range. Thus, we generated a range-wide ONT long-read sequencing dataset from 71 (142 haplotypes) *E. viminalis* individuals (Supplementary Table 7). From these, we selected 34 (68 haplotypes) samples with moderate coverage (as determined by Supplementary Note 8) to discover SVs using Sniffles2. Initially, Sniffles2 identified 883,116 SVs, which were then consolidated into 573,889 nonredundant SVs through singleton filtering (Supplementary Table 9). This result corresponds to an average gain of 8,439 SVs per haploid genome input, which is remarkably consistent with the genome-based results from Pannagram (8,610 SVs per haploid genome). These SVs covered 38% of the reference genome, and the remaining genome positions could therefore be considered structurally conserved (Supplementary Table 10; Supplementary Note 9).

Next, we characterised the size and frequency of SVs identified in the *E. viminalis* range-wide dataset, along with their genomic proximity to genes (Fig. 2c-d). Among different types of SVs, insertions and deletions were the most frequent (>99% SV loci), with median sizes of 231 bp and 119 bp respectively. In contrast, inversions and duplications were much larger (median sizes: 3 kb and 12 kb respectively; Fig. 2c), but also much rarer (<1% SV loci) than insertions and deletions. Many of these SVs (133,379) had the potential to influence genes, being within a 5 kb proximity (12.9%) or directly intersecting genes (4.8%), which were predominantly insertions and deletions due to their high frequencies (Fig. 2d-e). Interestingly, the SVs that were within 5 kb or intersecting genes were enriched in several disease resistance gene pathways (Supplementary Note 9). In particular, (tandem) duplications were enriched sixfold for events directly affecting exons (6% vs. 1% genome-wide), indicating that exon duplication is better tolerated than deletions or inversions are. The main findings in SV characteristics aligned closely with existing literature ^33,34^. In particular, exon duplication appears to be common in trees, potentially related to protection against pathogen attacks across their often long lifespans ^35^.

### Spatial and environmental gradients explain neutral genomic structural variations

Since having a strong underlying genomic structure complicates the identification of functional alleles, we first evaluated population structure driven by neutral factors, including geographical barriers, and reproductive isolation driven by environmental factors. For this analysis, we expanded the number of test individuals from 34 (68 haplotypes) to 49 (98 haplotypes; Fig. 3a), by lowering the ONT coverage requirements (from >12x to >8x; expanded in Supplementary Note 8). We then genotyped the 573,889 SVs discovered previously in these geo-referenced individuals, which generated diploid genotypes. Minor allele frequencies showed a broad spectrum of SV allele frequencies, with duplications as the most common type (Fig. 3b). Unpurprisingly, pairwise genomic distances derived from SV genotypes covaried with geographical distances across the 620 km sampled range due to travel limits of pollinators (Fig. 3c). However, no discrete population structure was found, suggesting the gene flow within this species is likely continuous. Additionally, principal component (PC) analysis showed PC1 only explained 5.49% of total variation, suggesting neutral variation only represented a small portion of total genomic variance (Supplementary Note 10).

**Figure 3.**
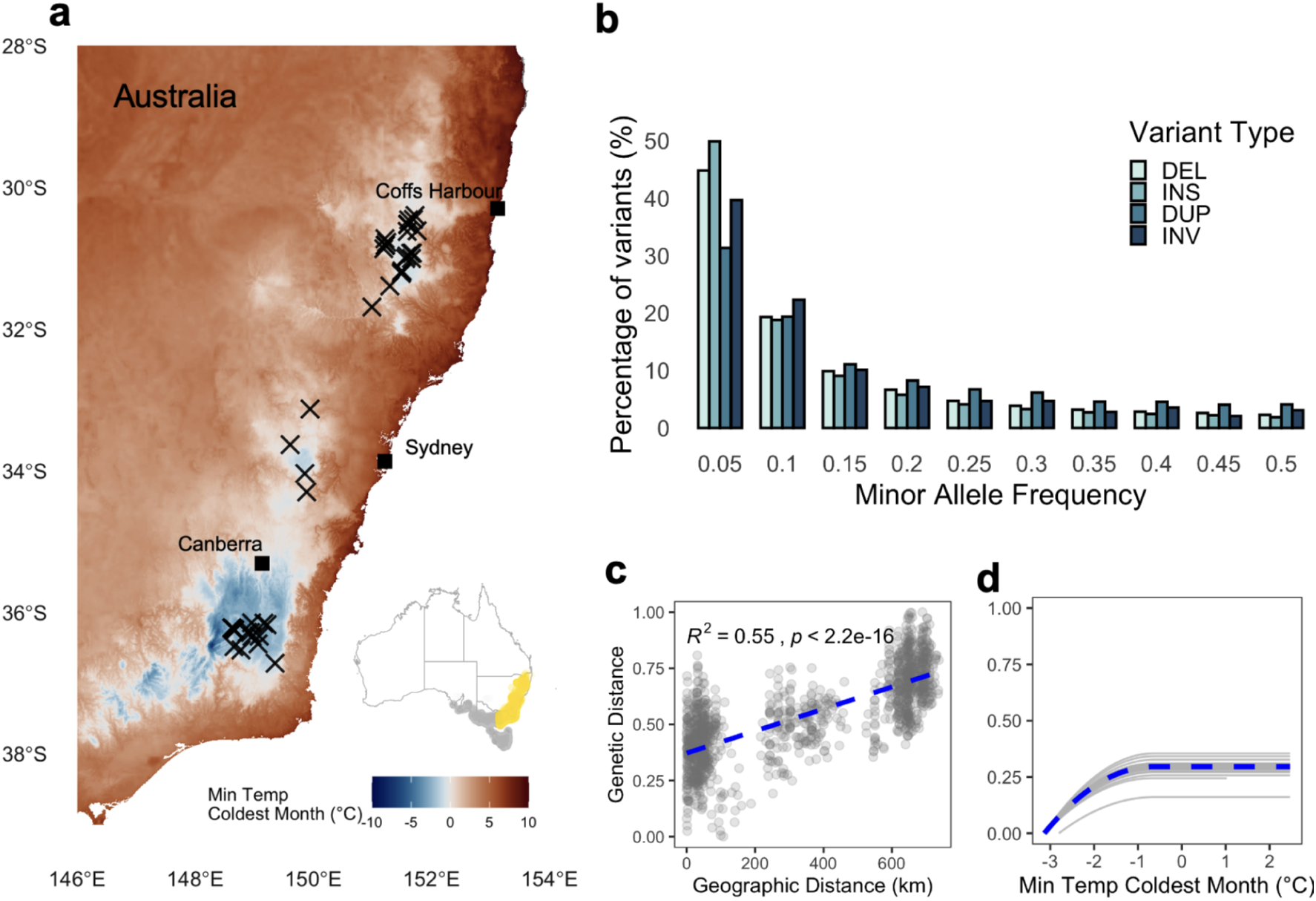
Intraspecific SV alleles of *Eucalyptus viminalis* segregate across its geographic and environmental ranges. (a) Spatial distribution of *E. viminalis* samples from which long-read data was obtained (49 samples; 98 haplotypes). Inset map of Australia highlights the sampled range (gold) relative to the natural distribution of *E. viminalis* (grey). (b) Minor allele frequency spectrum of four structural variant (SV) types (deletions, insertions, duplications and inversions). (c) Covariation between pairwise geographic distance and pairwise genetic distance calculated with SV genotypes (linear regression; R^2^ = 0.55, P < 0.01). (d) Covariation of genetic distance with minimum temperature of the coldest month (BioClim6) beyond the covariation explainable by geography alone, using generalised dissimilarity modelling. The dotted line shows the spline curve fit obtained using all the samples. The grey lines show cross-validations by iteratively leaving one sample out.

In addition to genographic segregation, genetic structure can also be driven by environmental factors, especially ones that potentially influence phenology ^24,36^. In particular, temperature extremes may act as selective pressure on the timing and success of plant reproduction. To explore the potential environmental drivers for neutral variation, we extracted long-term environmental data (BioClim variables), from the WorldClim database using the locations of each of the 49 trees. Secondly, we fitted generalised dissimilarity models to separate environmental gradients from their covariation with geography. In doing so, BioClim6 (minimum temperature of the coldest month) stood out as the climate variable that offered the strongest additional explanatory power beyond geography (Supplementary Note 10; Fig. 3d). This aligned with the known ecological niche of *E. viminalis,* a species recognised for its cold tolerance as a historic candidate for reforestation and forestry plantations in temperate regions ^37^. These results suggest that cold temperature extremes are a driver of genetic divergence in *E. viminalis*, although the underlying adaptive alleles are yet to be determined.

### Structural variations confer putative climate-adaptive loci in *Eucalyptus*

Due to the population structure explained by cold environments, we hypothesised that *E. viminalis* would harbour cold-adaptive alleles that enabled this apparent environmental covariation. To identify these potentially cold adaptive alleles, we performed a genome-wide association study (GWAS) to associate BioClim6 values with SV genotypes using Genome-wide Efficient Mixed Model Association (GEMMA). To control for demography-driven neutral variations as characterised in the previous section, we included a genome-wide SV derived genomic relationship matrix to correct for the underlying correlation structure. We thus identified several significant loci that are strongly associated with BioClim6 (Bonferroni threshold; -log10 P-value >7), which represent cold temperature extremes. In total, we identified 10 significantly associated SV marks across four different chromosomes (Fig. 4a; Supplementary Table 12). This revealed multiple candidate loci implicated in cold climate adaptation, that we named *CHILL*. Among these, we identified *CHILL1* as the most significant, with the -log10(P-value) of 12.5, almost twice the stringent Bonferroni threshold (Fig. 4a). We also identified other significant loci of interest (notably *CHILL8*; expanded in Supplementary Note 12 and Supplementary Table 13). We hereinafter primarily focus on *CHILL1*.

**Figure 4.**
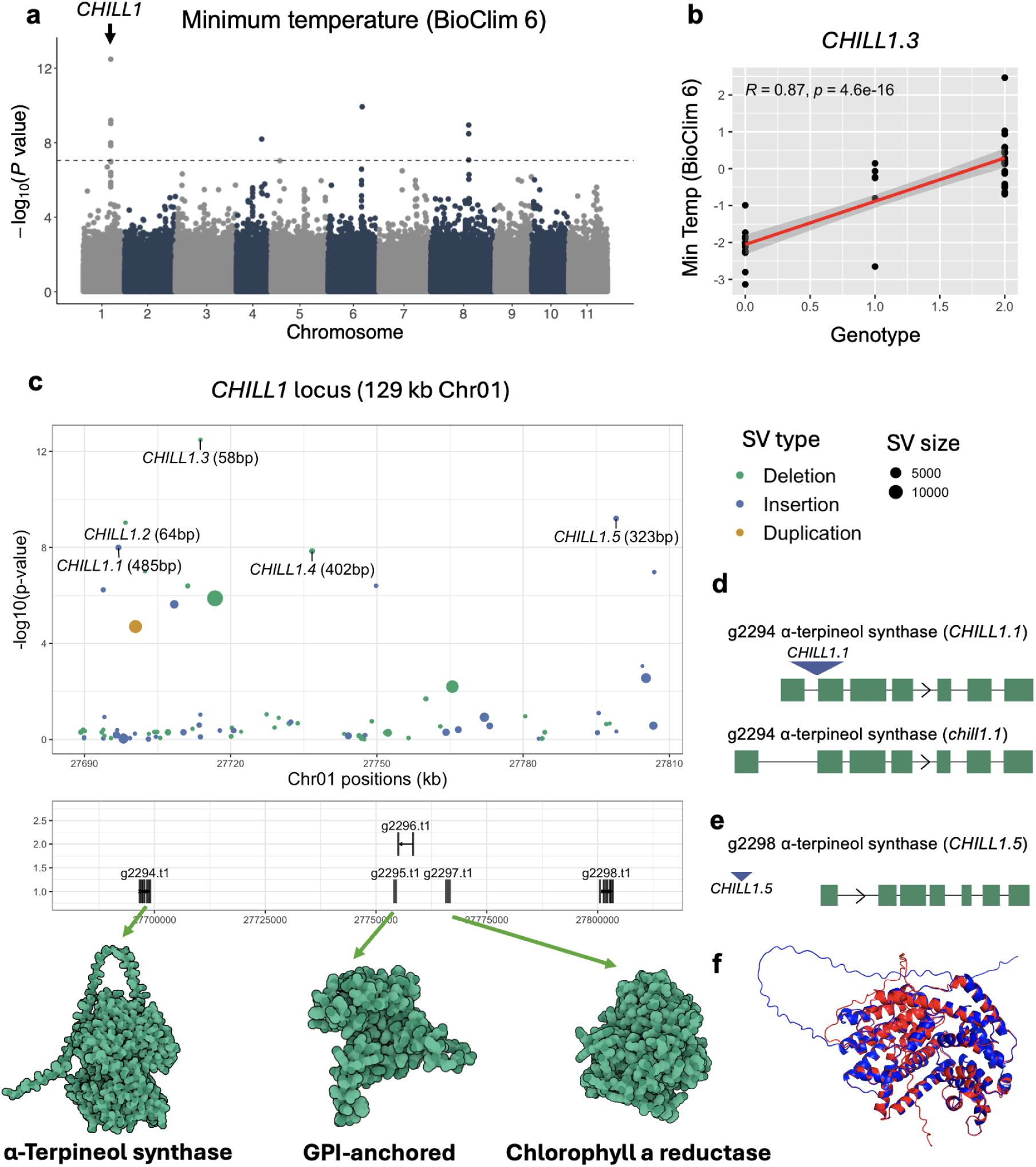
Discovery of *CHILL1*, a cold-adaptation locus driven by structural polymorphisms. (a) Manhattan plot of the SV-based genome-wide association study (GWAS) with the minimum temperature of the coldest month (BioClim6). (b) Genotype-environment associations at the *CHILL1* locus. (c) Zoomed-in 129 kb view of the *CHILL1* locus, including putatively associated SVs (upper panel), implicated genes (lower panel), and corresponding protein structures. Alphafold structure of three genes implicated in cold adaptation, including ones encoding an α-terpineol synthase protein, glycosylphosphatidylinositol (GPI) anchored protein, and a 7-hydroxymethyl chlorophyll a reductase. (d) The *CHILL1.1* insertion occurs at the first α-terpineol synthase gene, expanding the first intron of the *chill1.1* allele. (e) The *CHILL1.5* insertion occurs in the promoter region of the second α-terpineol synthase gene. (f) Comparison of Alphafold-predicted structures between the two α-terpineol synthase protein coding genes (g2292 and g2298).

The SV-defined *CHILL1* spanned across a 129 kb window between 27.7 and 27.8 Mb on chromosome 1 (Fig. 4c). This window was defined by five significant SVs (*CHILL1.1-CHILL1.5*), which were all small insertions or deletions (58-323 bp; Fig. 4c). Among them, *CHILL1.3* was most strongly associated with BioClim6. The allelic state of *CHILL1.3* suggested that plants with the alternate allele could only tolerate, on average, minimum temperature of 0°C; whereas trees with the reference allele could tolerate as low as -2°C, thus requiring much more substantial cold and frost tolerance for plants to survive (Fig. 4b). The *CHILL1* locus implicated several functionally important genes for further investigation (Fig. 4c; Supplementary Note 11; Supplementary Table 14). These included two gene copies encoding Glycosylphosphatidylinositol (GPI)-anchored proteins. Transcript expression of these genes was not detected in *E. viminalis* leaves, which may reflect their *Arabidopsis* homologue (At1G61900) being primarily expressed in flowering tissues and seeds ^38^ (Supplementary Note 11). Another gene encoding 7-hydroxymethyl chlorophyll a reductase, was also present in *CHILL1* and was expressed in *E. viminalis* leaves (TPM: 13.42). Both GPI-anchored proteins and 7-hydroxymethyl chlorophyll a reductase have been linked to plant cold acclimation in previous studies ^38,39^. Additionally, two genes encoding α-terpineol synthase proteins were present, one being directly intersected by *CHILL1.1* (a 485 bp insertion in the first intron), the other was in close proximity to *CHILL1.5* (a 323 bp insertion in close the suspected promoter region) (Fig. 4d-f). Neither gene was detected as expressed. Together, our findings highlight SV polymorphisms as a promising avenue for discovering putative climate adaptive loci, where SVs may confer functional adaptation in *E. viminalis* by influencing functionally important genes.

### *CHILL1* is a large-effect climate-adaptive locus that is conserved across *Eucalyptus* species

We next sought to validate the large climate adaptive effect of *CHILL1*. To do so, we generated Illumina whole-genome short-read datasets and called SNPs from 434 samples across three species; including 176 *E. viminalis* samples, 138 *E. dalrympleana* samples, and 120 *E. rubida* samples. These three species were closely related (*Eucalyptus* subg. *Symphyomyrtus* sect. Maidenaria), and were sampled from the same geographical range to capture similar selective pressures (Fig. 5a; Supplementary Note 13). Because we had identified *CHILL1* as a large-effect locus covering 129 kbp, we expected that many linked SNPs would exist within this region, allowing the validation of this locus ^40^. As predicted, SNP data from these three species confirmed that the *CHILL1* region contained numerous tightly linked SNPs (Fig. 5b). This resembled a recombination-reduced haploblock (known as a supergene) of ∼400 kb with three distinct clades of samples clustered by their SNP genotypes (Fig. 5b). Moreover, SNPs within the haploblock were strongly correlated with SV genotypes, especially in the subregion where *CHILL1.1-CHILL1.5* were located (chromosome 1: 27.7-27.8 Mb; Supplementary Note 14). Thus, these linked SNPs enabled the determination of *CHILL1* genotypes in the short-read dataset, which we then used to evaluate *CHILL1*-BioClim6 associations across species.

**Figure 5.**
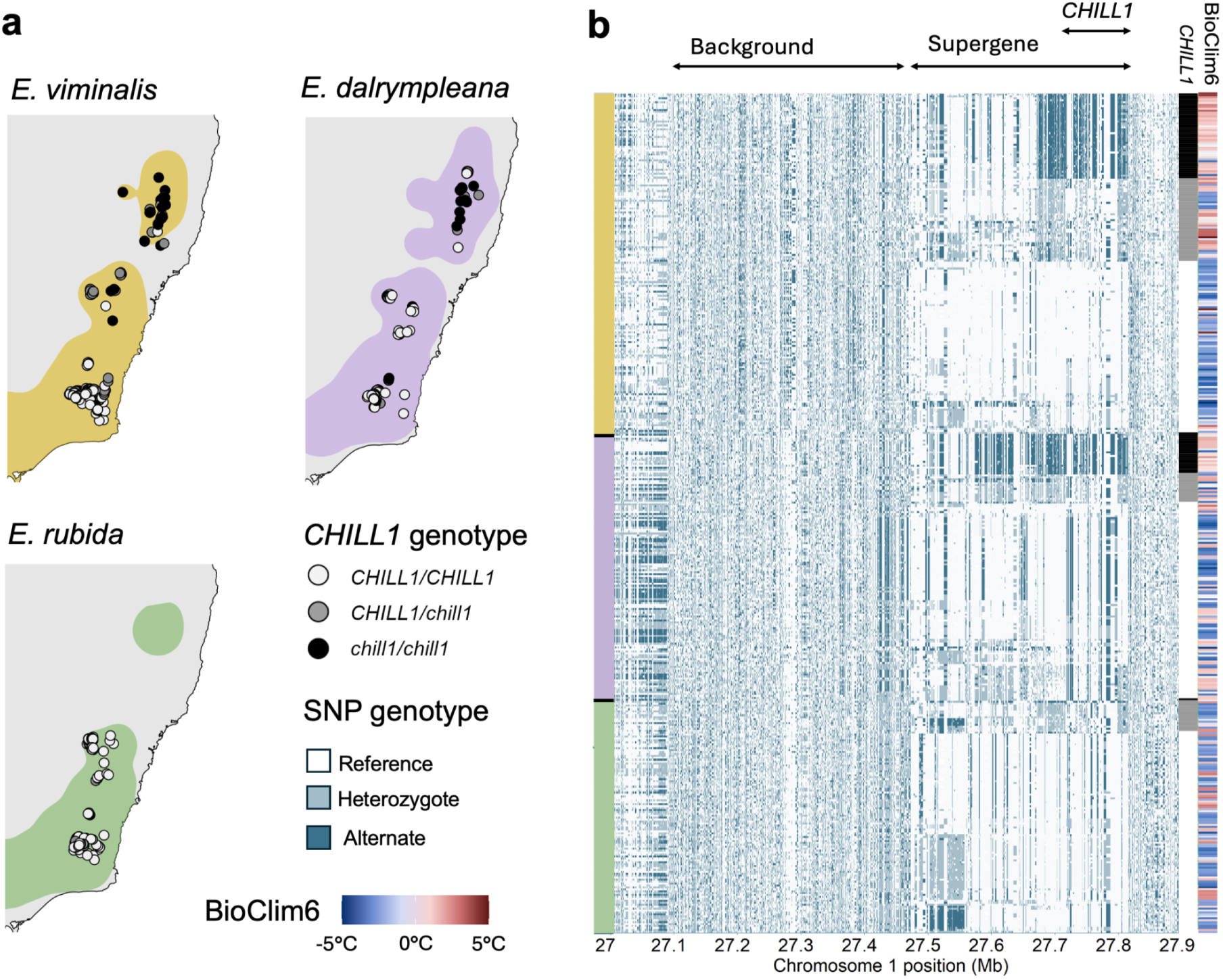
Validation of *CHILL1* demonstrates that it is a conserved supergene across-species. (a) Spatial distribution of the *CHILL1* haplotype across three *Eucalyptus* species; *E. viminalis, E. dalrympleana*, and *E. rubida*. Species distribution polygons represent kernel density–based approximations of occurrence records from Atlas of Living Australia. (b) Short-read SNP genotypes in the larger *CHILL1* region (chromosome 1: 27-28 Mb) for 434 trees across the three species.

Among the 434 samples, *CHILL1* was a common allele, carried by 75% of *E. viminalis* samples, 84% of *E. dalrympleana* samples, and 99% of *E. rubida* samples (Supplementary Table 16). Absence of the *CHILL1* allele was most prevalent in warmer northern regions, where selective pressure for cold adaptation is reduced. Remarkably, *CHILL1* allele frequencies were strongly differentiated across geographic zones, both within and across species as expected for an adaptation locus (Fig. 5; Supplementary Note 14). Multiple regression modelling of latitude, elevation, *CHILL1* genotype and species provided an estimated *CHILL1* effect size of 1.59°C (P = 10^-16^), which was largely consistent with the SV-based results (Supplementary Table 15). Strikingly, the geographic effect of shifting 111 km in latitude and 100 m in elevation were equal to the genetic effect of the *CHILL1* allele, while the overall species effect was minimal (Supplementary Table 15). Additionally, PC analyses of SNPs demonstrated that *CHILL1* haplotype explained 65% of the total genetic variance in the locus, much higher than the 9.5% variance explained by species labels (Supplementary Note 14). These results not only confirmed *CHILL1* as a large-effect cold-adaptation locus, but also demonstrated that a single large-effect allele could predict environmental suitability much better than species classification.

### *CHILL1* is a complex locus shaped by transposable elements and complex structural variation

Following the discovery and extensive validation of *CHILL1*, we sought to use the newly assembled haplotype genomes and their annotations to further characterise this important cold-adaptive locus. We first investigated the rearrangements between *CHILL1* and *chill1* haplotypes, selecting ACT hap1 and NSW-293 hap1 as representative assemblies that were gapless across this region. This revealed complex rearrangements within the *CHILL1* region defined by SVs (chromosome 1: 27.7-27.8 Mb; Fig. 6a). Interestingly, the broader region remained largely syntenic with no inversion. Synteny and lack of inversions were further confirmed by pair-wise comparisons of all our haplotype-resolved genomes (n = 10), in the broader region defined by synteny-resolved orthogroups on the reference genome (chromosome 1: 27-28 Mb; Supplementary Note 15).

**Figure 6.**
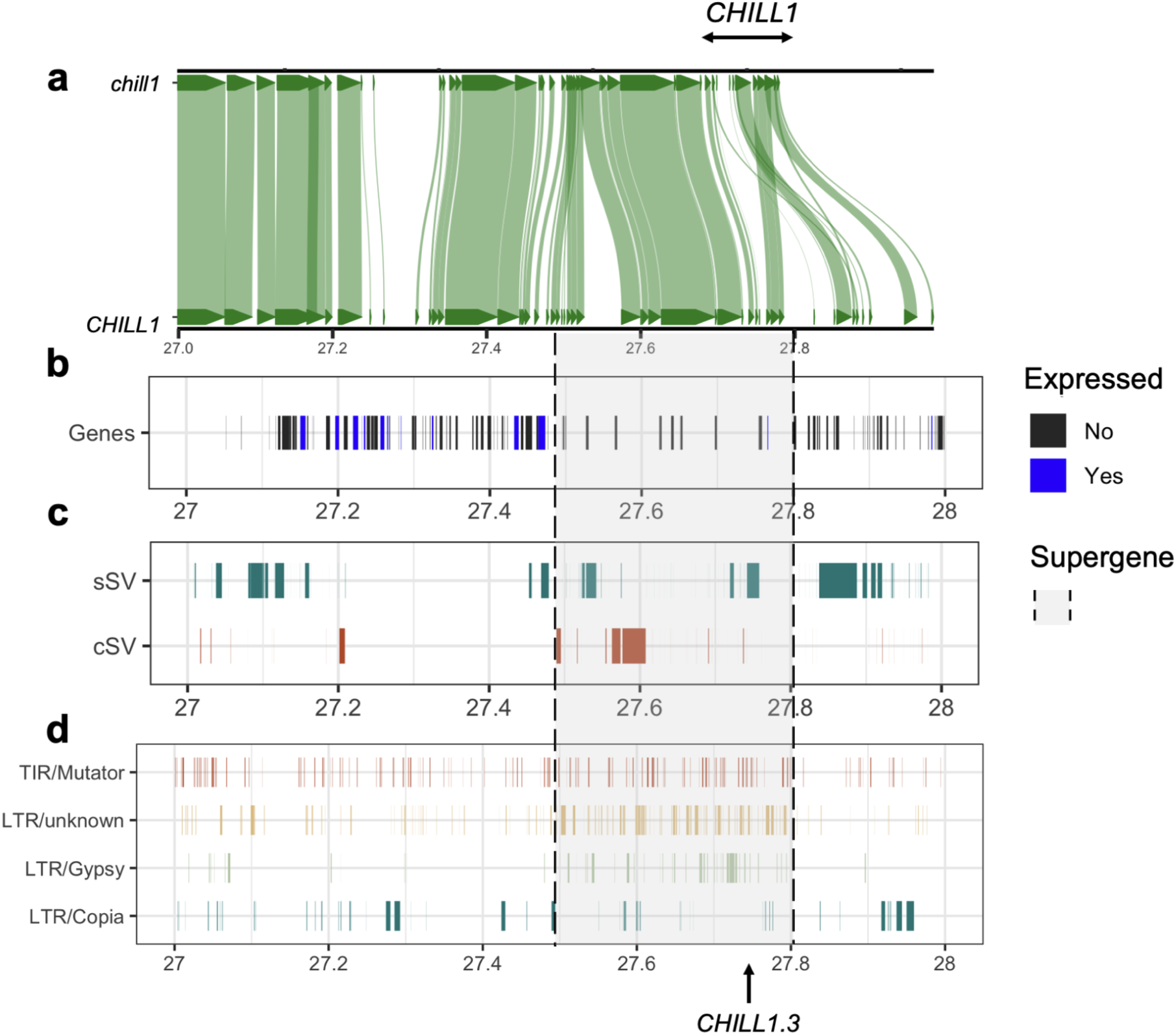
*CHILL1* is shaped by complex genomic features, including genes, structural variants, and transposable elements. (a) Homology plot of two *Eucalyptus viminalis* haplotypes in the wider *CHILL1* region (chromosome 1: 27-28 Mb) (b) Gene models annotated for the reference genome with expression validated by ONT direct RNA sequencing (≥1 transcript per million). (c) Simple and complex structural variants (sSVs, and cSVs) identified by Pannagram using all 10 haplotype-resolved assemblies of *Eucalyptus viminalis*. (d) Transposable element annotations across the *CHILL1* region.

Across the broader *CHILL1* region, we observed numerous genes (many being expressed in leaves), sSVs and cSVs, and multiple classes of TEs (Fig. 6b-d). These TEs belong to different classes of LTRs (retrotransposons) and terminal inverted repeats (TIRs; DNA transposons), therefore likely resulting from multiple previous TE expansion events. We observed varying TE densities in the region, where most TEs were concentrated in the supergene subregion, potentially suggesting a role of TE activity in the development of this locus (Fig. 6c). Additionally, multiple cSVs were present in *CHILL1* and the border region, making their detection challenging through traditional read-mapping methods. These results suggest that *CHILL1* likely arose from complex evolutionary events in the past, although the apparent lack of recombination in this region was not clear. In summary, *CHILL1* is a structurally and evolutionarily complex locus, highlighting the importance of using a diversity of data types and analyses in functional studies of naturally evolving plant species.

## Discussion

Wild forest trees harbour adaptive alleles valuable for forest protection under climate change, yet prior studies lacked the genomic resources to identify complex genomic variation and their adaptive potential. Here, we established the most comprehensive genomic resources for any naturally evolving *Eucalyptus,* including *de novo* assembled haploid genomes, direct RNA sequencing, a range-wide long-read dataset, and a cross-species short-read dataset. Using these new resources, we revealed vast genomic diversity, including novel complex structural variation that underlies climate adaptation across *Eucalyptus* species.

We observed a doubling of pangenome size from a single haploid reference of 540 Mbp to 1.1Gbp with 10 haploid genomes. This 103% sequence gain is significantly larger compared to species such as humans (6% per 116 haps) ^41^, *Arabidopsis* (31% per 27 haps) ^42^, and oak (79% per 22 haplotypes) ^43^. In our study, approximately 15% of genomic variations were SVs. Many of these SVs may have functions. Using a SV-GWAS approach, we identified the *CHILL1* locus that predicts cold adaptation with one of the largest effect-size for a single locus (2°C). Our short-read validation shows *CHILL1* is a cross-species cold adaptation locus with equal effects as range-wide migrations (in latitudes and altitudes), while the species effects are minimal. This resolved the previously unexplained large variance in cold-tolerance within *E. viminalis* (Jahromi, 1982; Paton, 1972) and makes *E.viminalis* as a competitive candidate for reforestation due to its lack of tradeoffs between cold-tolerance and growth rates. Although SV-GWAS has become a popular approach in (super)pangenome studies of agricultural crops in recent years ^3,29,44^, few have discovered alleles with such effects.

Several genes implicated by *CHILL1* provide novel insights into plant abiotic stress tolerance. GPI-anchored proteins are predicted to stabilise apoplastic membranes in reproductive tissues under frost ^38^. Interestingly, we found the GPI-anchored protein in the *CHILL1* locus belonged to the orthogroup (cobra-like proteins) that had the highest variance in CNV among all 21,780 orthogroups, which suggests high diversity and specialisation of these genes (Fig. 1i). Two α-terpineol synthases may mediate ABA/SA-independent stomatal closure in response to temperature and light cues ^45–47^, which may work in tandem with the 7-hydroxymethyl chlorophyll a reductase that adjusts chlorophyll a/b ratio to protect photosystem I under chilling conditions in trees ^39^, including *Eucalyptus* ^48^. These findings offer novel research avenues into plant adaptation strategies, including how shifts in photosynthetic biochemistry relate to whole-plant survival and how membrane fluidity influences tree fecundity under cold stress.

Beyond its large effect and functional candidates, *CHILL1* resides within a ∼400 kb supergene, similar to supergene architecture reported in sunflowers ^49^ and deer mice ^50^. However, inversions were proposed as the sole mechanism for supergenes, yet no inversion was detected near *CHILL1*. Many mechanisms could contribute to the development of this locus such as the influence of TEs. *E. viminalis* likely has previous and on-going bursts of TE expansion events (Supplementary Note 7), and that many classes of TEs are enriched in the *CHILL1* supergene region (Fig. 6d). TE-related insertions and deletions can modify genomic sequence repeats ^51^, methylation ^52^, and chromatin accessibility ^53^ that creates functional differences for selection to act upon. Whether this supergene is ancestral or derived remains unresolved and warrants future investigations.

In conclusion, we established pangenomic resources for naturally evolving *Eucalyptus* trees, including *de novo* genomes, SVs and SNPs. We demonstrated novel genomic polymorphisms like SVs can be used to define large-effect climate-adaptive loci. *CHILL1* is a cross-species cold adaptation allele with immediate real-world impact of guiding forest restoration, although further study is needed to resolve the mechanistic functions governing *CHILL1* and the larger supergene. This work establishes a strategic pathway to discover and utilise natural adaptability in forest trees for safeguarding global forests and biodiversity under accelerating climate change.

## Methods

### *Eucalyptus viminalis* reference genome assembly

A *de novo* telomere-to-telomere *Eucalyptus viminalis* genome was assembled for use as the reference genome in all analyses in this study. Fresh leaf tissue of *E. viminalis* (subsp. *viminalis*) tree was sourced from the Australian National Botanic Gardens, Canberra Australia (accession number CBG 7902535.8, located in section 111). Tissue was kept cool and hydrated until cryogenically stored at -80°C as soon as possible. Approximately 10 g of leaf tissue was ground to a fine powder using a mortar and pestle cooled under liquid nitrogen, and DNA was extracted according to ^54^. DNA was subsequently size selected for fragments ≥ 20 kb using a BluPippin (Sage Science). A Pacific Biosciences (PacBio) SMRTbell DNA library was prepared (SMRTbell Express Template Prep Kit 3.0) and sequenced on the Sequel II platform using an 8M SMRT cell. High-fidelity (HiFi) circular consensus reads were generated using DeepConsensus (version: 1.2.0) ^55^. To obtain ultra-long reads, an Oxford Nanopore Technologies (ONT) native DNA library was prepared (ligation kit SQK-LSK114) and sequenced on the PromethION P24 platform using a FLO-PRO114M R10.4.1 flow cell. When sequencing declined (low active pore count), the flow cell was treated with ONT Flow Cell Wash Kit (EXP-WSH004), then re-primed and reloaded with more libraries following the manufacturer’s instructions. Sequencing data (pod5) was basecalled to fastq with ONT Dorado (version: 0.3.4), using the super accurate model. A chromosome conformation capture Hi-C library was prepared with a Phase Genomics Proximo Hi-C Plant Kit (version 4, document KT3040B) and sequenced on an Illumina NovaSeq 6000 using an S4 flow cell with a 300 cycle kit (150 bp paired-end sequencing).

All PacBio HiFi reads were used in assembly (≥ Q20). ONT reads were filtered for ≥ Q7 and ≥ 55 kb using NanoFilt (version: 2.8.0) ^56^. Hi-C reads were filtered with Trim Galore (version: 0.6.10) ^57^, to remove any Illumina adapters and validate the read pairs. The *de novo* genome assembly was performed with Hifiasm ultra-long (UL) (version: 0.19.6-r597) ^58^, incorporating the PacBio HiFi, ONT ultra-long (--ul) and Hi-C reads (--h1 --h2).

After assembly, Hi-C contact maps were independently created for each haplotype using Juicer (version 1.6) ^59^ and subsequently scaffolded with 3D-DNA (parameters: --editor-repeat-coverage 5 -r 1; version: 190716) ^60^. Juicer used BWA-MEM for read alignment (version: 0.7.17) ^59^. After initial scaffolding, each haplotype was visually inspected and, when necessary, manually edited using Juicebox (version 1.11.08) ^61^.

For quality assessment of the genome, telomeres, single copy genes, and base accuracy was calculated. The telomere sequence for *E. viminalis* was discovered using tidk v0.2.65 ^62^ (using explore parameters: --distance 0.02 --minimum 5 --maximum 12; version: 0.2.4). The resulting list of candidate telomere k-mers was then processed with seqtk telo (version 1.4; https://github.com/lh3/seqtk), identifying the true telomeric kmer sequence and annotating all chromosomes for their telomere regions. Assembly completeness was assessed by BUSCO with compleasm v0.2.7 ^63^, using the Eudicotyledons OrthoDB v12 dataset ^64^. To assess base accuracy of the genome, yak v0.1 (r56) was used to compare k-mer spectra (k=31) between the assembly and sequencing reads to calculate an assembly quality value (QV) ^65^.

### Additional T2T genome assemblies

In order to confirm the conservation of karyotype, we sampled two more interstate samples for T2T genome assembly. These included a sample from the Tasmanian Herbarium, Sandy Bay, Tasmania (first draft genome reported in Ferguson et al., 2024) and a sample from Trentham, Victoria (accession GDH178). HMW DNA was extracted, size-selected (≥ 20 kb), and sequenced using FLO-PRO114M R10.4.1 PromethION flow cells as described above. Basecalling was performed with Dorado v7.6.8 (sup v4.3.0, 400 bps). Raw reads were filtered for ≥ Q20 and ≥ 20 kb using chopper v0.8.0 ^56^. *De novo* genome assemblies were performed with hifiasm (ONT) v0.25.0-r726 (Cheng et al., 2025) with ONT simplex reads (--ont), adding the telomere sequence (--telo-m AAACCCT), and dual scaffolding (--dual-scaf). Only chromosome two remained broken. Final scaffolding and ordering of chromosomes was performed with RagTag v2.1.0 ^66^, using the *E. viminalis* ACT genome as the reference. Quality assessment of the genomes was performed as described above for the ACT *E. viminalis* reference.

### Haplotype-resolved genome synteny analyses

To analyse the conservation of genome architecture and identify structural variants across *E. viminalis*, the highest quality genome assemblies were selected for phylogenetic and comparative genomic analyses. Samples were collected from four states and territories across Australia to capture geographic and genetic diversity. Only haplotype-resolved, chromosome-scale assemblies with > 92% BUSCO completeness were included to ensure accurate phylogenetic inference and detection of syntenic regions and structural rearrangements. These assemblies were subsequently used for pangenome construction and analysis.

A phylogenetic tree was constructed using highly conserved single-copy BUSCO genes via the BUSCO Phylogenomics workflow (https://github.com/jamiemcg/BUSCO_phylogenomics). The genome of closely related *E. globulus* was included as an outgroup ^67^. BUSCO v5.8.2 was used to identify the single-copy genes and evaluate completeness of the assemblies, using the eudicotyledons odb12 dataset ^64^. Protein sequences present in all genomes were aligned with Muscle5 v5.1 ^68^, and poorly aligned regions trimmed using trimAl v1.4.rev15 ^69^. A total of 2,710 complete and single-copy BUSCO sequences in ≥4 genomes remained (97% of the 2,805 BUSCOs). Maximum likelihood gene trees were inferred with IQ-TREE v2.3.6 ^70^, and the species tree estimated under the multi-species coalescent model using ASTRAL-IV v1.23.4.6 ^71^. The tree was generated and manually rooted on *E. globulus* (not displayed) using Figtree v1.4.5 (http://tree.bio.ed.ac.uk/software/figtree/).

To assess synteny and identify structural variations between genomes, pairwise alignments were performed using the MUMmer4 v4.0.1 DNA alignment tool nucmer (--maxmatch -l 50 -b 500 -c 200) ^72^. Alignments were filtered to retain only those ≥200 bp in length and ≥95% sequence identity using the MUMmer4 delta-filter tool. Synteny and structural variants were identified with SyRI v1.6.3 ^73^, and visualised with plotsr v1.1.1 ^74^. Across the distributed provinces of these assembled individuals, some levels of non-synteny were found but the geography did not seem to underpin any genetic isolations.

### Direct RNA sequencing and genome annotation

Total RNA was extracted from the cryogenically frozen leaf tissue of the *Eucalyptus viminalis* reference tree (described above), using a cetrimonium bromide lysis buffer and lithium chloride to precipitate RNA, described in ^75^. The RNA was quality checked on a NanoDrop 1000 Spectrophotometer (Thermo Fisher), a TapeStation 4200 automated electrophoresis instrument with a RNA ScreenTape (Agilent Technologies, 5067-5576), and a Qubit 4 Fluorometer with a Qubit RNA Integrity and Quality (IQ) assay (Thermo Fisher, Q33221), following the manufacturer’s instructions. Poly(A) transcripts were then purified with Oligo (dT)_25_ magnetic beads (NEB S1419S), to enrich intact mRNA. An ONT library direct RNA sequencing library was prepared (SQK-RNA004), using Induro Reverse Transcriptase (NEB M0681) at 60°C for 15 min, to create stable full-length RNA-cDNA molecules (to enhance length and output of the RNA that is sequenced). Direct RNA sequencing was performed on the ONT PromethION P24 platform using a FLO-PRO004RA flow cell. Only the RNA strand of the RNA-cDNA hybrid is sequenced. When sequencing declined, the flow cell was treated once with ONT Flow Cell Wash Kit (EXP-WSH004), using 5 µL DNAse I (NEB M0303) and 5 µL RNase A (Thermo Fisher 12091039). The flow cell was then re-primed and reloaded with more libraries following the manufacturer’s instructions. Sequencing data (pod5) was basecalled to fastq with ONT Dorado v7.6.8, using the super accurate model (sup v5.0.0, 130 bps). Sequence reads were filtered for read lengths ≥ 200 bp and mean read quality ≥ Q10 using Chopper v0.10.0 ^56^. Quality checks were performed before and after filtering with NanoStat 1.6.0 ^76^.

Before annotating genes, both haplotypes were repeat-masked to aid in gene prediction. First, *de novo* repeat libraries were created for each haplotype using EDTA (version: 1.9.6) ^77^, and subsequently repeat annotation was performed with RepeatMasker (version: 4.0.9; https://www.repeatmasker.org). Using these repeat annotations, both haplotypes were soft-masked. Initial transcript models were constructed using filtered long-read RNA data and IsoQuant (version 3.3.1; parameters: --report_novel_unspliced true --stranded forward --model_construction_strategy sensitive_ont ^78^. Based on these RNA-predicted transcripts, gene annotation was performed using BRAKER3 (version 3.0.8) ^79^.

### Species-specific basecaller training and basecalling

Training a species-specific basecaller improves the accuracy of ONT sequencing reads as well as the accuracy of their Q-scores, particularly in plant species (Ferguson et al., 2022). As such, a basecaller for *E. viminalis* was trained for population analyses. The *E. viminalis* basecaller model was trained using the ONT assembly reads and our combinatorial PacBio HiFi, ONT and HiC genome as the truth set. To aid in distinguishing true polymorphisms from sequencing errors, our two haplotypes were joined, creating a diploid truth set.

As ONT sequencing chemistry advanced, we had both 5 kHz and 4 kHz ONT data, accordingly two basecallers were trained, one for each data type. All pod5 reads were randomly sampled to capture variable read lengths. For the 4 kHz model, Bonito was initially used to basecall three read batches of 220 fast5 files (model: dna_r10.4.1_e8.2_400bps_hac@v4.1.0; version: 0.7.2; https://github.com/nanoporetech/bonito). Next, each read subset was used to iteratively train a basecaller model using Bonito with the following parameters: --epochs 16 --batch 160 --chunks 0 --lr 15e-5 --save-optim-every 2 --valid-chunks 112. The 5 kHz model was trained similarly, basecalling with three read subsets of 148 fast5 files using model dna_r10.4.1_e8.2_400bps_hac@v4.2.0 and the same Bonito parameters, with the exception of --epochs 8 and --batch 170. Different training parameters were used because 5 kHz reads contained more data and had a different read length distribution.

Raw population sequencing files in pod5 file format were basecalled with the *E. viminalis* basecaller model and demultiplexed with Dorado (version 0.5.0, linux-x64) (https://github.com/nanoporetech/dorado). The output read FASTQ files were filtered to remove fragmented and low-quality reads using Fastq-filter (version 0.3.0) (https://github.com/LUMC/fastq-filter) (-q 7 -l 1000). Sequencing results were summarised in Supplementary table 7.

### Pangenome analyses

In order to compare sequences and genome synteny across multiple haplotype-resolved genomes, T2T genomes described above and several high-quality genome assemblies of environmental samples (described in later sections) were annotated with Liftoff (version 1.6.3) ^80^ using the reference genome annotations. The annotations of each haplotype genomes were then grouped into orthogroups with OrthoFinder (version 2.5.5) ^81^ based on peptide sequence similarity and the DIAMOND database ^82^. Then, using the orthogroups as anchors across the genome, gene-centric synteny was analysed with GENESPACE (version 1.2.3) (https://github.com/jtlovell/GENESPACE). Additionally, saturations of novel orthogroups with the number of included genomes were also visualised.

### Population sampling

Leaf tissues from 71 wild *E. viminalis* individuals were collected across the natural range of this species from New South Wales, Australia, between 2019 and 2021 for ONT long-read sequencing. These 71 were a subset of a total of 434 samples of *E. viminalis* and two closely related species - *E. dalrympleana* and *E. rubida* - collected for short-read sequencing. Samples derived from three geographically separated populations (populations A-C) delineated by latitude breakpoints of -34.42 and -31.80. Notably, population A is located near the Monaro region where a large-scale dieback event recently occurred ^83^. The collected leaf samples were silica-dried and stored in air-tight containers for long-term storage. All samples collected in this study originated from woodlands as characterised by their rainfall and temperature conditions.

### High-molecular weight DNA extractions of landscape leaf samples

To extract high-molecular weight DNA from the silica-dried leaf samples, a custom protocol was followed. Approximately 40 x 3 mm leaf Uni-Core punches (Qiagen, WB100078) and three 1/8” (3.175 mm) ball bearings were added to a 2 mL microcentrifuge tube for each sample. Samples were frozen under liquid nitrogen and homogenised using a TissueLyser II (Qiagen). The DNA was then extracted with a magnetic bead-based protocol, described in ^54^. In brief, the homogenised tissues were washed with a 0.35 M D-sorbitol solution (DTT freshly added when use), lysed with sodium dodecyl sulphate (SDS) based lysis buffer, washed with Chloroform: Isoamyl alcohol (24:1), and the DNA was bound to Sera-Mag SpeedBead Carboxylate-Modified beads (Cytiva 65152105050250) in the presence of binding buffer (20% PEG 8,000 + 3 M NaCl). A NaCl-based Long fragment buffer (LFB) wash step was introduced (0.75M NaCl + 5% PEG 8,000) to remove small DNA fragments (selecting for ∼3 kb and above). This greatly aided barcoding efficiency and read lengths.

The HMW DNA was eluted from beads with 50 μL of 10 mM Tris-HCl pH 8. The extracted HMW DNA was quantified with Qubit Broad Range Assay Kits (Thermo Fisher Scientific) and their purities were evaluated with the NanoDrop spectrophotometer (Thermo Fisher Scientific). The HMW DNA was stored at 4°C until sequencing. Quality check of DNA was performed on six representative samples, using a Femto Pulse System (Agilent Technologies) in gDNA mode.

### Long-read library preparation and sequencing

Barcoding was performed with the Oxford Nanopore Technologies (ONT) SQK-NBD114.96 according to the manufacturer’s instructions. In brief, 2 μg DNA was aliquoted for each sample and added Ultra II End-prep enzyme and FFPE DNA Repair enzyme (New England BioLabs). The incubation of the repair step was doubled due to the DNA being highly damaged from the silica drying process and storage condition. A barcode was added to each sample (2.5 μL), after which every 12 samples were pooled to be sequenced in batches.The pooled samples were size-selected with BluePippin (Sage Science) for fragments 8 kb and above. An ONT adapter was ligated to the barcoded pool and DNA sequencing was performed on the PromethION platform using a FLO-PRO114M R10.4.1 flow cell following the manufacturer’s instructions. Each flow cell was flushed three times (or until the pore viability had diminished), where more sequencing libraries were added.

To assemble genomes of *E. viminalis* sourced across the landscape, ONT long-reads sequenced with the R10.4.1 chemistry was passed to Hifiasm (version 0.24.0-r702) ^34^. Hifiasm 0.24.0-r702 is able to take the raw reads from ONT sequenced with R10 chemistry or later (--ont --hom-cov [0.8 x estimated coverage] --telo-m AAACCCT --dual-scaf). Hifiasm then performs error corrections in adaptor and homopolymer regions as well as base corrections according to ONT sequencing error profiles. The completeness of the assembled contigs were then assessed with Compleasm (version 0.2.6) ^63^ and reported in Supplementary table 8.

### Structural variant discovery and genotyping

The filtered long-read sequences from each sample were mapped to the reference genome (haplotype 1) with Minimap2 (version 2.24) ^84^ to generate Binary Alignment Map (BAM) files for all 71 sequenced samples. The output BAM files are then sorted and indexed with SAMtools (version 1.9) ^85^. After the appropriate level of genome-wide coverage was determined for both SV discovery and genotyping (Supplementary Note 8), 34 samples with genome-wide coverage >12x were included for SV discovery with Sniffles2 (version 2.0.7) ^86^ (n = 34; population A: 18, B: 1, C: 15), and 49 samples were included in the population genetic studies of SV genotype allele frequency as well as subsequent GEA analyses.

SVs were merged based on breakpoint locations, percentage overlaps (relative to SV size), and in some cases, sequence similarities. Testing different SV merging criteria revealed that the number of mergeable SVs monotonically decreases as merging criteria becomes increasingly stringent (Supplemental Note 8). To retain SV diversity without inflating allele frequencies of certain SVs, intermediate levels of merging criteria were chosen. For the majority of SVs, a threshold of 0.2 (20%) maximum linear distance (in relation to SV size) between breakpoint positions was chosen. For insertions > 1 kb (SVTYPE == ‘INS’, SVLEN >= 1000), a sequence similarity of 0.75 (75%) were added in addition to the maximum distance criterion to separate large insertions with similar breakpoint positions, but have different sequence content novel to the reference genome. Merged SVs were genotyped with CuteSV (version 2.0.3; -Ivcf) ^87^ in all 71 sequenced individuals in order to remove singletons while retaining maximum number of SVs, resulting. Singleton SVs were removed with BCFtools (version 1.12) ^85^ by variant ID (-m id). Then, variants found in fewer than 2 individuals (i.e., singletons) were filtered out, resulting in 573,889 long-read discovered structural variants that are 1) found by both Sniffles2 and CuteSV, and 2) found in at least two biological samples.

### Associating SVs with environmental variables

Prior to performing a genomic-environment association (GEA) study with structural variants, SVs were filtered at high stringency to ensure quality. First, only SVs supported by a minimum of three reads during SV genotyping were maintained: the rest were treated as missing data. Second, SV loci with more than 10% missing data were filtered out. Last, rare SVs (minor allele frequency < 0.02) were filtered out. In order to keep the sample individuals comparable, only the 49 trees with > 8x genome-wide coverage were used in the GEA.

Tests for associations between SV polymorphisms and environmental variables were implemented in GEMMA (version 0.98.3) ^88^. The sample relatedness matrix (or genomic relation matrix, GRM) was calculated using genome-wide SV genotypes (option -gk). For each of the 49 tree locations, long-term climate variables at 2.5 minutes spatial resolution were extracted from the BioClim data base ^89^ using the R package geospatial (https://github.com/rspatial/geodata). Then, a linear mixed effect model (-lmm) was fitted by using SV allele frequencies to predict BioClim variables using the GRM information as the random effect in order to account for neutral variation in allele frequencies. After the GEA was performed, the Bonferroni threshold (p-value < 10^-12^) was applied to identify putatively adaptive loci.

### Short-read validation of climate adaptive loci

Leaf tissue was collected from 176 *E. viminalis*, 138 *E. dalrympleana*, and 120 *E. rubida* (total of 434) samples (metadata in Supplementary Table 16). These included the 49 samples that were long-read sequenced above. Leaves were added to envelopes and dried on silica gel for long-term storage. Approximately twenty 3 mm leaf punches (Qiagen Uni-Core, WB100078) and one 3 mm ball bearing per sample were added to 96-well arrayed 1.1 mL tubes (Axygen Scientific, MTS-11-8-C-R-10 and MTS-8CP-C-1). Samples were frozen in liquid nitrogen and homogenized using TissueLyser II (Qiagen). DNA was extracted with a NucleoSpin 96 Plant II 24×96 kit using CTAB lysis buffer PL1 according to the manufacturer’s instructions (Macherey-Nagel, 740663.24).

Whole-genome short-read sequencing libraries were prepared using our cost-optimized transposase protocol ^90^, based on Illumina DNA Prep (M) Tagmentation (24 Samples, IPB) (Cat# 20060060) with bead-linked transposase BLT (Illumina document# 1000000033561 v05). Fragmented DNA was amplified using custom index primers with 14 PCR cycles. Libraries were pooled and size-selected for 300-500 bp inserts using PippinHT (Sage Science). Sequencing was performed on NovaSeq 6000 (Illumina) using an S4 flow cell with a 300-cycle kit (150 bp paired-end) at the Biomolecular Resource Facility, Australian National University, Canberra, Australia.

Variants were called using Acanthophis ^91^, using AdapterRemoval2 ^92^, BWA MEM ^93^, bcftools ^94^, DeepVariant ^95^, GLNexus ^96^, and Snakemake ^97^ against the reference genome. Variants were filtered with SNPSnip (https://pypi.org/project/snpsnip/), VCFtools ^98^ and R scripts. SNPs were filtered out if they had a quality score of <24, a minor allele frequency of <0.10 and a missingness rate of >0.7 across the 434 samples.

For the 49 samples that had both long-read and short-read data, correlations between SV-based *CHILL1* genotype and SNP genotypes at individual sites within the 27.0 to 27.9 Mb wider *CHILL1* region were computed in order to discover potentially diagnostic SNPs for *CHILL1*. Principal components (PCs) based on SNPs within the region containing 5 adjacent SVs (positions 27.8 to 27.8 Mb) and where the highest correlations occurred, were computed using the pcaMethods package in R (Stacklies et al., 2007) to determine the threshold values of the first PC that reliably distinguished *CHILL1* genotypes (Supplementary Note 13).

To explore haplotype structure in the region surrounding *CHILL1*, genotypes from SNPs at a minimum spacing of 100bp within a 0.9Mb segment surrounding the locus (positions 27.0-27.9Mb) were visualised using the pheatmap package in R (https://github.com/raivokolde/pheatmap) and clustered using the hclust() function (option ‘method - Manhattan’) from the stats package in R. Genetic similarities among samples were further visualised by computing genetic distances using the dist() function from the stats package in R (https://stat.ethz.ch/R-manual/R-devel/library/stats/html/00Index.html).

