## Supplemental Tables for "Complex Genomic Structural Variation Underlies Climate Adaptation across *Eucalyptus* species"

### Supplementary Tables

**Supplementary Table 1 Sequencing reads for *Eucalyptus viminalis* ACT, VIC and TAS.** All PacBio HiFi reads, filtered ONT ultra-long reads (≥55 kb and ≥Q20) and filtered Hi-C reads (described in next table) were used in the genome assembly of *E. viminalis* ACT. Only filtered ONT reads (≥20 kb and ≥Q20) were used in the genome assembly of *E. viminalis* VIC and TAS. Therefore, these reads were filtered more stringently, but also benefited from updated sequencing chemistry and assembly tools.

| | **Evim ACT** | | | **Evim VIC** | | **Evim TAS** | |
|---|---|---|---|---|---|---|---|
| **Metric** | **HiFi** | **ONT** | **ONT ≥55 kb ≥Q7** | **ONT** | **ONT ≥20 kb ≥Q20** | **ONT** | **ONT ≥20 kb ≥Q20** |
| Reads | 1,493,183 | 6,802,892 | 471,525 | 3,119,232 | 1,182,139 | 2,684,427 | 894,493 |
| Yield (bp) | 30,268,834,753 | 176,995,195,958 | 32,674,908,869 | 85,888,723,721 | 53,511,358,482 | 66,828,255,396 | 40,656,940,200 |
| Mean read length (bp) | 20,271.3 | 26,017.6 | 69,296.2 | 27,535.2 | 45,266.6 | 24,894.8 | 45,452.5 |
| Median read length (bp) | 19,515 | 23,524 | 64,736 | 28,391 | 42,292 | 19,822 | 42,314 |
| N50 (bp) | 20,332 | 36,617 | 66,993 | 46,246 | 47,264 | 46,378 | 47,725 |
| Mean read quality | 29.2 | 16.4 | 17.5 | 9.8 | 22.6 | 9.2 | 22.6 |
| Median read quality | 30.2 | 18.9 | 18.4 | 20.5 | 23.6 | 20.1 | 23.6 |
| Five longest read lengths (and Q score) #1 | 55,114 (15.2) | 463,629 (3.5) | 564,886 (7.5) | 1,875,904 (6.2) | 375,909 (21.3) | 1,410,201 (5.2) | 307,800 (23.5) |
| #2 | 55,332 (24.0) | 469,113 (3.9) | 484,984 (7.3) | 1,843,354 (6.2) | 281,550 (22.9) | 1,385,395 (3.5) | 284,444 (21.6) |
| #3 | 55,411 (15.1) | 484,984 (7.3) | 293,940 (19.6) | 1,554,198 (5.5) | 269,294 (24.3) | 1,218,598 (4.5) | 259,566 (20.1) |
| #4 | 56,218 (17.9) | 564,886 (7.5) | 286,672 (19.9) | 1,310,354 (6.4) | 254,787 (24.0) | 1,191,299 (2.9) | 258,757 (20.8) |
| #5 | 57,165 (15.8) | 806,429 (2.7) | 282,977 (7.8) | 1,271,470 (6.3) | 250,349 (25.0) | 1,123,223 (8.2) | 251,260 (21.3) |
| Coverage (547 Mbp genome) | 55x | 324x | 60x | 157x | 98x | 122x | 74x |
| Coverage per haplotype | 28x | 162x | 30x | 79x | 49x | 61x | 37x |

**Supplementary Table 2. Hi-C sequencing results and chromosomal linkage statistics for the *E. viminalis* ACT reference genome.** Statistics are shown for both haplotypes and were used for chromosome scaffolding and assembly validation.

| | **Haplotype 1** | **Haplotype 2** |
|---|---|---|
| **Hi-C Read Pairs** | 89,043,318 | |
| **Inter-chromosomal** | 10,570,983 | 10,599,530 |
| **Intra-chromosomal** | 19,716,057 | 19,602,535 |
| **Uninformative Read Pairs** | 58,756,278 | 58,841,253 |

**Supplementary Table 3. Genome assembly statistics for *Eucalyptus viminalis* ACT, VIC and TAS.** For each genome, the 11 chromosomes were haplotype resolved, at chromosome scale or higher (many being telomere-to-telomere). The *E. viminalis* ACT genome (first genome) was constructed with PacBio HiFi, ONT ultra-long and Hi-C. The *E. viminalis* VIC and TAS genomes were assembled with ONT ultra-long reads only, but benefited from newer sequencing chemistry and assembly algorithms, having only 1 gap at the chromosome two centromere. Karyotype is 2n = 22.

| Metric | Evim ACT | | Evim VIC | | Evim TAS | |
|---|---|---|---|---|---|---|
| | Hap1 | Hap2 | Hap 1 | Hap 2 | Hap 1 | Hap 2 |
| Sequences | 294 | 287 | 392 | 25 | 298 | 28 |
| Sequences >1 Mbp | 11 | 11 | 11 | 11 | 11 | 11 |
| N50 (bp) | 51,735,356 | 50,558,970 | 43,272,629 | 52,040,736 | 40,964,566 | 52,728,202 |
| N50 (bp) >1 Mbp | 51,735,356 | 50,558,970 | 51,281,642 | 52,040,736 | 50,289,127 | 53,905,933 |
| Size (bp) | 546,531,677 | 551,267,619 | 589,735,305 | 531,995,131 | 559,924,252 | 536,652,899 |
| Size (bp) >1 Mbp | 530,050,200 | 530,973,922 | 538,353,018 | 529,912,749 | 526,300,622 | 533,344,571 |
| Telomeres | 2/22 | 3/22 | 22/22 | 22/22 | 22/22 | 21/22 |
| T2T contigs | 0/11 | 0/11 | 10/11 | 10/11 | 10/11 | 9/11 |
| T2T scaffolds | 0/11 | 0/11 | 11/11 | 11/11 | 11/11 | 10/11 |
| Gaps | 94 | 165 | 1 | 1 | 1 | 1 |

**Supplementary Table 4. BUSCO completeness assessment of *E. viminalis* genome assemblies.** Single-copy orthologs from the eudicotyledons odb12 dataset were identified using compleasm v0.2.7 to evaluate the completeness of the 11 haplotype-resolved chromosomal scaffolds for each genome.

| | Evim ACT | | Evim VIC | | Evim TAS | |
|---|---|---|---|---|---|---|
| BUSCO | Hap 1 | Hap 2 | Hap 1 | Hap 2 | Hap 1 | Hap 2 |
| Complete | 99.61% (2,794) | 99.68% (2,796) | 99.72% (2,797) | 99.72% (2,797) | 99.72% (2,797) | 99.68% (2,796) |
| Single | 98.11 | 97.79% | 97.83% | 98.04% | 98.40% | 97.93% |

| | | | | | | |
|---|---|---|---|---|---|---|
| | (2,752) | (2,743) | (2,744) | (2,750) | (2,760) | (2,747) |
| Duplicated | 1.50% (42) | 1.89% (53) | 1.89% (53) | 1.68% (47) | 1.32% (37) | 1.75% (49) |
| Fragmented s1 | 0.29% (8) | 0.25% (7) | 0.21% (6) | 0.21% (6) | 0.21% (6) | 0.25% (7) |
| Fragmented s2 | 0.00% (0) | 0.00% (0) | 0.00% (0) | 0.00% (0) | 0.00% (0) | 0.00% (0) |
| Missing | 0.11% (3) | 0.07% (2) | 0.07% (2) | 0.07% (2) | 0.07% (2) | 0.07% (2) |
| Number genes | 2,805 | 2,805 | 2,805 | 2,805 | 2,805 | 2,805 |

**Supplementary Table 5. Assembly quality assessment using Quality Value (QV) scores for haplotype-resolved *E. viminalis* genomes.** Assembly accuracy was assessed using yak v0.1 (r56) with default k=31 for each chromosome-scale haplotype (Cheng et al 2021 https://www.nature.com/articles/s41592-020-01056-5). Raw and coverage adjusted QV scores are shown as Phred-scaled values and accuracy percentages.

| | Evim ACT (HiFi) | | Evim VIC (ONT ≥20 kb ≥Q20) | | Evim TAS (ONT ≥20 kb ≥Q20) | |
|---|---|---|---|---|---|---|
| | Hap 1 | Hap 2 | Hap 1 | Hap 2 | Hap 1 | Hap 2 |
| Raw QV | 56.415 | 55.525 | 66.772 | 67.085 | 64.368 | 64.116 |
| Raw QV % | 99.999772 | 99.999720 | 99.999979 | 99.999980 | 99.999963 | 99.999962 |
| Adjusted QV | 57.591 | 55.845 | 64.562 | 64.620 | 60.366 | 60.395 |
| Adjusted QV % | 99.999826 | 99.999740 | 99.999965 | 99.999965 | 99.999910 | 99.999911 |

**Supplementary Table 6. ONT direct RNA sequencing results for an *E. viminalis* ACT leaf sample.** Reads were filtered to retain only those ≥200 bp in length and ≥Q10 quality score for the annotation of the genome and identification of isoforms.

| **Metric** | **Raw RNA reads** | **Filtered RNA reads** |
|---|---|---|
| Mean Read Length (bp) | 1,134.2 | 1,303.4 |
| Mean Quality | 12.2 | 19.2 |
| Median Read Length (bp) | 937.0 | 1,092.0 |
| Median Quality | 20.9 | 21.6 |
| Reads | 22,125,880 | 18,024,536 |
| N50 | 1,605.0 | 1,618.0 |
| STDEV read length | 1,130.1 | 877.9 |
| Yield (bp) | 25,094,940,983 | 23,493,482,035 |
| Five longest reads (and Q score) #1 | 241,297 (5.7) | 27,417 (22.8) |
| #2 | 224,525 (3.9) | 25,793 (22.0) |
| #3 | 210,414 (4.2) | 21,167 (21.4) |
| #4 | 208,103 (6.7) | 20,781 (22.2) |
| #5 | 205,564 (4.0) | 20,457 (23.4) |

**Supplementary Table 7. Sequencing summary from Oxford Nanopore Technologies (ONT) long-read sequencing of environmental samples (n = 72).** after quality (average Q score > 7) and read size (minimum read length > 1 kb) filtering.

| | sample_name | coverage | num_reads | read_length_N50 | total_bases_Gb | max_read_length | median_read_length |
|---|---|---|---|---|---|---|---|
| 2 | NSW0083 | 8.95098 | 962129 | 6.156 | 4.70225 | 56.823 | 4.745 |
| 3 | NSW0105 | 11.4163 | 1486687 | 5.395 | 5.99737 | 61.321 | 3.132 |
| 4 | NSW0108 | 5.89502 | 685649 | 5.833 | 3.09686 | 43.319 | 4.271 |
| 5 | NSW0110 | 2.04307 | 225281 | 6.623 | 1.0733 | 44.32 | 3.978 |
| 6 | NSW0111 | 7.71839 | 992345 | 5.721 | 4.05473 | 42.2 | 3.167 |
| 7 | NSW0113 | 11.3181 | 950043 | 8.076 | 5.9458 | 66.902 | 5.839 |
| 8 | NSW0119 | 7.69559 | 864114 | 5.994 | 4.04276 | 47.049 | 4.427 |
| 9 | NSW0120 | 3.36433 | 429528 | 5.481 | 1.7674 | 79.467 | 3.651 |
| 10 | NSW0133 | 1.00533 | 154295 | 4.417 | 0.528136 | 42.071 | 2.562 |
| 11 | NSW0141 | 18.6357 | 1586697 | 7.729 | 9.78998 | 84.917 | 5.53 |
| 12 | NSW0144 | 13.3328 | 1213005 | 7.767 | 7.00416 | 74.695 | 4.778 |
| 13 | NSW0159 | 12.5647 | 884301 | 10.233 | 6.60067 | 84.868 | 6.54 |
| 14 | NSW0163 | 13.6504 | 1025529 | 9.094 | 7.17101 | 73.836 | 6.453 |
| 15 | NSW0170 | 17.7472 | 1428360 | 8.257 | 9.32323 | 92.544 | 5.734 |
| 16 | NSW0175 | 9.84936 | 829086 | 8.335 | 5.1742 | 98.098 | 5.216 |
| 17 | NSW0192 | 3.55784 | 437749 | 5.807 | 1.86906 | 47.266 | 3.498 |
| 18 | NSW0194 | 4.04678 | 472992 | 6.177 | 2.12591 | 121.675 | 3.702 |
| 19 | NSW0207 | 11.1162 | 1514158 | 5.189 | 5.8397 | 89.842 | 3.055 |
| 20 | NSW0209 | 2.39105 | 301305 | 5.776 | 1.2561 | 105.191 | 3.288 |
| 21 | NSW0250 | 16.8051 | 1240465 | 9.404 | 8.82829 | 87.516 | 6.374 |
| 22 | NSW0293 | 29.9479 | 1556811 | 13.1 | 15.7326 | 156.083 | 9.485 |
| 23 | NSW0296 | 13.7463 | 1044468 | 8.396 | 7.22143 | 73.649 | 6.264 |
| 24 | NSW0299 | 9.98593 | 1434379 | 4.918 | 5.24595 | 65.534 | 2.84 |
| 25 | NSW0306 | 24.4949 | 2717245 | 6.317 | 12.868 | 66.36 | 3.937 |
| 26 | NSW0308 | 19.8459 | 2221206 | 6.244 | 10.4257 | 343.493 | 3.921 |
| 27 | NSW0313 | 18.9491 | 1681010 | 7.596 | 9.95462 | 81.822 | 5.156 |
| 28 | NSW0315 | 6.49355 | 544534 | 8.263 | 3.41129 | 76.15 | 5.524 |
| 29 | NSW0323 | 9.19755 | 886616 | 6.575 | 4.83179 | 67.277 | 5.444 |
| 30 | NSW0324 | 14.9675 | 1095822 | 9.919 | 7.86292 | 82.278 | 6.192 |
| 31 | NSW0328 | 11.1906 | 775432 | 10.455 | 5.8788 | 105.051 | 6.5255 |
| 32 | NSW0334 | 12.7057 | 907636 | 10.161 | 6.67474 | 90.242 | 6.347 |
| 33 | NSW0335 | 10.751 | 806298 | 9.894 | 5.64788 | 71.769 | 5.852 |

| | | | | | | | |
|---|---|---|---|---|---|---|---|
| 34 | NSW0337 | 16.0358 | 1313350 | 7.708 | 8.42416 | 63.244 | 5.994 |
| 35 | NSW0341 | 13.883 | 1202451 | 7.381 | 7.29322 | 77.341 | 5.73 |
| 36 | NSW0343 | 11.1506 | 863310 | 9.66 | 5.85778 | 74.817 | 5.524 |
| 37 | NSW0347 | 9.58598 | 758783 | 9.411 | 5.03584 | 73.146 | 5.453 |
| 38 | NSW0348 | 10.1471 | 781891 | 9.652 | 5.33063 | 86.062 | 5.603 |
| 39 | NSW0350 | 15.6948 | 1651628 | 6.49 | 8.24502 | 61.998 | 4.421 |
| 40 | NSW0354 | 14.2373 | 1517831 | 6.588 | 7.47936 | 65.054 | 4.12 |
| 41 | NSW0356 | 13.5844 | 1428460 | 6.751 | 7.13637 | 59.6 | 3.999 |
| 42 | NSW0357 | 13.7632 | 1045792 | 8.816 | 7.23029 | 58.483 | 6.504 |
| 43 | NSW0359 | 16.8905 | 1608212 | 7.258 | 8.87317 | 79.355 | 4.755 |
| 44 | NSW0360 | 0.177873 | 21904 | 5.78 | 0.0934427 | 39.709 | 3.305 |
| 45 | NSW0366 | 21.2167 | 1967794 | 7 | 11.1459 | 102.709 | 5.291 |
| 46 | NSW0438 | 23.6787 | 1369640 | 11.573 | 12.4392 | 105.884 | 8.143 |
| 47 | NSW0482 | 0.643269 | 91807 | 4.935 | 0.337931 | 32.976 | 2.775 |
| 48 | NSW0491 | 17.7734 | 2328606 | 5.465 | 9.33695 | 52.861 | 3.165 |
| 49 | NSW0493 | 14.8659 | 1202607 | 8.905 | 7.80955 | 55.711 | 5.677 |
| 50 | NSW0496 | 20.0196 | 2301955 | 6.008 | 10.517 | 84.553 | 3.886 |
| 51 | NSW0507 | 0.891656 | 123623 | 5.077 | 0.468417 | 110.646 | 2.835 |
| 52 | NSW0532 | 17.8178 | 1163923 | 10.001 | 9.36032 | 90.239 | 7.416 |
| 53 | NSW0536 | 12.7913 | 1175455 | 7.649 | 6.71972 | 77.711 | 4.936 |
| 54 | NSW0544 | 18.8856 | 1480507 | 8.65 | 9.92127 | 90.58 | 6.162 |
| 55 | NSW0555 | 11.2373 | 1646519 | 4.768 | 5.90335 | 206.865 | 2.805 |
| 56 | NSW0561 | 13.5694 | 1104498 | 8.972 | 7.12846 | 66.231 | 5.372 |
| 57 | NSW0566 | 12.2311 | 1820740 | 4.64 | 6.42541 | 160.07 | 2.794 |
| 58 | NSW0573 | 12.383 | 1622781 | 5.407 | 6.50519 | 118.511 | 3.243 |
| 59 | NSW0575 | 8.35824 | 850173 | 6.382 | 4.39087 | 64.073 | 5.01 |
| 60 | NSW0576 | 20.1635 | 1455137 | 10.067 | 10.5926 | 100.658 | 6.28 |
| 61 | NSW0934 | 17.2554 | 1979918 | 6.175 | 9.06485 | 61.207 | 3.759 |
| 62 | NSW0935 | 2.77763 | 382010 | 5.059 | 1.45918 | 56.067 | 2.916 |
| 63 | NSW0936 | 1.81097 | 251023 | 5.055 | 0.951362 | 55.849 | 2.801 |
| 64 | NSW0938 | 2.33443 | 313364 | 5.242 | 1.22636 | 68.637 | 2.966 |
| 65 | NSW0939 | 4.3979 | 573694 | 5.349 | 2.31037 | 58.417 | 3.116 |
| 66 | NSW0940 | 3.20183 | 425685 | 5.346 | 1.68203 | 105.466 | 3.278 |
| 67 | NSW0954 | 6.24472 | 492787 | 8.714 | 3.28057 | 64.006 | 6.035 |
| 68 | NSW0962 | 19.2845 | 1780998 | 7.04 | 10.1308 | 104.905 | 5.262 |
| 69 | NSW0963 | 4.04471 | 491526 | 5.778 | 2.12483 | 45.066 | 3.639 |
| 70 | NSW0966 | 6.39348 | 672144 | 6.409 | 3.35871 | 97.581 | 4.335 |

| | | | | | | | |
|---|---|---|---|---|---|---|---|
| 71 | NSW0970 | 10.1207 | 857272 | 8.3 | 5.31674 | 94.322 | 5.231 |
| 72 | NSW0972 | 11.1123 | 936771 | 8.229 | 5.83765 | 80.982 | 5.496 |

**Supplementary Table 8. Haplotype-resolved assembly quality of environmental samples (n = 71).** Genomes assembled with hifiasm (--ont). Table ordered by BUSCO scores, highest to lowest.

| sample_name | BUSCO | population | coverage | haplotype | mean_length | median_length | contigs | n50_length | stdev_length | total_bases |
|---|---|---|---|---|---|---|---|---|---|---|
| NSW0293 | 98.79 | C | 29.9479 | hap2 | 1,547,592 | 1,036,918 | 343 | 2,588,597 | 1,521,859 | 530,824,011 |
| NSW0293 | 98.45 | C | 29.9479 | hap1 | 1,103,047 | 658,789 | 503 | 2,037,153 | 1,278,829 | 554,832,757 |
| NSW0438 | 95.18 | B | 23.6787 | hap1 | 151,300 | 95,764 | 3,758 | 255,713 | 162,315 | 568,583,354 |
| NSW0438 | 93.9 | B | 23.6787 | hap2 | 184,573 | 131,570 | 2,727 | 280,315 | 175,737 | 503,330,782 |
| NSW0306 | 92.73 | C | 24.4949 | hap1 | 67,245 | 43,741 | 8,176 | 106,540 | 80,182 | 549,792,405 |
| NSW0306 | 91.23 | C | 24.4949 | hap2 | 75,405 | 50,902 | 6,516 | 113,726 | 84,158 | 491,339,593 |
| NSW0576 | 93.64 | A | 20.1635 | hap1 | 109,783 | 74,990 | 4,996 | 170,727 | 110,774 | 548,475,552 |
| NSW0576 | 90.15 | A | 20.1635 | hap2 | 121,546 | 87,360 | 4,018 | 176,608 | 112,517 | 488,372,190 |
| NSW0544 | 88.74 | A | 18.8856 | hap1 | 62,473 | 44,269 | 8,418 | 86,132 | 69,306 | 525,899,167 |
| NSW0496 | 86.98 | A | 20.0196 | hap1 | 47,287 | 32,496 | 11,132 | 69,946 | 47,864 | 526,402,658 |
| NSW0496 | 85.04 | A | 20.0196 | hap2 | 52,953 | 37,983 | 8,709 | 75,038 | 51,707 | 461,170,727 |
| NSW0544 | 84.73 | A | 18.8856 | hap2 | 70,038 | 52,252 | 6,428 | 93,761 | 68,081 | 450,202,316 |
| NSW0313 | 85.43 | C | 18.9491 | hap1 | 50,778 | 37,686 | 10,318 | 69,271 | 46,491 | 523,922,927 |
| NSW0141 | 84.44 | A | 18.6357 | hap1 | 51,439 | 38,021 | 9,645 | 70,661 | 45,653 | 496,128,445 |
| NSW0313 | 82.93 | C | 18.9491 | hap2 | 52,157 | 39,787 | 8,186 | 69,384 | 44,181 | 426,959,842 |
| NSW0962 | 83.71 | A | 19.2845 | hap1 | 45,806 | 33,925 | 11,199 | 62,089 | 40,252 | 512,979,448 |
| NSW0532 | 83.79 | A | 17.8178 | hap1 | 59,209 | 45,638 | 8,512 | 76,829 | 46,281 | 503,987,373 |
| NSW0532 | 81.99 | A | 17.8178 | hap2 | 69,239 | 54,565 | 6,472 | 90,321 | 51,601 | 448,114,270 |
| NSW0250 | 85.34 | A | 16.8051 | hap1 | 59,296 | 44,576 | 8,721 | 79,777 | 55,503 | 517,118,994 |
| NSW0141 | 80.56 | A | 18.6357 | hap2 | 57,661 | 43,917 | 7,491 | 77,501 | 48,296 | 431,935,882 |
| NSW0308 | 81.17 | C | 19.8459 | hap1 | 36,601 | 26,961 | 13,107 | 49,317 | 36,583 | 479,725,070 |
| NSW0170 | 81.6 | A | 17.7472 | hap1 | 47,744 | 36,535 | 10,120 | 62,879 | 38,601 | 483,167,736 |
| NSW0962 | 79.63 | A | 19.2845 | hap2 | 47,885 | 36,780 | 8,806 | 63,026 | 39,211 | 421,673,339 |
| NSW0324 | 81 | C | 14.9675 | hap1 | 48,921 | 38,543 | 9,700 | 61,626 | 38,011 | 474,530,096 |
| NSW0366 | 79.32 | C | 21.2167 | hap1 | 37,129 | 27,810 | 12,808 | 48,089 | 33,381 | 475,543,718 |
| NSW0250 | 78.42 | A | 16.8051 | hap2 | 63,539 | 49,071 | 6,428 | 83,853 | 50,076 | 408,425,847 |
| NSW0170 | 77.73 | A | 17.7472 | hap2 | 55,849 | 44,249 | 7,543 | 72,053 | 46,242 | 421,268,467 |
| NSW0359 | 77.08 | C | 16.8905 | hap1 | 33,615 | 26,849 | 13,125 | 41,914 | 24,960 | 441,196,281 |
| NSW0324 | 75.15 | C | 14.9675 | hap2 | 53,583 | 42,312 | 6,934 | 66,843 | 42,523 | 371,544,227 |

| | | | | | | | | | | |
|---|---|---|---|---|---|---|---|---|---|---|
| NSW0493 | 76.27 | A | 14.8659 | hap1 | 42,726 | 33,208 | 10,838 | 52,725 | 40,837 | 463,068,729 |
| NSW0308 | 73.99 | C | 19.8459 | hap2 | 38,652 | 29,756 | 9,969 | 49,974 | 34,279 | 385,323,715 |
| NSW0337 | 77.04 | C | 16.0358 | hap1 | 37,298 | 29,626 | 12,406 | 45,914 | 27,221 | 462,714,827 |
| NSW0934 | 76.35 | A | 17.2554 | hap1 | 32,211 | 24,209 | 14,508 | 43,082 | 40,785 | 467,323,567 |
| NSW0350 | 75.67 | C | 15.6948 | hap1 | 31,941 | 24,022 | 14,059 | 40,295 | 33,279 | 449,051,886 |
| NSW0366 | 74.46 | C | 21.2167 | hap2 | 41,448 | 31,864 | 9,612 | 52,411 | 46,169 | 398,394,630 |
| NSW0491 | 73.69 | A | 17.7734 | hap1 | 26,088 | 19,805 | 16,265 | 33,807 | 26,558 | 424,318,531 |
| NSW0337 | 72.39 | C | 16.0358 | hap2 | 44,662 | 35,637 | 8,661 | 55,664 | 34,082 | 386,812,955 |
| NSW0357 | 72.36 | C | 13.7632 | hap1 | 36,389 | 29,986 | 11,831 | 43,592 | 24,032 | 430,515,197 |
| NSW0296 | 71.03 | C | 13.7463 | hap1 | 34,693 | 28,372 | 11,964 | 41,449 | 27,297 | 415,066,115 |
| NSW0356 | 69.05 | C | 13.5844 | hap1 | 26,038 | 21,124 | 14,579 | 31,617 | 19,647 | 379,613,800 |
| NSW0561 | 71.02 | A | 13.5694 | hap1 | 36,892 | 30,253 | 11,306 | 45,057 | 25,756 | 417,099,249 |
| NSW0934 | 69.35 | A | 17.2554 | hap2 | 33,799 | 26,285 | 10,835 | 42,864 | 38,797 | 366,214,720 |
| NSW0359 | 68.44 | C | 16.8905 | hap2 | 39,196 | 31,340 | 9,010 | 49,461 | 30,879 | 353,157,770 |
| NSW0163 | 70.68 | A | 13.6504 | hap1 | 37,948 | 31,294 | 11,167 | 45,785 | 25,790 | 423,762,578 |
| NSW0341 | 70.55 | C | 13.883 | hap1 | 31,540 | 25,543 | 13,042 | 38,004 | 23,224 | 411,339,821 |
| NSW0491 | 68.15 | A | 17.7734 | hap2 | 31,553 | 23,989 | 11,359 | 40,929 | 32,769 | 358,408,570 |
| NSW0159 | 71.41 | A | 12.5647 | hap1 | 40,826 | 33,108 | 9,976 | 49,117 | 28,723 | 407,276,499 |
| NSW0334 | 69.39 | C | 12.7057 | hap1 | 38,525 | 32,167 | 10,696 | 46,056 | 26,053 | 412,068,205 |
| NSW0536 | 67.67 | A | 12.7913 | hap1 | 29,309 | 24,168 | 13,056 | 34,831 | 21,233 | 382,652,192 |
| NSW0493 | 67.15 | A | 14.8659 | hap2 | 46,060 | 36,519 | 7,733 | 56,505 | 41,002 | 356,179,336 |
| NSW0354 | 66.13 | C | 14.2373 | hap1 | 24,837 | 20,157 | 14,853 | 29,936 | 18,419 | 368,908,822 |
| NSW0144 | 65.52 | A | 13.3328 | hap1 | 29,448 | 24,281 | 12,461 | 36,041 | 20,534 | 366,954,707 |
| NSW0350 | 64.1 | C | 15.6948 | hap2 | 33,495 | 26,306 | 9,678 | 41,664 | 29,408 | 324,166,737 |
| NSW0113 | 64.75 | B | 11.3181 | hap1 | 29,275 | 24,316 | 12,271 | 33,704 | 21,479 | 359,238,294 |
| NSW0343 | 64.19 | C | 11.1506 | hap1 | 32,968 | 27,677 | 10,813 | 39,119 | 21,501 | 356,477,162 |
| NSW0328 | 63.5 | C | 11.1906 | hap1 | 37,828 | 31,262 | 9,464 | 44,571 | 27,518 | 357,999,517 |
| NSW0972 | 61.74 | A | 11.1123 | hap1 | 29,178 | 24,511 | 11,805 | 34,175 | 19,772 | 344,446,817 |
| NSW0335 | 62.12 | C | 10.751 | hap1 | 33,237 | 28,349 | 10,180 | 38,965 | 20,677 | 338,351,538 |
| NSW0348 | 59.76 | C | 10.1471 | hap1 | 30,964 | 26,286 | 9,699 | 36,252 | 18,850 | 300,317,063 |
| NSW0561 | 58.17 | A | 13.5694 | hap2 | 42,154 | 34,781 | 7,095 | 51,612 | 28,216 | 299,079,802 |
| NSW0573 | 57.61 | A | 12.383 | hap1 | 17,521 | 14,628 | 17,567 | 21,024 | 12,790 | 307,798,328 |
| NSW0970 | 58.86 | A | 10.1207 | hap1 | 27,345 | 23,157 | 10,559 | 32,242 | 17,587 | 288,739,168 |
| NSW0347 | 56.06 | C | 9.58598 | hap1 | 29,475 | 25,165 | 9,350 | 34,616 | 18,034 | 275,591,477 |
| NSW0105 | 55.46 | B | 11.4163 | hap1 | 17,650 | 14,559 | 17,951 | 21,708 | 13,209 | 316,834,955 |
| NSW0357 | 55.5 | C | 13.7632 | hap2 | 42,089 | 34,188 | 7,177 | 51,001 | 28,214 | 302,069,105 |
| NSW0163 | 54.73 | A | 13.6504 | hap2 | 41,844 | 34,414 | 6,988 | 50,526 | 29,291 | 292,403,542 |
| NSW0175 | 54.47 | A | 9.84936 | hap1 | 27,837 | 23,107 | 9,981 | 32,607 | 18,523 | 277,839,838 |

| | | | | | | | | | | |
|---|---|---|---|---|---|---|---|---|---|---|
| NSW0341 | 53.87 | C | 13.883 | hap2 | 33,571 | 27,632 | 7,944 | 40,210 | 22,829 | 266,684,305 |
| NSW0207 | 53.13 | A | 11.1162 | hap1 | 15,593 | 13,197 | 18,349 | 18,658 | 10,552 | 286,122,173 |
| NSW0566 | 51.72 | A | 12.2311 | hap1 | 14,295 | 11,874 | 19,866 | 17,219 | 10,254 | 283,989,607 |
| NSW0296 | 51.46 | C | 13.7463 | hap2 | 38,452 | 31,668 | 7,167 | 45,860 | 30,357 | 275,587,477 |
| NSW0555 | 50.94 | A | 11.2373 | hap1 | 14,123 | 11,668 | 17,706 | 17,025 | 10,983 | 250,060,142 |
| NSW0159 | 51.33 | A | 12.5647 | hap2 | 45,933 | 37,900 | 5,489 | 55,355 | 31,654 | 252,127,784 |
| NSW0334 | 47.89 | C | 12.7057 | hap2 | 41,908 | 35,064 | 5,784 | 49,933 | 28,214 | 242,393,300 |
| NSW0144 | 45.58 | A | 13.3328 | hap2 | 34,396 | 28,716 | 6,565 | 41,939 | 23,445 | 225,809,010 |
| NSW0356 | 44.59 | C | 13.5844 | hap2 | 29,764 | 24,635 | 7,763 | 36,712 | 22,953 | 231,054,062 |
| NSW0536 | 44.11 | A | 12.7913 | hap2 | 32,975 | 27,500 | 7,089 | 39,516 | 22,896 | 233,755,982 |
| NSW0354 | 43.29 | C | 14.2373 | hap2 | 29,040 | 24,054 | 7,711 | 35,144 | 21,541 | 223,925,553 |
| NSW0083 | 43.21 | B | 8.95098 | hap1 | 18,659 | 16,348 | 10,773 | 20,780 | 11,728 | 201,015,574 |
| NSW0299 | 38 | C | 9.98593 | hap1 | 13,100 | 11,088 | 14,452 | 15,771 | 9,511 | 189,326,335 |
| NSW0343 | 38.61 | C | 11.1506 | hap2 | 37,569 | 31,721 | 4,641 | 44,563 | 24,672 | 174,357,823 |
| NSW0575 | 37.14 | A | 8.35824 | hap1 | 19,149 | 17,056 | 9,225 | 21,242 | 10,629 | 176,644,650 |
| NSW0328 | 36.71 | C | 11.1906 | hap2 | 43,967 | 36,150 | 4,134 | 52,360 | 33,175 | 181,760,317 |
| NSW0113 | 31.81 | B | 11.3181 | hap2 | 34,306 | 27,857 | 5,136 | 40,054 | 27,180 | 176,196,528 |
| NSW0119 | 31.55 | B | 7.69559 | hap1 | 16,842 | 14,862 | 8,219 | 18,886 | 11,410 | 138,422,365 |
| NSW0335 | 31.99 | C | 10.751 | hap2 | 37,694 | 32,057 | 4,177 | 44,913 | 23,454 | 157,448,980 |
| NSW0972 | 31.51 | A | 11.1123 | hap2 | 32,158 | 26,939 | 4,885 | 37,448 | 22,556 | 157,093,964 |
| NSW0348 | 28.85 | C | 10.1471 | hap2 | 35,548 | 30,037 | 3,597 | 41,265 | 22,751 | 127,865,091 |
| NSW0315 | 27.99 | C | 6.49355 | hap1 | 24,122 | 20,990 | 4,659 | 27,553 | 14,370 | 112,385,863 |
| NSW0970 | 26.61 | A | 10.1207 | hap2 | 31,801 | 26,366 | 3,707 | 37,754 | 22,294 | 117,884,897 |
| NSW0573 | 25.88 | A | 12.383 | hap2 | 20,394 | 16,793 | 6,974 | 24,263 | 15,526 | 142,229,146 |
| NSW0111 | 25.58 | B | 7.71839 | hap1 | 15,157 | 12,913 | 8,899 | 18,135 | 10,910 | 134,884,111 |
| NSW0105 | 24.63 | B | 11.4163 | hap2 | 20,766 | 17,204 | 6,855 | 24,944 | 16,346 | 142,351,612 |
| NSW0347 | 23.73 | C | 9.58598 | hap2 | 33,867 | 28,881 | 3,075 | 39,472 | 21,440 | 104,140,489 |
| NSW0954 | 23.82 | A | 6.24472 | hap1 | 25,288 | 21,753 | 4,025 | 28,704 | 16,036 | 101,783,963 |
| NSW0175 | 20.59 | A | 9.84936 | hap2 | 32,866 | 27,394 | 3,308 | 38,857 | 23,069 | 108,720,548 |
| NSW0207 | 20.08 | A | 11.1162 | hap2 | 18,475 | 15,593 | 6,041 | 22,104 | 12,884 | 111,608,044 |
| NSW0966 | 18.61 | A | 6.39348 | hap1 | 17,200 | 14,685 | 4,228 | 19,468 | 11,857 | 72,720,101 |
| NSW0566 | 18.78 | A | 12.2311 | hap2 | 17,078 | 13,954 | 6,846 | 20,149 | 13,110 | 116,914,401 |
| NSW0555 | 15.99 | A | 11.2373 | hap2 | 17,208 | 13,819 | 4,818 | 20,549 | 15,110 | 82,907,674 |
| NSW0108 | 15.18 | B | 5.89502 | hap1 | 15,495 | 13,852 | 4,399 | 17,346 | 9,272 | 68,160,652 |
| NSW0083 | 9.24 | B | 8.95098 | hap2 | 21,249 | 18,281 | 2,389 | 23,485 | 17,719 | 50,763,044 |
| NSW0299 | 7.74 | C | 9.98593 | hap2 | 16,302 | 13,639 | 2,936 | 19,384 | 14,463 | 47,861,999 |
| NSW0575 | 6.83 | A | 8.35824 | hap2 | 21,558 | 18,333 | 1,731 | 24,342 | 15,440 | 37,316,211 |
| NSW0939 | 6.11 | A | 4.3979 | hap1 | 12,216 | 9,979 | 2,049 | 14,303 | 13,893 | 25,031,483 |

| | | | | | | | | | | |
|---|---|---|---|---|---|---|---|---|---|---|
| NSW0133 | 5.42 | B | 1.00533 | hap1 | 12,999 | 11,088 | 1,856 | 15,295 | 17,427 | 24,125,211 |
| NSW0133 | 5.42 | B | 1.00533 | hap2 | 12,951 | 11,088 | 1,842 | 15,186 | 15,987 | 23,856,154 |
| NSW0119 | 3.49 | B | 7.69559 | hap2 | 20,002 | 16,268 | 1,065 | 22,419 | 24,296 | 21,302,456 |
| NSW0194 | 3.23 | A | 4.04678 | hap1 | 14,395 | 11,986 | 1,069 | 16,836 | 15,621 | 15,388,655 |
| NSW0954 | 3.06 | A | 6.24472 | hap2 | 27,621 | 23,009 | 541 | 30,764 | 23,414 | 14,943,080 |
| NSW0315 | 2.75 | C | 6.49355 | hap2 | 28,804 | 23,845 | 571 | 33,159 | 22,767 | 16,446,969 |
| NSW0111 | 2.71 | B | 7.71839 | hap2 | 19,511 | 15,968 | 1,338 | 22,716 | 16,889 | 26,105,869 |
| NSW0963 | 2.24 | A | 4.04471 | hap1 | 13,226 | 11,274 | 1,028 | 14,918 | 13,527 | 13,596,083 |
| NSW0192 | 2.06 | A | 3.55784 | hap1 | 13,592 | 11,371 | 783 | 15,911 | 15,257 | 10,642,654 |
| NSW0966 | 1.42 | A | 6.39348 | hap2 | 22,514 | 17,738 | 444 | 26,181 | 25,232 | 9,996,073 |
| NSW0940 | 1.2 | A | 3.20183 | hap1 | 12,505 | 9,736 | 437 | 14,141 | 16,856 | 5,464,778 |
| NSW0120 | 0.99 | B | 3.36433 | hap1 | 13,049 | 10,269 | 479 | 14,475 | 18,775 | 6,250,283 |
| NSW0108 | 0.86 | B | 5.89502 | hap2 | 18,742 | 15,794 | 440 | 20,743 | 18,608 | 8,246,245 |
| NSW0194 | 0.9 | A | 4.04678 | hap2 | 16,722 | 12,954 | 326 | 18,536 | 25,152 | 5,451,270 |
| NSW0963 | 0.95 | A | 4.04471 | hap2 | 16,164 | 12,943 | 319 | 17,557 | 21,887 | 5,156,270 |
| NSW0939 | 0.56 | A | 4.3979 | hap2 | 18,832 | 13,931 | 231 | 22,391 | 26,325 | 4,350,083 |
| NSW0935 | 0.47 | A | 2.77763 | hap1 | 11,526 | 7,646 | 366 | 13,891 | 22,458 | 4,218,391 |
| NSW0209 | 0.39 | A | 2.39105 | hap1 | 11,634 | 9,309 | 205 | 14,924 | 12,026 | 2,384,899 |
| NSW0192 | 0.34 | A | 3.55784 | hap2 | 18,521 | 13,397 | 211 | 20,835 | 26,583 | 3,907,861 |
| NSW0120 | 0.26 | B | 3.36433 | hap2 | 19,429 | 12,870 | 135 | 20,567 | 31,756 | 2,622,878 |
| NSW0209 | 0.26 | A | 2.39105 | hap2 | 12,415 | 9,818 | 136 | 15,372 | 13,614 | 1,688,383 |
| NSW0935 | 0.26 | A | 2.77763 | hap2 | 12,359 | 8,231 | 231 | 14,489 | 21,278 | 2,854,909 |
| NSW0938 | 0.3 | A | 2.33443 | hap1 | 12,667 | 8,809 | 235 | 16,530 | 16,705 | 2,976,792 |
| NSW0110 | 0.21 | B | 2.04307 | hap1 | 20,843 | 13,308 | 96 | 23,739 | 44,267 | 2,000,950 |
| NSW0938 | 0.21 | A | 2.33443 | hap2 | 14,441 | 9,365 | 145 | 18,588 | 19,016 | 2,093,878 |
| NSW0940 | 0.17 | A | 3.20183 | hap2 | 19,011 | 12,053 | 124 | 20,198 | 29,443 | 2,357,384 |
| NSW0110 | 0.13 | B | 2.04307 | hap2 | 25,362 | 13,411 | 59 | 35,391 | 55,638 | 1,496,370 |
| NSW0936 | 0.13 | A | 1.81097 | hap1 | 14,616 | 8,967 | 119 | 19,628 | 20,827 | 1,739,330 |
| NSW0936 | 0.13 | A | 1.81097 | hap2 | 14,289 | 8,855 | 116 | 19,124 | 20,894 | 1,657,519 |

**Supplementary Table 9 Size and frequency of SVs discovered by Sniffles2**

| Variant type | Count | Median size | Mean size | Total bases (Mb) | Percentage (%) |
|---|---|---|---|---|---|
| Deletions | 339,168 | 119 | 807.44 | 273.86 | 59.10 |
| Insertions | 229,942 | 231 | 641.01 | 147.39 | 40.07 |
| Duplications | 3,280 | 12,467.5 | 20,343.52 | 66.73 | 0.57 |
| Inversions | 1,499 | 3,709 | 16,788.48 | 25.17 | 0.26 |

| | | | | | |
|---|---|---|---|---|---|
| **Total** | 573,889 | 16,526.5 | 38,580.45 | **513.14** | 100.00 |

**Supplementary Table 10. Percent of different chromosomes of the reference genome covered by SVs discovered by Sniffles2**

| Chr | Percent of chromosome with SVs (%) |
|---|---|
| 1 | 37.57 |
| 2 | 40.15 |
| 3 | 40.44 |
| 4 | 39.11 |
| 5 | 46.25 |
| 6 | 32.8 |
| 7 | 40.29 |
| 8 | 40.51 |
| 9 | 34.65 |
| 10 | 33.79 |
| 11 | 33.67 |
| **Overall** | **38.46** |

**Supplementary Table 11. Number of SVs intersecting with different gene features**

| SV type | 5 kb upstream | Genic | Exonic | Introngenic | 5 kb downstream | Intergenic |
|---|---|---|---|---|---|---|
| Deletion | 19,705 | 15,039 | 3,546 | 10,789 | 19,612 | 291,341 |
| Insertion | 17,661 | 12,361 | 5,266 | 8,167 | 17,345 | 190,302 |
| Duplication | 617 | 623 | 568 | 548 | 596 | 2,353 |
| Inversion | 219 | 180 | 155 | 166 | 216 | 1,160 |
| All | 38,202 | 28,203 | 9,535 | 19,670 | 37,769 | 485,156 |
| Percentage | **6.66%** | **4.91%** | **1.66%** | **3.43%** | **6.58%** | **84.54%** |

**Supplementary Table 12. SV-GWAS results for *CHILL1* and *CHILL8***

| SV type | Size (bp) | Locus | Chr | Pos | GWAS -log10 (p-value) | Freq (%) | Effect (°C) | SE | % variation | Associated genes |
|---|---|---|---|---|---|---|---|---|---|---|
| | | **Intercept (BioClim6)** | | | | | **-0.85** | **0.18** | | |
| | | **Full model** | | | | | | | **84** | |
| Insertion | 485 | *CHILL-1.1* | 1 | 27,696,962 | 8.0 | 45 | -1.95 | 0.15 *** | 69 | |
| Deletion | 64 | *CHILL-1.2* | 1 | 27,698,414 | 9.0 | 43 | -2.03 | 0.14 *** | 73 | |
| **Deletion** | **58** | ***CHILL-1.3*** | **1** | **27,713,800** | **12.5** | **43** | **-2.06** | **0.13 ***** | **76** | GPI-anchor protein, Chlorophyll a reductase |
| Deletion | 402 | *CHILL-1.4* | 1 | 27,736,703 | 7.8 | 45 | -1.96 | 0.15 *** | 68 | |
| Insertion | 323 | *CHILL-1.5* | 1 | 27,799,070 | 9.2 | 43 | -2.04 | 0.14 *** | 74 | |
| Insertion | 1024 | *CHILL-4.1* | 4 | 25,787,765 | 8.2 | 47 | -1.96 | 0.14 *** | 71 | |
| Deletion | 123 | *CHILL-6.1* | 6 | 34,505,641 | 9.9 | 49 | -1.92 | 0.13 *** | 76 | |
| **Deletion** | **60** | ***CHILL-8.1*** | **8** | **38,494,586** | **7.1** | **10** | **1.25** | **0.26 ***** | **64** | Phosphatase, Phosphate transporter |
| Deletion | 52 | *CHILL-8.2* | 8 | 38,503,991 | 8.5 | 51 | -1.85 | 0.14 *** | 71 | |
| Deletion | 221 | *CHILL-8.3* | 8 | 38,507,136 | 9.0 | 51 | -1.86 | 0.13 *** | 72 | |

**Supplementary Table 13. Characterisation of genes residing at the *CHILL8* locus.** Expression was detected by Oxford Nanopore Technologies direct RNA sequencing and reported as Transcripts Per Million (TPM).

| Gene ID | Gene name | Protein length | Arabidopsis ortholog | Expression (TPM) |
|---|---|---|---|---|
| g34497.t1 | probable inorganic phosphate transporter 1-3 [Eucalyptus grandis] | 534 | AT5G43350 phosphate transporter 1;1 (PHT1;1) | 1,476.39 (reverse strand) |
| g34498.t1 | serine/threonine-protein phosphatase PP1 isozyme 3 isoform X1 [Eucalyptus grandis] | 324 | AT2G39840 type one serine/threonine protein phosphatase 4 (TOPP4) | 40.27 (forward strand) |

**Supplementary Table 14. Characterisation of genes residing at the *CHILL1* locus.** Expression was detected by Oxford Nanopore Technologies direct RNA sequencing and reported as Transcripts Per Million (TPM). Uncharacterised genes with no match or ortholog are labelled NA.

| Gene ID | Gene name | Protein length | Arabidopsis ortholog | Expression (TPM) |
|---|---|---|---|---|
| g2294.t1 | α-Terpineol synthase | 711 | Terpene Synthase 4 AT4G16740 | 0 |
| g2295.t1 | GPI-anchored protein | 205 | GPI-anchored protein At1G61900 | 0 |
| g2296.t1 | NA | 169 | NA | 0 |
| g2297.t1 | 7-hydroxymethyl chlorophyll a reductase | 350 | AT1G04620 | 13.42 |
| g2298.t1 | (+)-α-Terpineol synthase-like [Syzygium oleosum] | 714 | terpene synthase-like sequence-1,8-cineole AT3G25820 | 0 |
| g2299.t1 | GPI-anchored protein | 212 | GPI-anchored protein At1G61900 | 0 |

**Supplementary Table 15. Results from linear models exploring relationships between BioClim6, geography, *CHILL1*, species and principal components (PCs) based on genome-wide and *CHILL1* SNPs.**

| Dependent variable | Factors in model | Variance explained by model ($R^2$) (%) | Factor level | P-value | Effect size | Units for effect size | Change required to achieve equivalent shift in BioClim6 from -2⁰C to 0⁰C |
|---|---|---|---|---|---|---|---|
| BioClim6 | Latitude + elevation | 72.5 | Latitude | $10^{-16}$ | 0.50 | Latitude degrees (111km) | Translocate from latitude -36º to -32º (444km) if at 1200m |
| | | | Elevation | $10^{-16}$ | -0.35 | 100m elevation | Translocate from altitude of 1200m to 600m if at latitude of -36⁰ |
| BioClim6 | *CHILL1* | 15.8 | *CHILL1/CHILL1* | - | 0[a] | - | From *CHILL1/CHILL1* to *chill1/chill1* |
| | | | *CHILL1/chill1* | $10^{-6}$ | 0.83 | One allele | |
| | | | *chill1/chill1* | $10^{-16}$ | 1.59 | Two alleles | |
| BioClim6 | Species | 0.8 | *E. dalrympleana* | - | 0[a] | - | - |
| | | | *E. rubida* | 0.37 | -0.17 | | - |
| | | | *E. viminalis* | 0.36 | 0.164 | | - |
| BioClim6 | Latitude + elevation + *CHILL1* | 74.7 | Latitude | $10^{-16}$ | 0.59 | - | - |
| | | | Elevation | $10^{-16}$ | -0.34 | - | - |
| | | | *CHILL1/CHILL1* | - | 0[a] | | |
| | | | *CHILL1/chill1* | 0.02 | -0.24 | - | - |
| | | | *chill1/chill1* | $10^{-9}$ | -0.82 | - | - |
| BioClim6 | Latitude + | 73.8 | Latitude | $10^{-16}$ | 0.49 | - | - |

| | | | | | | | |
|---|---|---|---|---|---|---|---|
| | elevation + species | | | | | | |
| | | | Elevation | $10^{-16}$ | -0.38 | - | - |
| | | | *E. dalrympleana* | - | 0[a] | - | - |
| | | | *E. rubida* | 0.007 | -0.29 | - | - |
| | | | *E. viminalis* | $10^{-7}$ | -0.44 | - | - |
| BioClim6 | Latitude + elevation + species + *CHILL1* | 75.4 | Latitude | $10^{-16}$ | 0.57 | - | - |
| | | | Elevation | $10^{-16}$ | -0.37 | - | - |
| | | | *CHILL1/CHILL1* | - | 0[a] | - | - |
| | | | *CHILL1/chill1* | 0.007 | -0.18 | - | - |
| | | | *chill1/chill1* | $10^{-7}$ | -0.74 | - | - |
| | | | *E. dalrympleana* | - | 0[a] | - | - |
| | | | *E. rubida* | 0.006 | -0.29 | - | - |
| | | | *E. viminalis* | 0.001 | -0.32 | - | - |
| BioClim6 | Genome-wide PC1 & PC2 | 0.9 | PC1 | 0.10 | 0.002 | - | - |
| | | | PC2 | 0.250 | 0.002 | - | - |
| BioClim6 | Genome-wide PC3 & PC4 | 54.00 | PC3 | $10^{-16}$ | 0.031 | - | - |
| | | | PC4 | 0.07 | -0.004 | - | - |
| BioClim6 | *CHILL1* + PC1 & PC2 | 15.0 | PC1 | $10^{-16}$ | 0.022 | - | - |
| | | | PC2 | 0.79 | 0.003 | - | - |

| | | | | | | | |
|---|---|---|---|---|---|---|---|
| BioClim6 | *CHILL1* + PC3 & PC4 | 0.04 | PC3 | 0.68 | 0.005 | - | - |
| | | | PC4 | 0.99 | 0.0001 | - | - |
| *CHILL1* | BioClim6 | [b]14.4 | BioClim6 | $10^{-11}$ | [c]0.58 | | Decrease in frequency of *CHILL1* allele from 0.88 to 0.69. |
| | BioClim6 + species | 24.0 | BioClim6 | $10^{-11}$ | 0.60 | | Decrease in frequency of *CHILL1* allele from 0.79 to 0.53 if *E. viminalis.* |
| | | | *E. dalrympleana* | - | 0 | | - |
| | | | *E. rubida* | 0.005 | -1.19 | | |
| | | | *E. viminalis* | 0.002 | 0.86 | | |
| *CHILL1* | BioClim6 + Genome-wide PC1 + Genome-wide PC2 | 22.1 | BioClim6 | $10^{-11}$ | 0.60 | | - |
| | | | PC1 | $10^{-7}$ | 0.011 | | - |
| | | | PC2 | 0.60 | 0.0026 | | - |
| *CHILL1* | Latitude + elevation | 37.3 | Latitude | $10^{-17}$ | 0.68 | | |
| | | | Elevation | 0.08 | -0.13 | | - |
| *CHILL1* | BioClim6 + latitude + elevation | 40.1 | BioClim6 | 0.0007 | -0.65 | | - |
| | | | Latitude | $10^{-15}$ | 1.020 | | - |

| | | | | | | |
|---|---|---|---|---|---|---|
| | | | Elevation | 0.0006 | -0.35 | - |
| *CHILL1* | BioClim6 + Genome-wide PC1 + Genome-wide PC2 + Genome-wide PC3 + Genome-wide PC4 | 46.2 | BioClim6 | 0.3 | -0.16 | |
| | | | PC1 | 0.001 | -0.020 | - |
| | | | PC2 | 0.50 | 0.006 | - |
| | | | PC3 | $10^{-10}$ | 0.103 | - |
| | | | PC4 | $10^{-7}$ | -0.074 | - |

a. Reference factor level against which estimates for other factor levels are compared. “-” denotes irrelevant or uninterpretable.
b. McFadden’s $R^2$ was used for models where the dependent variable was binomial.
c. Estimates of effects are on the logit scale but translated to the response scale in the table’s last column.

**Supplementary Table 16. Frequencies of *CHILL1* genotypes and alleles by species and geographic population**

| Species | | No. of genotypes | [a]Population | | |
|---|---|---|---|---|---|
| | | | South | Central | North |
| ***E. viminalis*** | | | | | |
| *CHILL1/CHILL1* | | 89 | 0.83 | 0.08 | 0.03 |
| *CHILL1/chill1* | | 43 | 0.15 | 0.51 | 0.24 |
| *chill1/chill1* | | 44 | 0.02 | 0.38 | 0.74 |
| | Total | 176 | 102 | 26 | 38 |
| ***E. dalrympleana*** | | | | | |
| *CHILL1/CHILL1* | | 102 | 0.89 | 0.92 | 0.09 |
| *CHILL1/chill1* | | 15 | 0.11 | 0.06 | 0.22 |
| *chill1/chill1* | | 21 | 0.03 | 0.04 | 0.70 |
| | Total | 138 | 66 | 48 | 24 |
| ***E. rubida*** | | | | | |
| *CHILL1/CHILL1* | | 103 | 0.83 | 0.90 | 0.00 |
| *CHILL1/chill1* | | 16 | 0.15 | 0.11 | 0.00 |
| *chill1/chill1* | | 1 | 0.01 | 0.00 | 0.00 |
| | Total | 120 | 82 | 38 | 0 |
| **All species** | | | | | |
| *CHILL1/CHILL1* | | 294 | 0.84 | 0.66 | 0.05 |
| *CHILL1/chill1* | | 74 | 0.14 | 0.21 | 0.23 |
| *chill1/chill1* | | 66 | 0.02 | 0.13 | 0.73 |
| | Total | 434 | 250 | 122 | 62 |

a. South, below $-35^0$ latitude; North, greater than $-30^0$; Central, between $-35^0$ and $-30^0$
