## Supplemental Notes for "Complex Genomic Structural Variation Underlies Climate Adaptation across *Eucalyptus* species"

### Supplementary Notes

### Note 1: Hi-C validation of *Eucalyptus viminalis* reference genome chromatin structure

To achieve a haplotype-resolved, chromosome-scale, reference for *E. viminalis* ACT, we used Hi-C chromosome conformation capture data with long-read contigs, to partition the haplotypes and scaffold into chromosomes (22 haplotypes). This approach also validated the quality of the *de novo* genome assembly; no error correction of contigs was required. However, ongoing advancements in long-read sequencing technologies, combined with improved assembly algorithms, have increasingly diminished the need for Hi-C. Emerging telomere-to-telomere *Eucalyptus* assemblies now routinely demonstrate near-complete contiguity, with only a few structural gaps, occurring primarily in complex repeat regions such as the centromere of chromosome 2 (i.e. the single gap remaining in *E. viminalis* VIC and TAS).

**a** Haplotype 1 **b** Haplotype 2

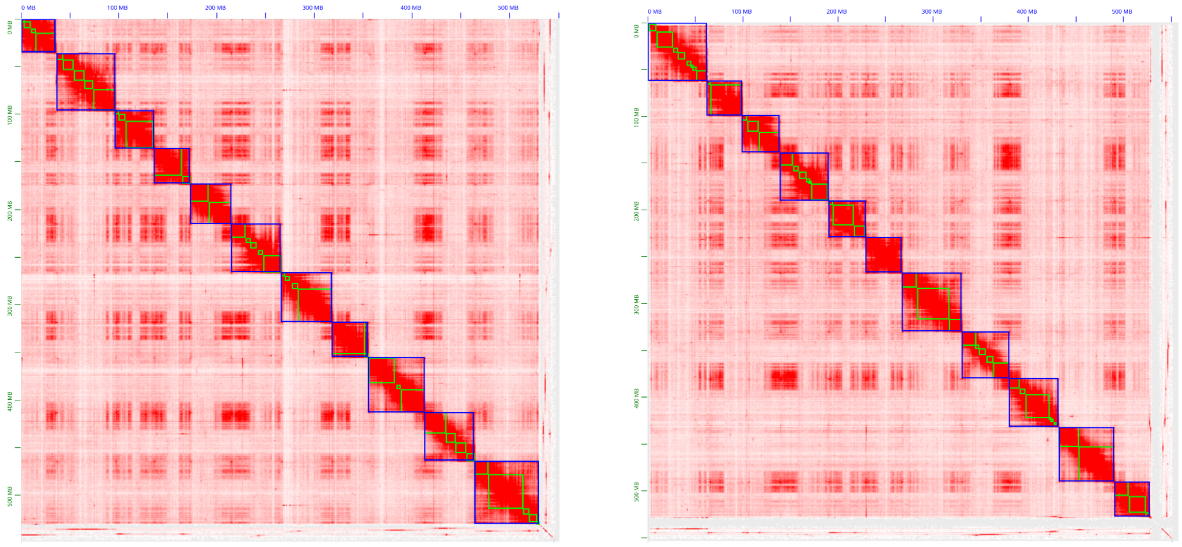

**Supplementary Figure 1. Chromosome-scale scaffolding and validation of the *E. viminalis* ACT reference genome using Hi-C.** Hi-C contact maps showing the scaffolding of contigs into haplotype-resolved chromosomes for the *E. viminalis* ACT reference genome. **a** Haplotype 1, **b** Haplotype 2. Contact maps were generated and visualized using Juicebox (Durand et al., 2016).

### Note 2: *Eucalyptus viminalis* genomic sequence-based synteny is conserved across species range

To further investigate structural variations across chromosomes, we analysed large interchromosomal SVs (Fig. S2). We identified genome-wide duplications and translocations ≥10 kbp in length (with ≥95% sequence identity), including events spanning non-homologous chromosomes. The spatial proximity required for such inter-chromosomal rearrangements may reflect underlying three-dimensional genome architecture (Hoencamp et al., 2021). Trans-chromosomal interactions observed in Hi-C contact maps support this (Supplementary Figure 1). The size distribution of duplications (5-20 kbp) correlates to annotated LTR retrotransposons (Gypsy and Copia superfamilies), implicating recent transposable element activity as the primary driver of structural variation. The inter-chromosomal distribution of these TE-associated duplications suggests that three-dimensional genome organisation may facilitate rearrangements between spatially proximate chromosomes.

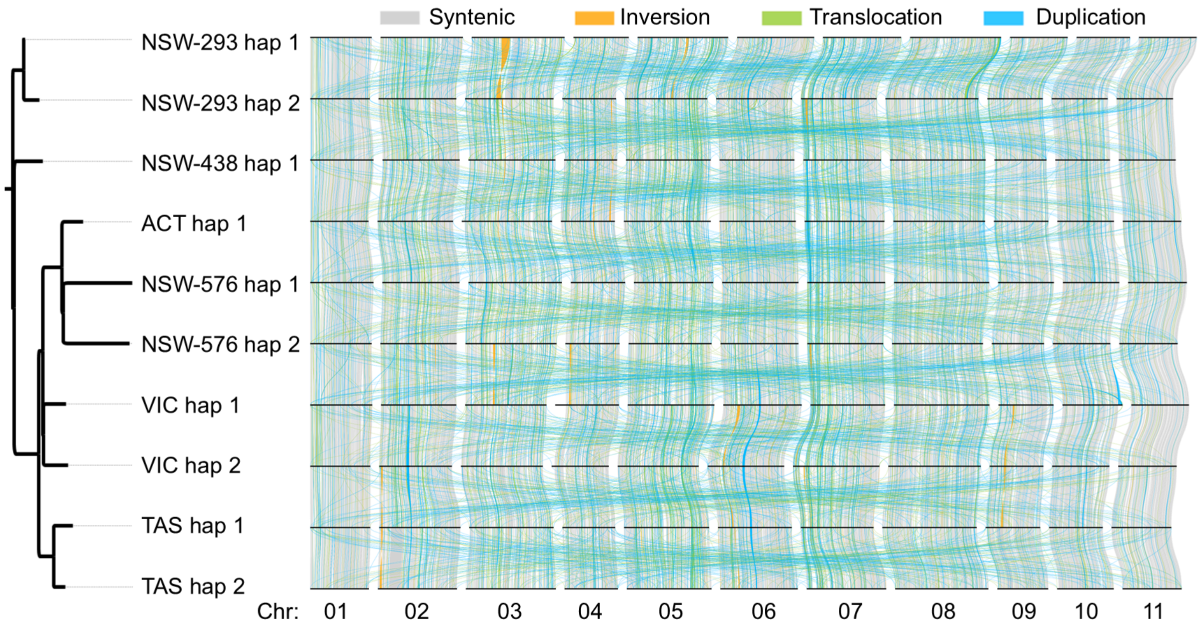

**Supplementary Figure 2. Phylogenetic relationships, synteny, and structural variation between genomes, including interchromosomal rearrangements.** Both intrachromosomal and interchromosomal structural variants ≥10 kb are displayed, showing duplications and translocations across chromosomes.

### Note 3: Reference RNA quality check and ONT direct RNA sequencing

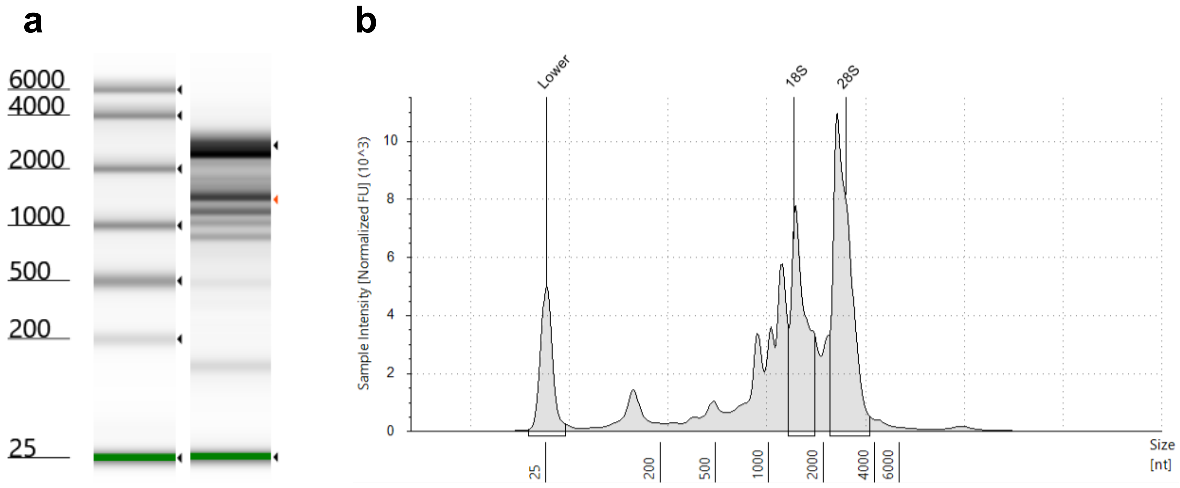

**Supplementary Figure 3. Quality check of total RNA using an Agilent TapeStation RNA ScreenTape.** (a) Electrophoresis image. First column is the sizes in nucleotides, second column an RNA ladder, and third column is *Eucalyptus viminalis* total RNA. The green band is an internal lower marker added to each sample (25 nucleotides in length). (b) Electropherogram, plotting fragment size (nucleotides, nt), against sample intensity (normalised fluorescence units, FU). Three labelled peaks are the internal lower marker, 18S rRNA and 28S rRNA peaks. Applying the TapeStation eukaryotic model to calculate an RNA integrity number equivalent (RINe), a value of 7.8 was calculated. Noting this does not account for plant rRNA, the value was deemed high quality.

### Note 4: Limitations and challenges of pan-genome graphs for wild plant species

While pan-genome graphs can illuminate structural variation, in our data, using chromosome 1 as an example, we found that different runs of Pangenome Graph Builder (PGGB) generated variable graph structures and pangenome size. This is likely to be due to repetitive and complex parts of the genome that may, in turn, be the result of current and historically active transposable element activity. Additionally, PGGB reveals large loops (51 self-loops, all unique) and large inversions throughout the chromosome which are likely to be artefactual. Furthermore, genome-based haplotype comparison methods in non-model species can be subject to biases and errors arising from assembly algorithms. Thus, genome-wide graph representations in wild plant species such as *Eucalyptus* are often difficult to interpret.

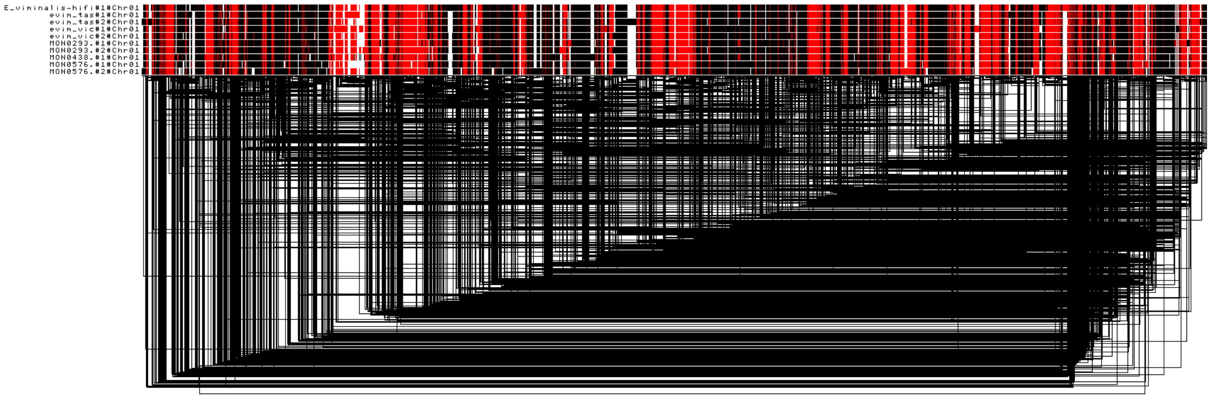

**Supplementary Figure 4. ODGI 1D graph of chromosome 1.** Red and black show inverted segments. Black lines below are links between nodes. Analyses performed using 10 haplotypes.

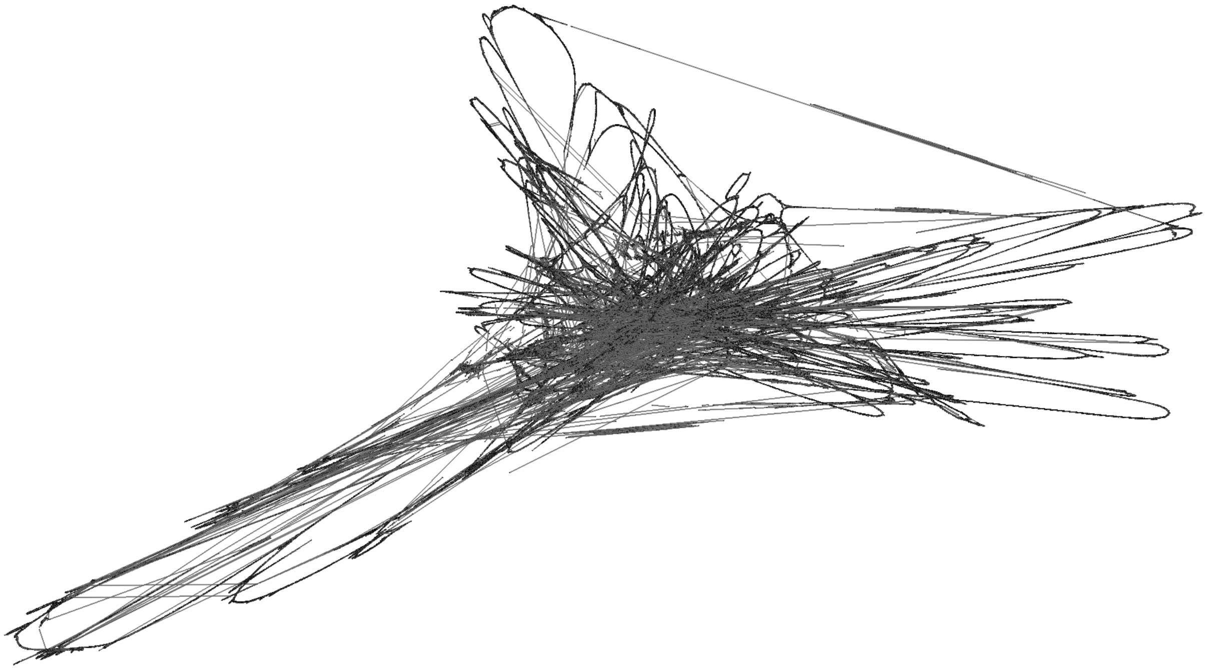

**Supplementary Figure 5. ODGI 2D graph visualisation of chromosome 1.** Pangenome graph generated using 10 haplotypes. 2D graph showing large self-loops, likely induced by repetitive genomic content.

### Note 5: Gene-based pangenome - extended results

Preservation of gene orders, or gene-based synteny has important implications of post-zygotic reproductive barriers. This affects sexual reproduction within a species and hybridisation between species. Here, we use ordered gene orthogroups identified by Orthofinder (version 2.5) to investigate gene-based synteny with GENESPACE (https://github.com/jtlovell/GENESPACE). Gene synteny *Eucalyptus viminalis* synteny (gene-based) was conserved across its ecological range, allowing sexual reproduction and exchange of alleles (Fig S6).

The accessory part of the orthogroup-based pangenome showed enrichments in annotations for signal transduction, secondary metabolite biosynthesis, and DNA repair. More detailed functional classifications showed enrichment in many disease-resistance related pathways such as Leucine-rich repeats (LRRs). Other pathways such as synthase and many kinases families also showed variable presence-absence variations (PAVs). These results suggest many dimensions of within-species functional variation within the wide-ranging *Eucalyptus viminalis* species.

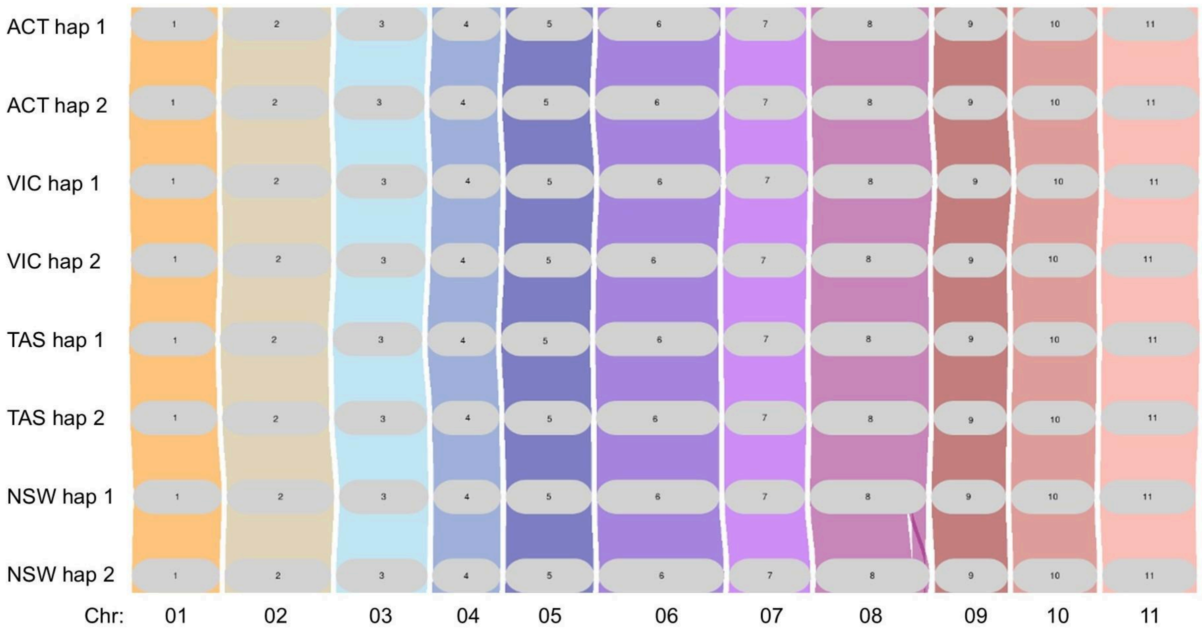

**Supplementary Figure 6. Gene orthogroup synteny is mostly conserved across representative genomes across four states and territories of Australia.** Orthrogroups identified by OrthroFinder were used by GENESACE as anchors to show the preservation of synteny-resolved orthogroups (synthegroups) across genographic range. ACT: Australian Capital Territory, VIC: Victoria, TAS: Tasmania, and NSW: New South Wales.

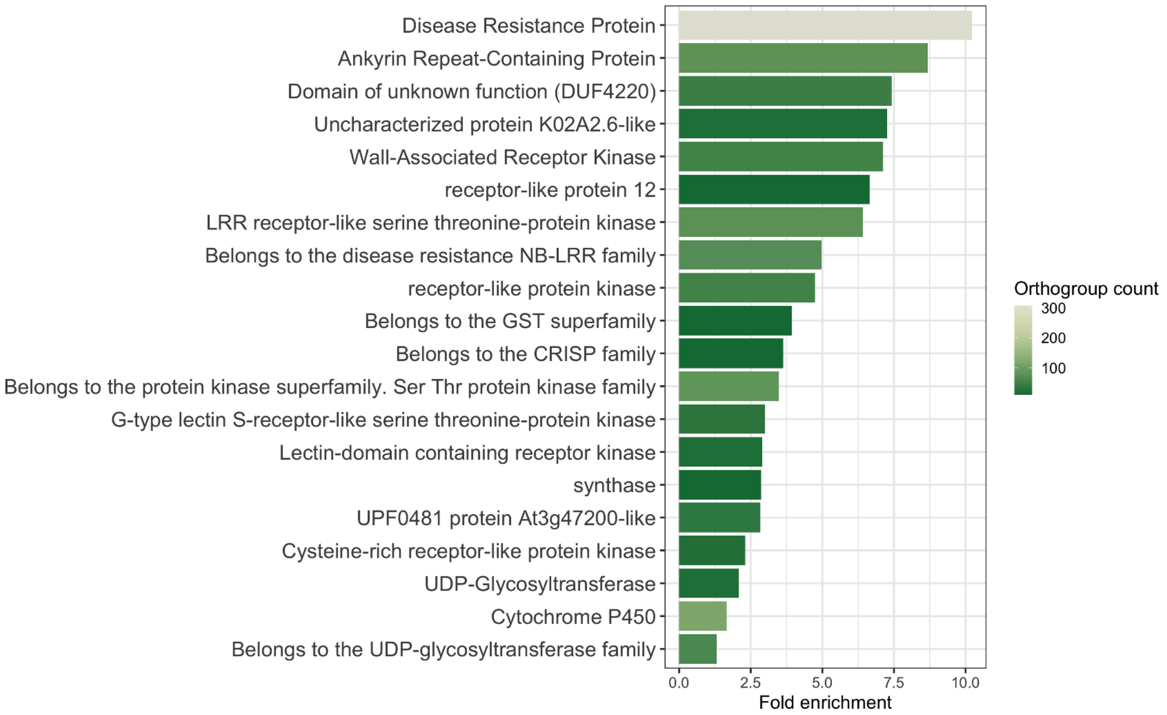

**Supplementary Figure 7. Pathways enriched in presence and absence variations of genes.** Orthogroups with variable presence or absence across haplotypes (i.e., the accessory pangenome). They are enriched in many functional genes such as Cytochrome P450 and disease resistance related to stress tolerance.

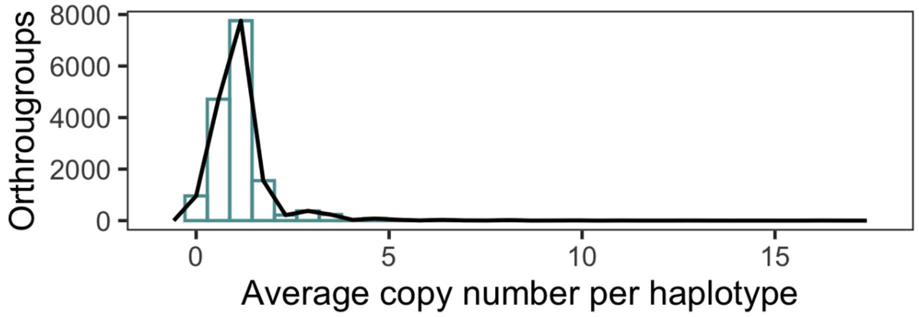

**Supplementary Figure 8. Some orthogroups have high copy numbers.** In addition to absence and presence variation, different orthogroups can also exhibit heterogeneous copy numbers. This can contribute to sequence and functional difference among haplotypes and individuals.

#### Note 6: Assembly-based SV characterisation - extended results

Structural variants typically represent ~50% of total (pan)genomic polymorphic sequences (Alonge et al., 2020; Lian et al., 2024; Liao et al., 2023), yet they can be represented inconsistently (e.g., Sniffles, PGGB, and Pannagram). Traditionally, SVs with different breakpoints would be considered as independent variants, which may not be true. Comparing many genome assemblies with few gaps and mis-assemblies provides a holistic, albeit complicated, representation of SVs to help us gain deeper insights over their generation mechanisms. PGGB identified 266 Mb of non-SNP sequences not present in the reference whereas Pannagem identified 164 Mb of non-SNP sequences not in the reference.

Among sexually reproducing populations, complex, or non-biallielic structural variants can be either independent or nested events, mechanistically affecting their inheritance. Complex SVs are identified from their inconsistent breakpoints, types, or both (Audano and Beck, 2024). Pannagram is a novel tool for identifying such variants as an alternative to the graph-based pangenomes (e.g., PGGB) using multiple genomes alignments (https://github.com/iganna/pannagram). In our target species, *E. viminalis*, we found that SVs consistently consist of 10 - 30% complex variants across all 11 chromosomes (Fig. S9).

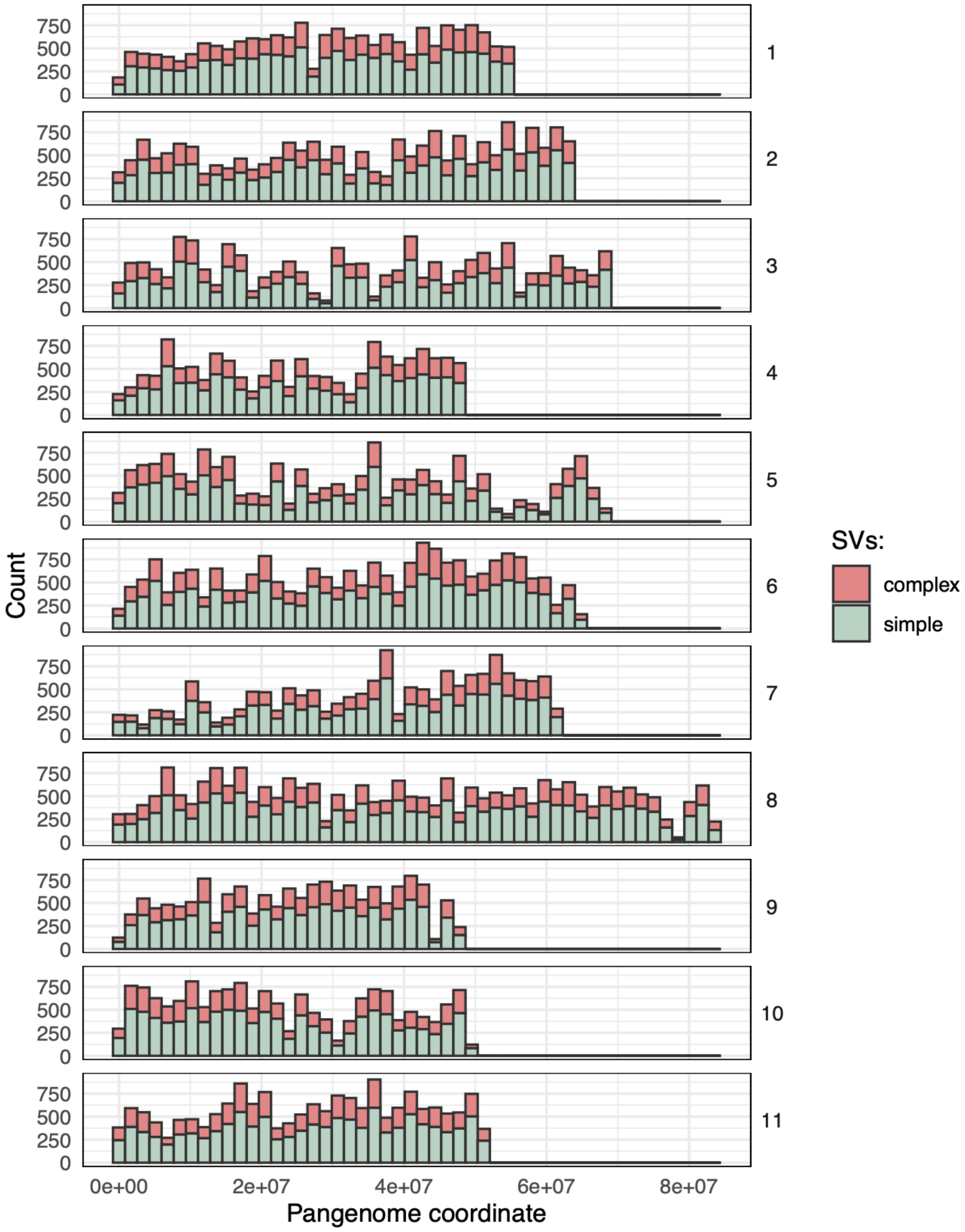

**Supplementary Figure 9. Complex structural variants consistently occur across chromosomes.** Simple and complex structural variants (sSVs and cSVs) are defined by being bi-allelic and multi-allelic in the 10 input haplotypes. Consistently, 10 - 30% of all SVs are considered complex due to having multiple (potentially related) variants with overlapping breakpoints.

We investigated the sequence homology network of SVs called by Pannagram. Clusters of SVs with similar sequence identities were found from SV sequences (Fig. S10). The majority of TE-associated SVs were between 800 bp and 3 kb in size, similar to typical TE sizes (Fig. S10). These SVs with sequence homology might have originated from the same diversification events. Among other biological mechanisms, TE activity is a commonly investigated source of novel sequences and genome perturbation. We therefore directly investigated the relationship between SVs and TEs.

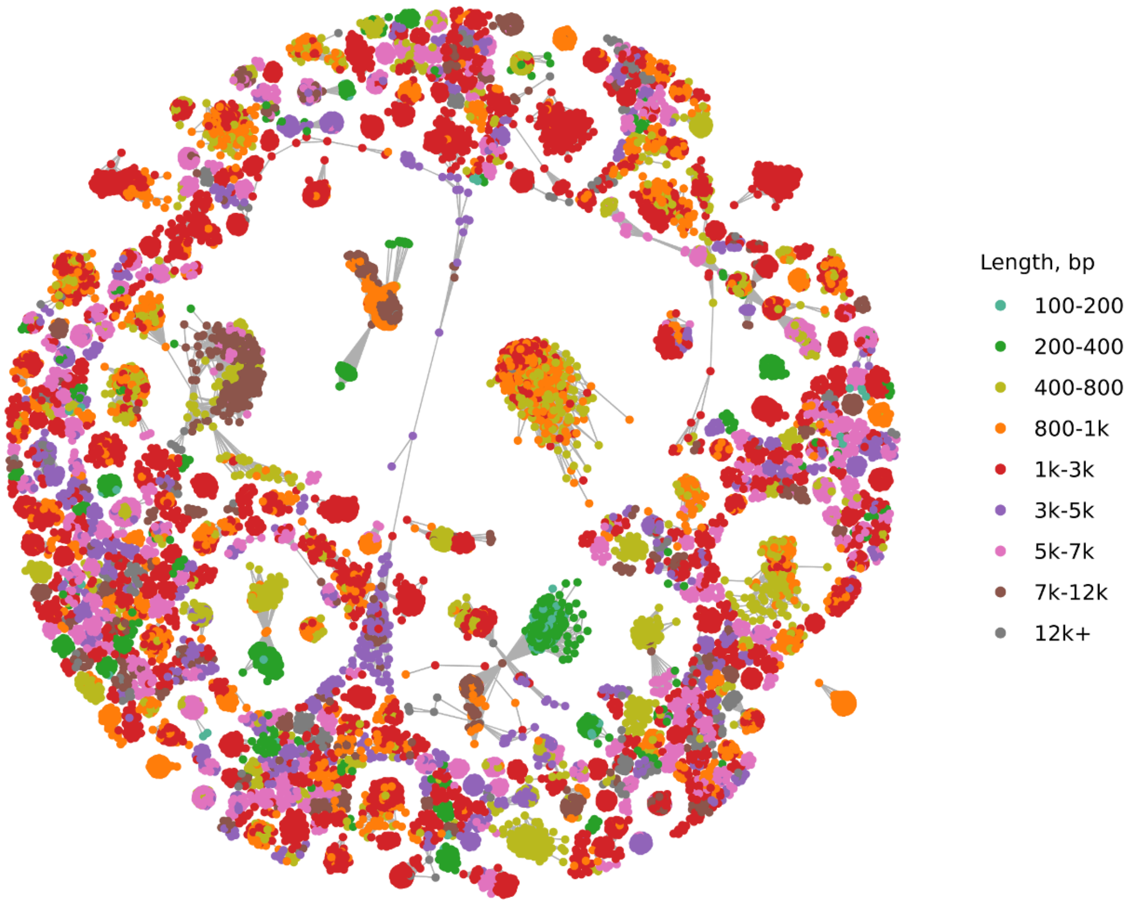

**Supplementary Figure 10. Simple structural variants form clusters according to reciprocal sequence similarity.** Simple SVs were investigated further with sequence homology networks to explore their relationships and infer common source of origin. Edges were connected if sequences were 90% similar with 90% overlapping sequences. SVs (nodes) colored by size range.

### Note 7: Transposons - extended results

After the initial characterisation, we identified causal or associative relationships between SVs and TEs. First, the genomes were *de novo* TE annotated using Extensive *de novo* TE Annotator (EDTA). Distributions of TE sequence divergence were visualised, showing peaks at 0% and 9% sequence divergence. This suggests recent TE expansion events that may have played important roles in the evolution and adaptation of this species (Fig. S11). Across the genome, a permutation test revealed a significant ($p < 0.001$, $z = 141.27$) enrichment of SV-TE association, which is 15% higher than expected by random chance. Using the sequence similarity search function of Pannagram, 13% of SV sequences were identified as TEs of different types (with over 85% sequence similarity and at least 85% coverage). TE-associated SVs were then clustered into sequence homology networks (Fig. S12). Across all TE-associated SVs, nearly half (48%) of all TE-associated SVs remain uncharacterised beyond the order level. Of the SV identified as TEs belonging to known families, the most common was Copia (29%), a long-terminal repeat (LTR) retrotransposon (Class I) superfamily that occurs across plants, metazoans and fungi (Fig. S13). Copia is commonly found in *Eucalyptus* and was suggested to be actively expanding (Marcon et al., 2011) and inserting polymorphic sequences (Marcon et al., 2015).

However, we also discovered a cluster of longer SV sequences (7 kb - 12 kb) associated with long-terminal repeat (LTR) retrotransposons (Class I). From these larger TE-associated SVs, we extracted long open reading frames (ORFs) that reached over 1000 unbroken amino acids of peptide sequence (~ 3 kb coding sequence; Fig. S14). The peptide sequence had homology in Retrovirus-related Pol polyprotein in transposon of common grape vines (Vitis vinifera; Fig. S14). Moreover, ONT direct RNA seq of *E. viminalis* confirmed the expression of this transcript in the reference *E. viminalis* with 566 primary alignments of intact transcripts. These results suggested recent ongoing TE activities in *E. viminalis* that might have contributed to the evolution and adaptation of this species.

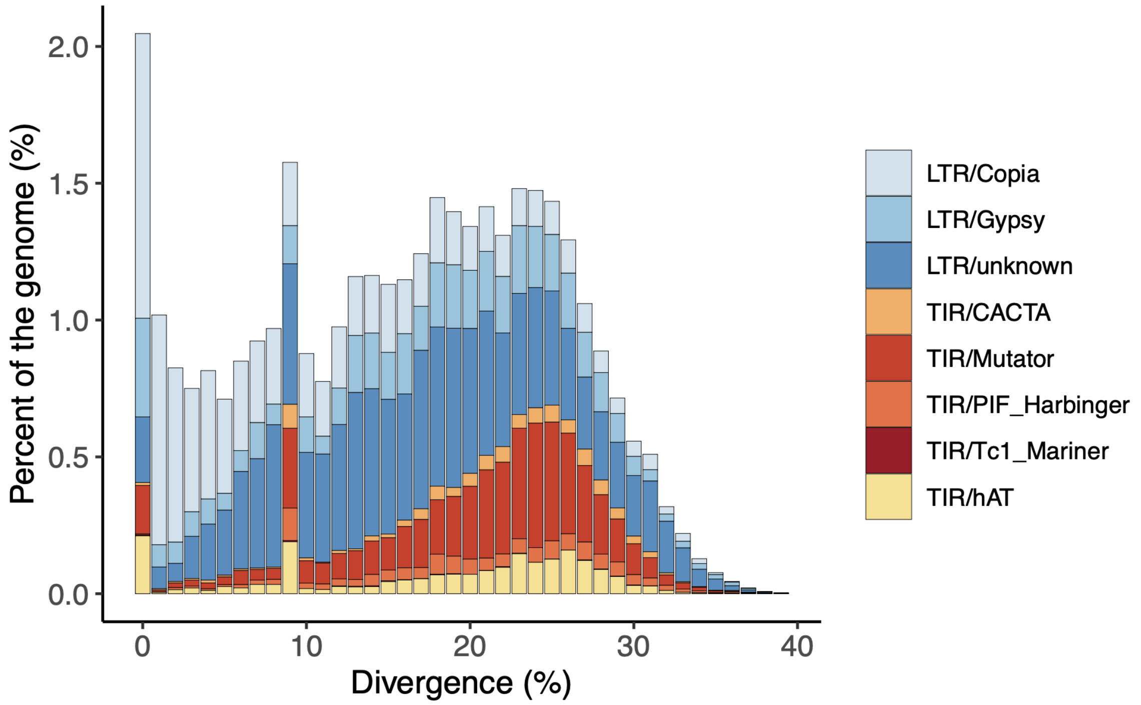

**Supplementary Figure 11. Two peaks of TEs with low sequence divergence suggest ongoing and previous TE expansion events.** Transposons were annotated with Extensive *de novo* TE Annotator (EDTA) and their sequence similarities analysed. Two peaks were identified (at 0% and 9% respectively), suggesting recent activities.

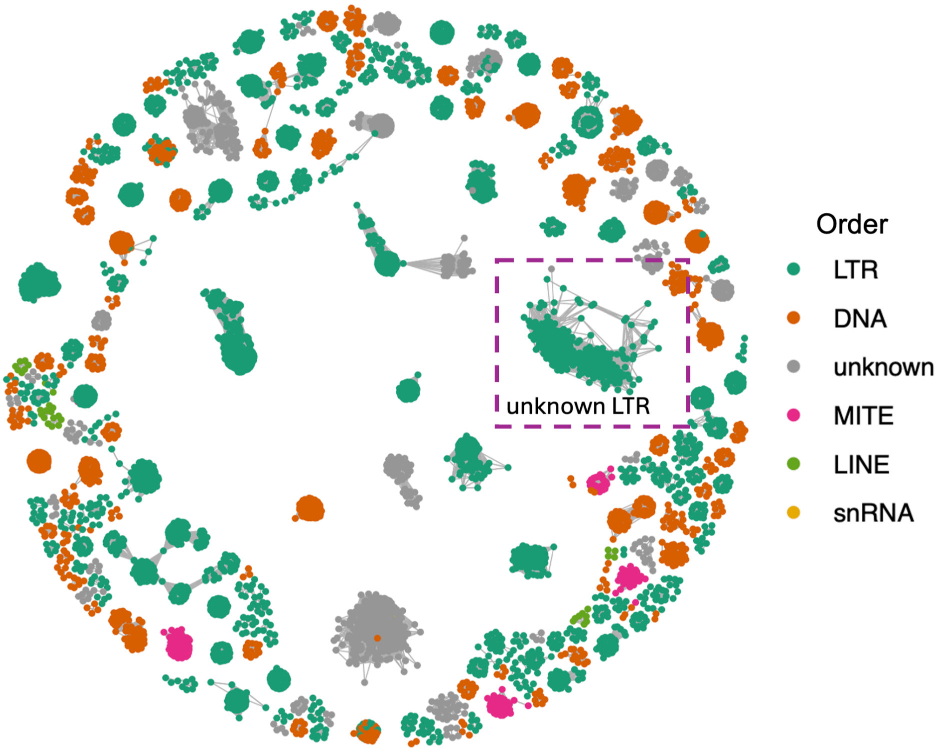

**Supplementary Figure 12. SVs were associated with TEs forming clusters that suggest a common source of origin.** SV association with TEs identified by 85% similarity, at 85% coverage. TE-associated SVs colored by TE orders according to *de novo* annotations.

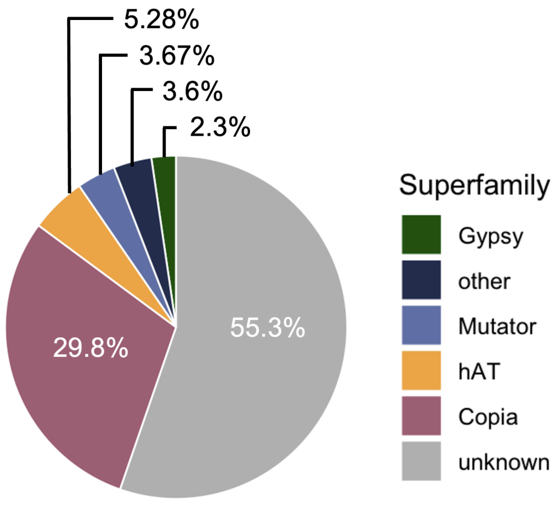

**Supplementary Figure 13. Most TE-associated SVs are from unknown superfamilies.** Shown by the proportions of TE superfamilies in TE-associated SVs.

a

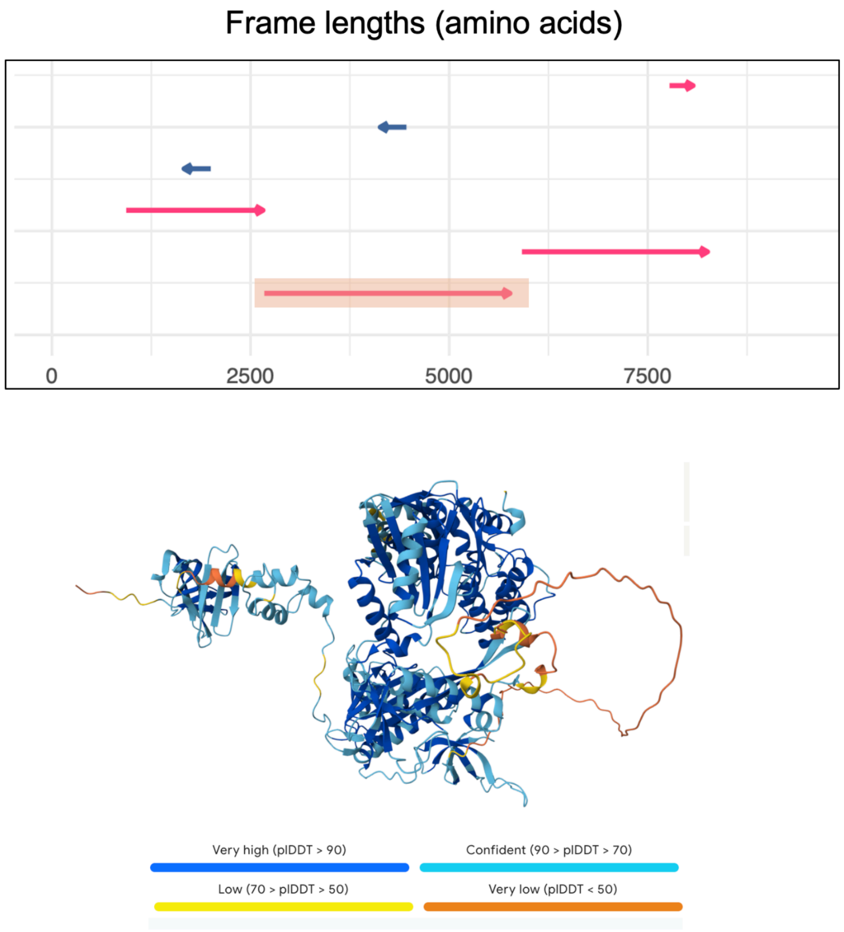

b

**Peptide sequence:**

MQVSVKKKVKWYLDSGCSRHMTGDSNCFIKIVQVNGGKVSFGGNSKGSIVGFGTVKIGNLTISNVSLVEGLNYNLLSISQLCDTG
FKISFQEGTCSGISKDSTQSFMGRRHGNIYLLDVKPNEAQCLISIQDEAVLWHRKLGHVNMKQLAKISSKKLVRGLPKLPYQKSDS
CTPCILGKQVRNSFKQINHVSTNHVLQLLHMDLFGPTRTQSIGGKKYCLVIVDDYSRFTWVYFLSSKSETFSYFEKFAKKVQNEK
GCAISSIRTDHGSEFENKDFTKFCDKSGFNHMFSSPYTPEQNGVVERKNRSLQEMARTLLIESKISSRFWAEAVSTACYIINRVFL
RPILEKTPYELFKGKEPIVSYFHVFGCKCFILKNANDRVGKFEERSDEGIFLGYSTSSKAYRVYNKKSQLVEESTNVKFQDSMQDE
SSQTQYEESEPTPDLPMLTTQEASQSLEVQHQEAHEDSNKERDHQLGRTSSNWKHKSSHPKDLIIGEMNEGIRTRSKRHEESS
AEALISEIEPKSIEEALSDESWIEAMQEELRQFNINDVWELTSKPKGKTVIGTKWVFRNKMNEEGKVVRNKARLVAKGYTQEEGID
YDETYAPVARLEAIRLLLAFACYKNFRLFQMDVKSAFLNGFINEEVYVEQPPGFEDPKKQHSVFRLKKALYGLKQAPRAWYDRLS
KFLLQNGFVKGKVDTTLFIKKESKSFLLVQIYVDDIIFGSSNECLCEKFSKSMQDEFEMSMMGELTFFLGLQIKQMKEGTFIHQEKY
ANDLVKRFGLKNCKKTDIPMSCSQKIDKDEEGKKVDQKLYRSMIGSLLYLTASRPDILLSVCICARFQSDPRESHLSAVKRVIKYVA
SSSSIGLWYPKKGDFKLLGYSDVDLAGCRVDRKSTSGICQILGNRTVSWFSRKQSTVSLSTTEAEYVALGSCCSQILWIKQQLQD
FGIEDSCTEIRCDNTSAINLTKNPILHSRAKHIEIRHHFIRDHIQNGEISIQFVDSKSQLADIFTKPLEKNQFNLIRSRLNILKPEEVKV

**Supplementary Figure 14. Characterisation of a novel LTR in *Eucalyptus*.** (a) Intact open reading frames (ORFs) in the full-length LTR. (b) Alpha fold results of the *Eucalyptus* Retrovirus-related Pol polyprotein.

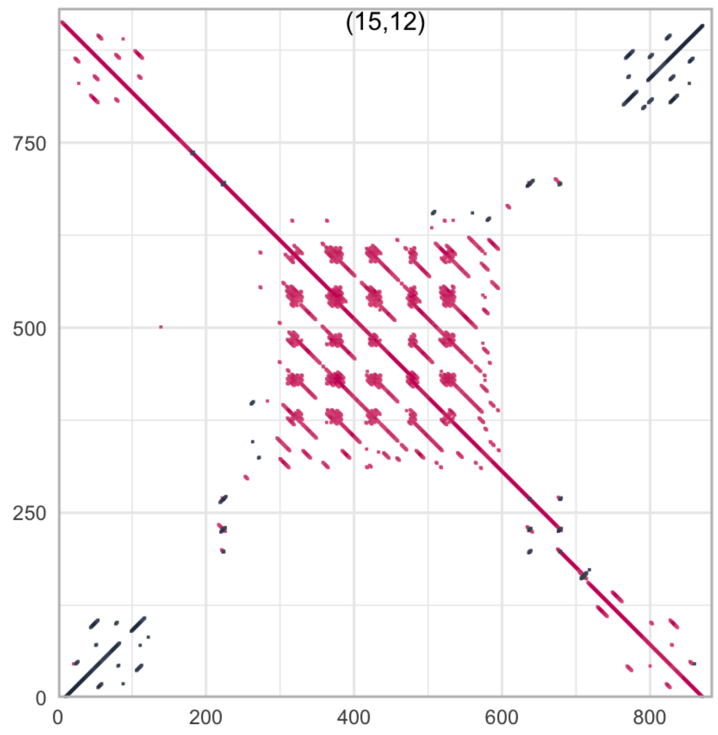

**Supplementary Figure 15. Other TE structures identified from TE-associated SVs.** Dot plot representation of Hiltron elements in TE-associated SVs.

### Note 8: Sniffles2 SV discovery, merging and genotyping parameter tuning

Sequencing coverage is often a source of non-biological variation for SV discovery and genotyping. To ensure that SV discovery and genotyping in this study are not biased by sequence coverage, per-base coverage was calculated for each sample with Mosdepth (version 0.3.3) (Pedersen and Quinlan, 2018) (Supplemental Table 2). Supplemental Table 2 shows that at 12x coverage, 90% of the genome is covered by at least three reads, the minimum number of reads for SV discovery. At 8x coverage, 90% of the genome is covered by at least two reads in all samples (except for MON0575), the preferred number of reads needed for SV genotyping. As such, the thresholds of >12x coverage for discovery and >8x for genotyping were preliminarily determined.

To empirically quantify the effects of sequencing coverage on SV discovery, we performed a down-sample experiment. In doing so, 18 samples with coverage >20x were down-sampled to decreasing levels of coverages (20x, 15x, 10x, 5x, and 3x) with SAMtools (version 1.9) (Danecek et al., 2021). The number of discovered SVs increases rapidly from 3x to 10x, but then slows down after 10x and plateaus by 15x coverage (Fig. S16). Together with the previous per-base depth analyses, the criteria of using samples with >12x coverage for SV discovery and samples with >8x coverage for SV genotyping were well-supported by both empirical evidence and the literature (De Coster et al., 2021).

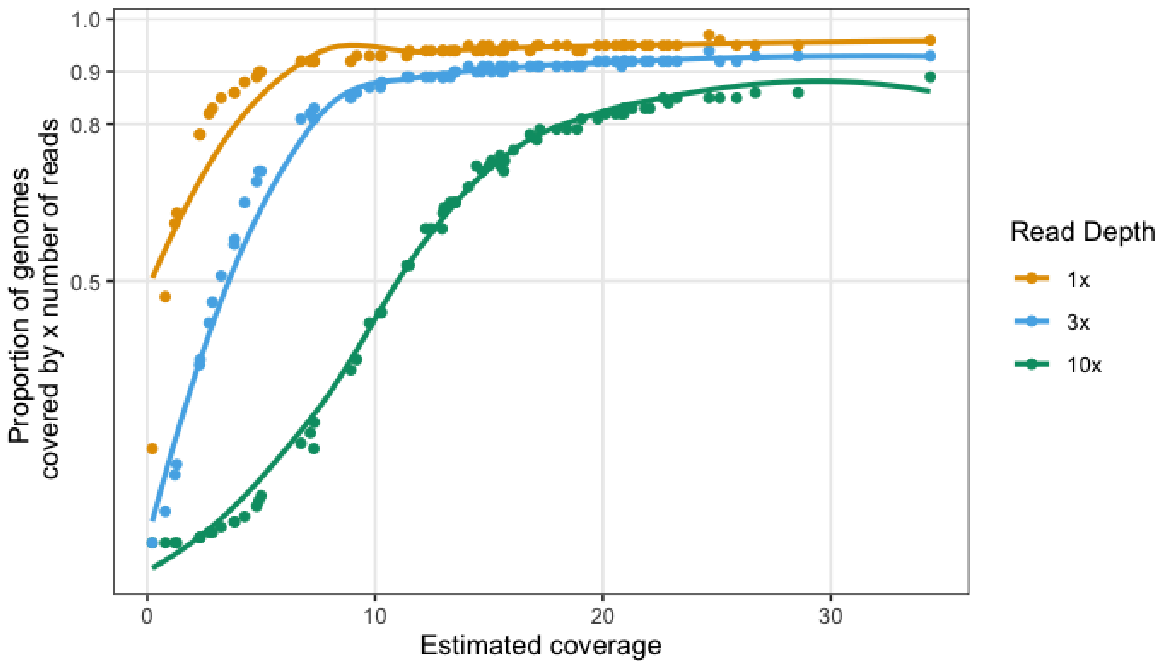

**Supplementary Figure 16. the effects of decreasing levels of coverage on the number of SVs discovered.** Results from the downsample experiment showing the number of insertions and deletions discovered at different  levels of coverages (3x, 5x, 10x, 15x, and 20x) for 18 samples with original coverages over 20x.

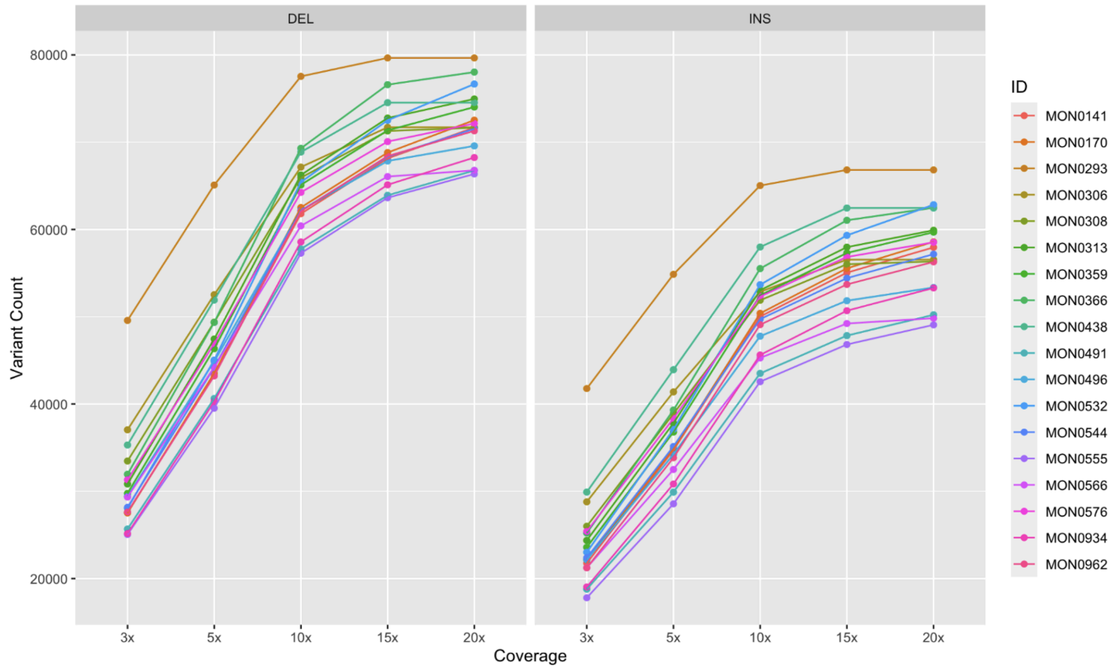

**Supplementary Figure 17. Subsampling experiment reveals robust coverage criteria.** By randomly subsetting high-coverage samples, we identified Sniffles2 discovers consistent number of SVs with higher than 10x genomic coverage using long-read data.

**a**

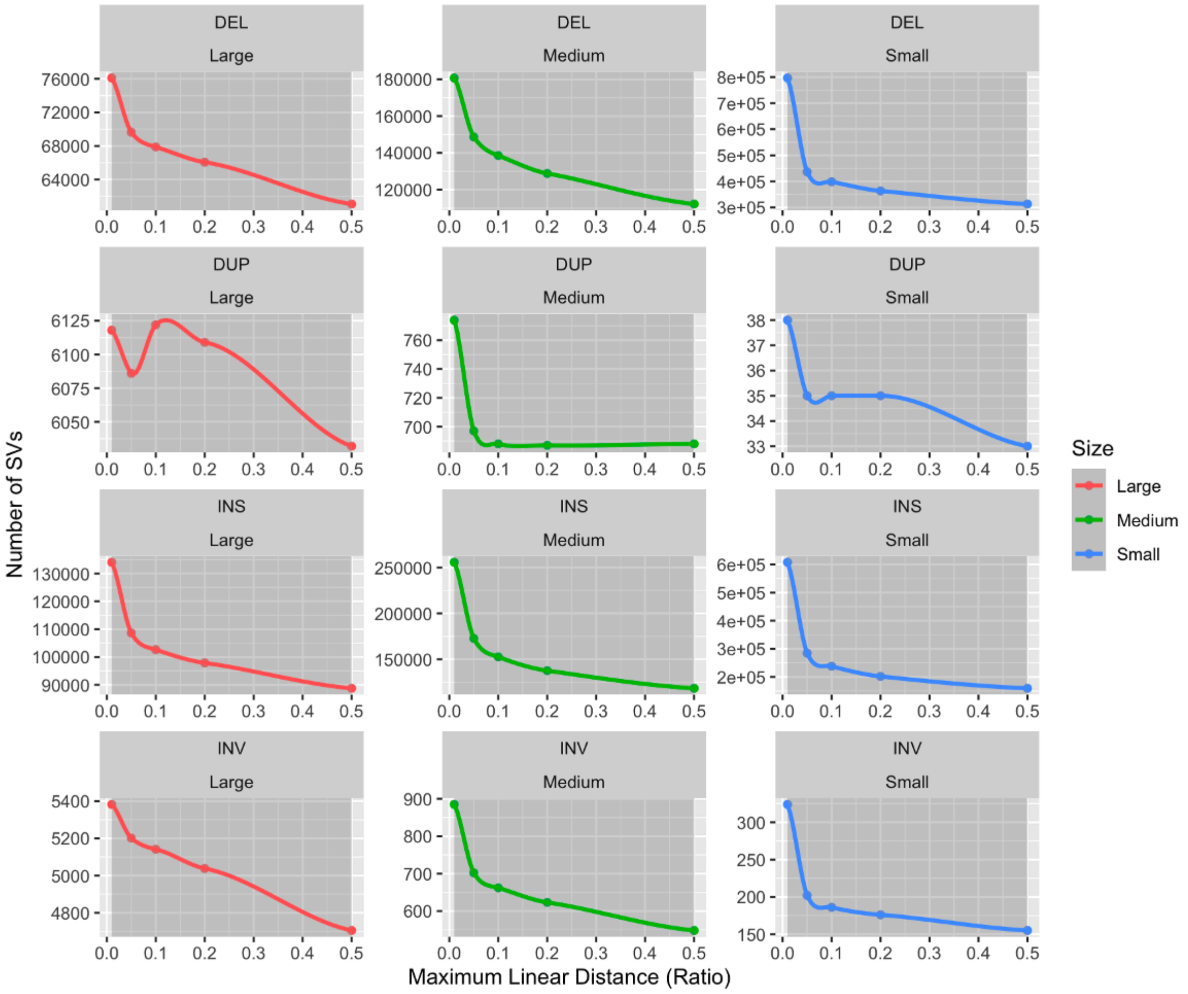

**b**

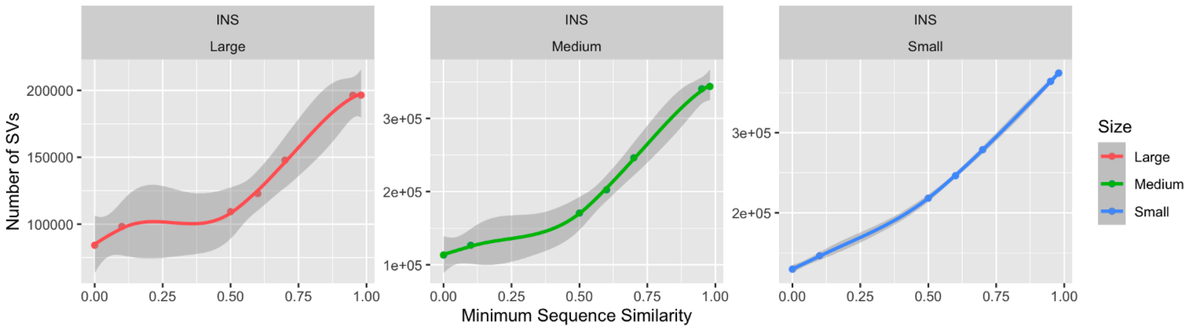

**Supplementary Figure 18. Both breakpoint and sequence identity have consequences in SV merging. (a)** The effects of maximum break point distance on merging capacity. (b) The effects of sequence identity criteria on merging capacity.

### Note 9: Sniffles2 SV discovery and analyses - extended results

In order to investigate if SVs influence certain gene pathways more than others, the annotated gene models were first functionally annotated according to homology of genes in related species. Using Fisher's exact tests, the most enriched gene pathways affected and avoided by SVs were identified.Multiple plant defence related pathways were found significantly enriched ($p < 0.05$) with SVs. These include: disease resistance, disease resistance protein and nucleotide-binding domain, and leucine-rich repeat (NB-LRR), suggesting higher mutation rates of these pathways (mediated by SVs) (Fig. S21). In contrast, many transcription factor related gene pathways were found to be avoided by SVs , indicating lower SV mediated mutations in these pathways (Fig. S21).

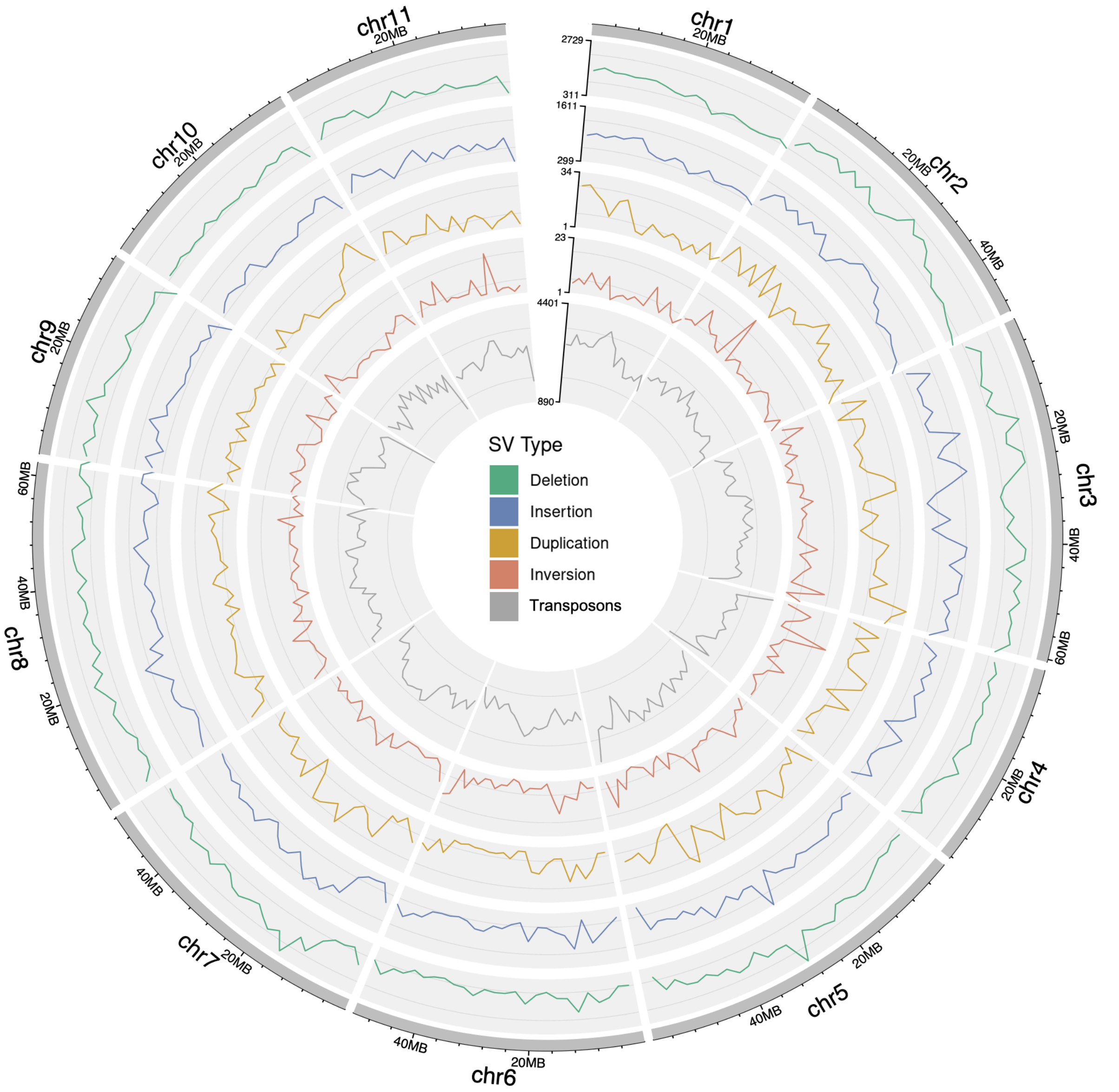

**Supplementary Figure 19. Sniffles2 discovers SVs consistently across the genome.** Using Sniffles2, SVs of different types were discovered evenly all across the genome with some local maxima and minima.

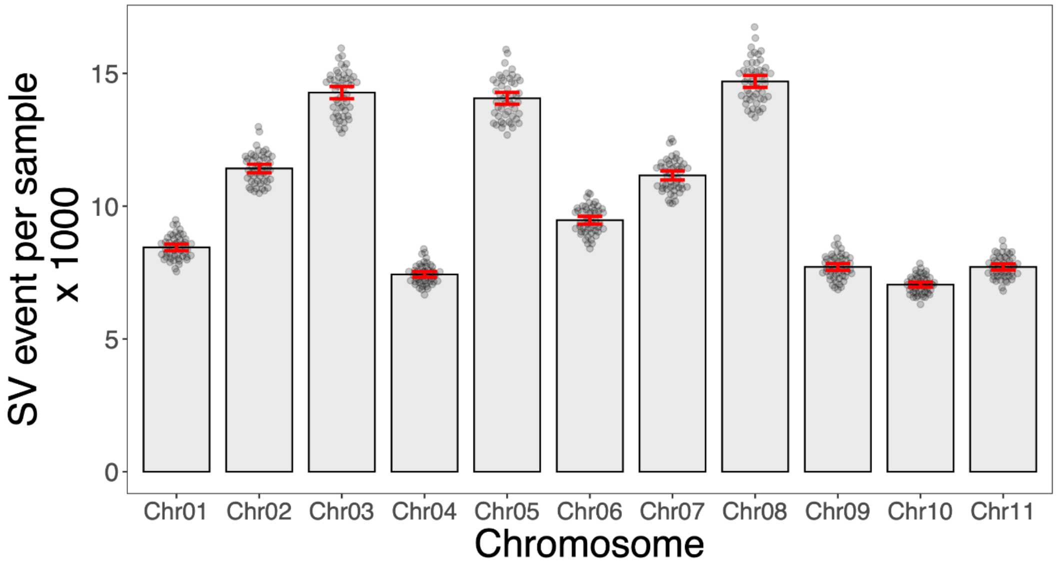

**Supplementary Figure 20. Landscape samples contain similar SV loci.** Across the 11 chromosomes, each chromosome contains a similar number of SVs in each input sample (sd < 200).

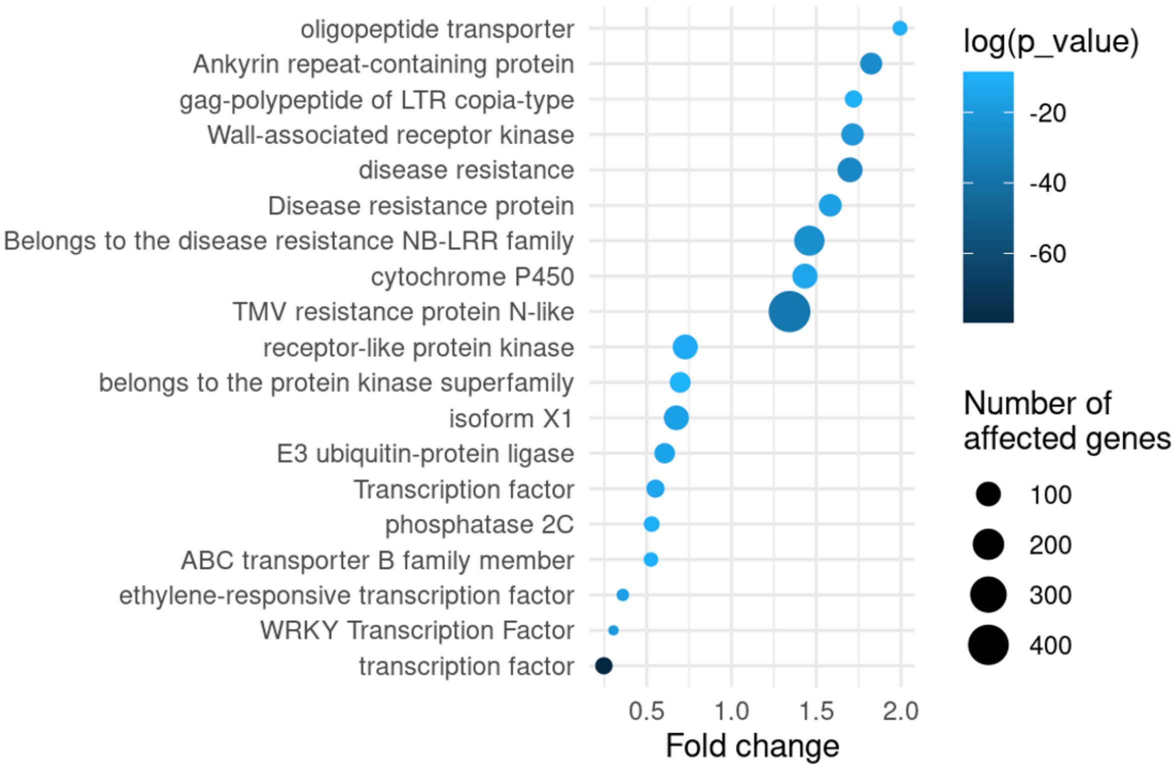

**Supplementary Figure 21. Structural variant enrichments in gene pathways.** (B) The most and least affected gene pathways by structural variations.

### Note 10: SV-based landscape analyses - extended results

Because of multi-colinearity between different BioClim variables, competing for variance, we performed a redundancy analysis for the 49 geo-referenced *E. viminalis* accessions included in the association study. However, BioClim4 had one flat line in one of the leave-one-out scenarios, therefore is not a robust candidate for being tested for climate associations.

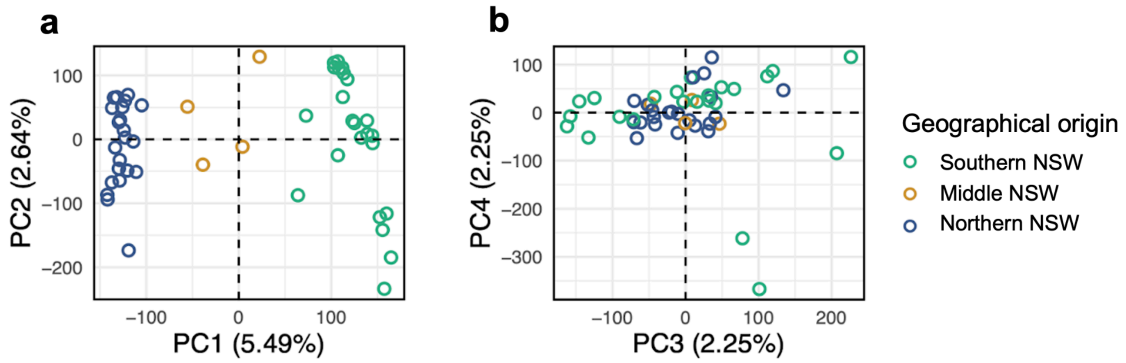

**Figure 22. Spatial distributions of structural variant allele frequencies.** (a) The first two Principal Components (PC1, PC2) of the Principal component analysis (PCA), (b) The third and fourth Principal Components (PC3, PC4).

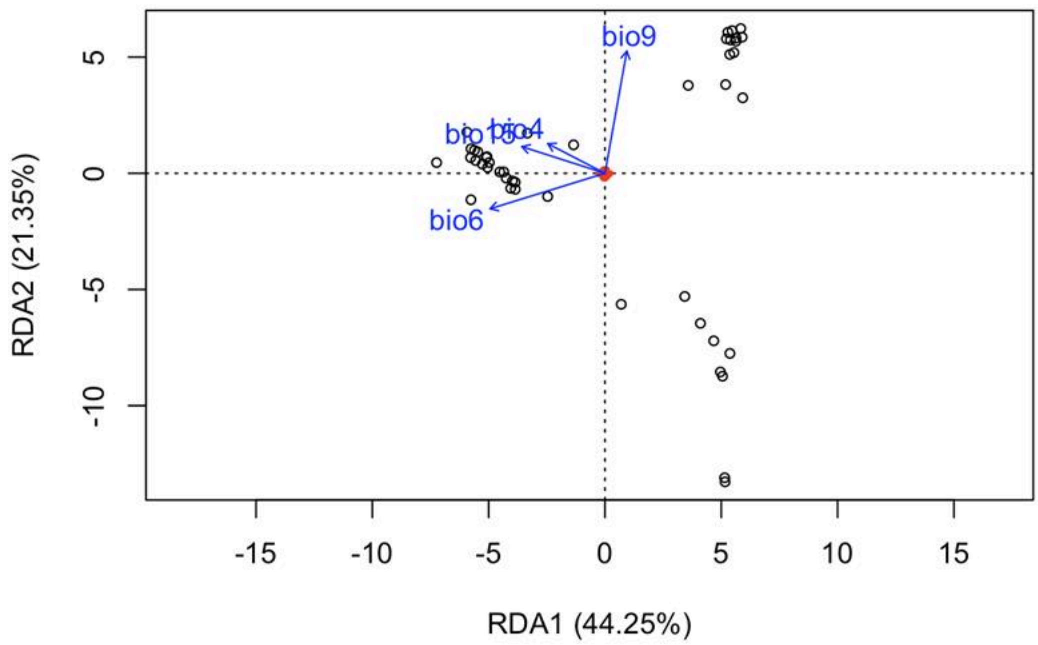

**Supplementary Figure 23. Non-redundant BioClim variables drove genetic clustering.** Due to high multi-colinearity between many BioClim variables, a redundancy analysis (RDA) was performed. Variables with variance inflation factors (VIF) larger than 10 were removed iteratively, to obtain non-redundant drivers of SV-based genetic structure.

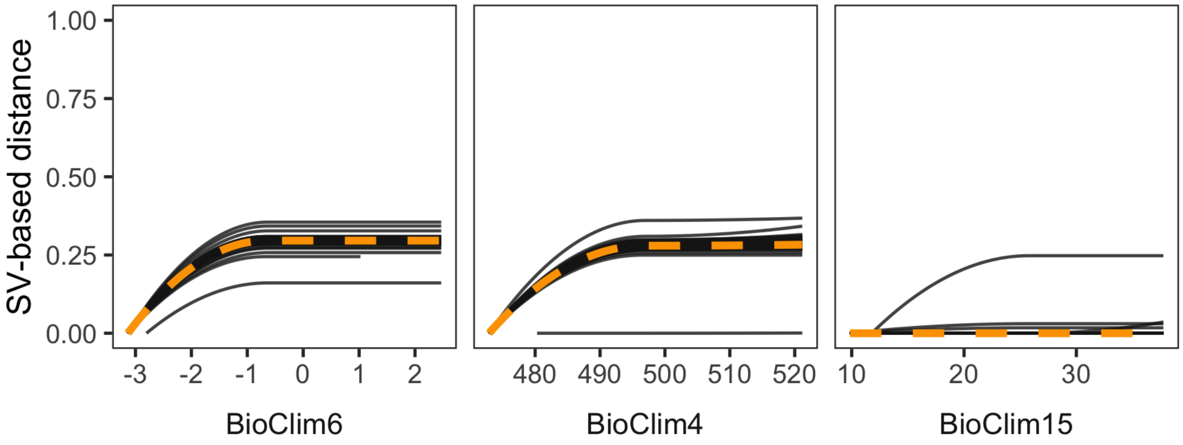

**Supplementary Figure 24. Clinical co-variation between BioClim and genetic differences.** We detect variations in non-redundant BioClim variables defined in the previous plot. BioClim15 was left out due to no genetic covariation, likely absorbed by BioClim4.

### Note 11: Characterisation of candidate genes at *CHILL1*

Through an SV-GWAS, the cold climate adaptation locus *CHILL1* was identified, with structural variants localizing proximal to several genes of interest (Table S11). Two glycosylphosphatidylinositol (GPI)-anchored proteins were located in this locus (g2295.t1 and g2299.t1). The closest *Arabidopsis thaliana* ortholog for both genes was GPI-anchored protein At1G61900, which has been genomically predicted (Borner et al., 2002) and subsequently verified by proteomic analysis (Borner et al., 2003). Although this GPI-anchored protein remains largely uncharacterized, others have been identified to have roles in cold acclimation (Takahashi et al., 2016). In *Arabidopsis*, the GPI-anchored protein At1G61900 is expressed across multiple tissues (Fig.S25), with strong expression in reproductive tissues including flowers, carpels, pollen, and seeds. In our study, we observed minimal to no detectable expression (0 TPM) in leaf tissue using ONT direct RNA sequencing, suggesting potential tissue-specific or environmentally regulated expression patterns that may contribute to cold adaptation.

The 7-hydroxymethyl chlorophyll a reductase gene (g2297.t1), orthologous to *Arabidopsis* AT1G04620, showed measurable expression at 13.42 TPM. This enzyme catalyzes the final step of the chlorophyll cycle by converting 7-hydroxymethyl chlorophyll a to chlorophyll a, a process essential for light acclimation, greening, and senescence (Meguro et al., 2011). Consistent with the expression observed in *E. viminalis* leaf tissue, strong expression in *A. thaliana* leaf and vegetative tissues has also been reported (Fig.S26).

Additionally, two genes encoding α-terpineol synthase proteins were present, both of which appeared to be directly influenced by SVs. The first, α-terpineol synthase (g2294.t1), shares homology with *Arabidopsis* terpene synthase 4 (AT4G16740) and is directly disrupted by the 485 bp *CHILL1.1* insertion within its first intron. The second, (+)-α-terpineol synthase-like gene (g2298.t1) with homology to *Arabidopsis* terpene synthase-like sequence-1,8-cineole (AT3G25820), was in close proximity to *CHILL1.5* (a 323 bp insertion in close the suspected promoter region). Neither α-terpineol synthase gene was detected as expressed in *E. viminalis leaves.* Terpene synthase genes constitute a large and rapidly evolving gene family in *Eucalyptus*, with members influencing multiple aspects of leaf biochemistry including foliar oil content and composition (Külheim et al., 2015). α-Terpineol synthase proteins have not been characterised in cold acclimation, although α-terpineol can regulate important leaf traits such as stomatal closure in tomato (*Solanum lycopersicum*) (Pérez-Pérez et al., 2024). Lastly, an unannotated gene (g2296.t1) was present, with no matches identified in NCBI databases.

**a**

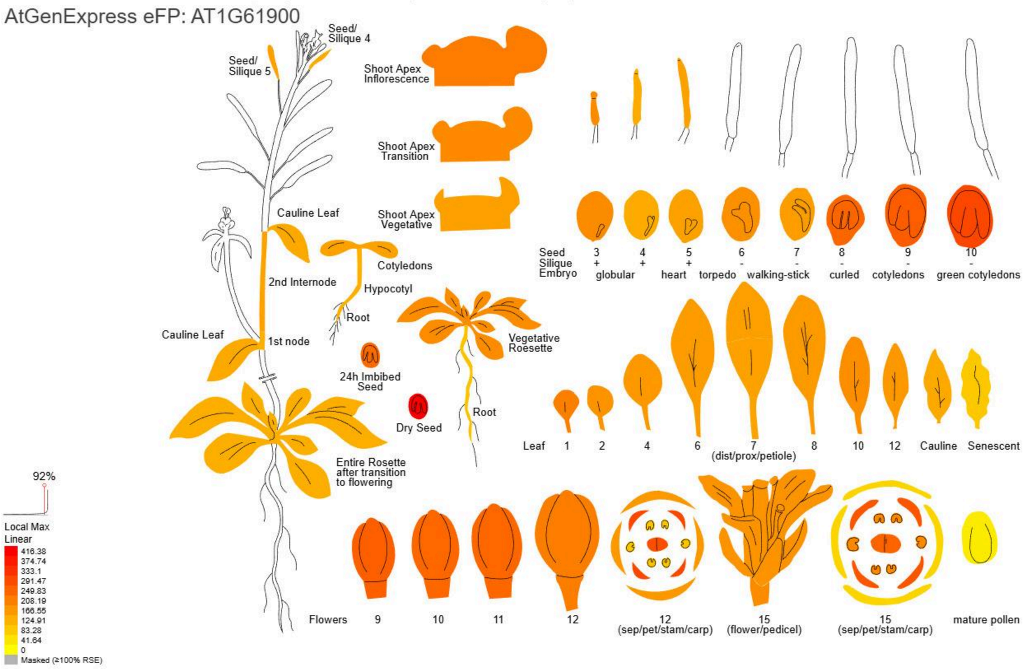

**b**

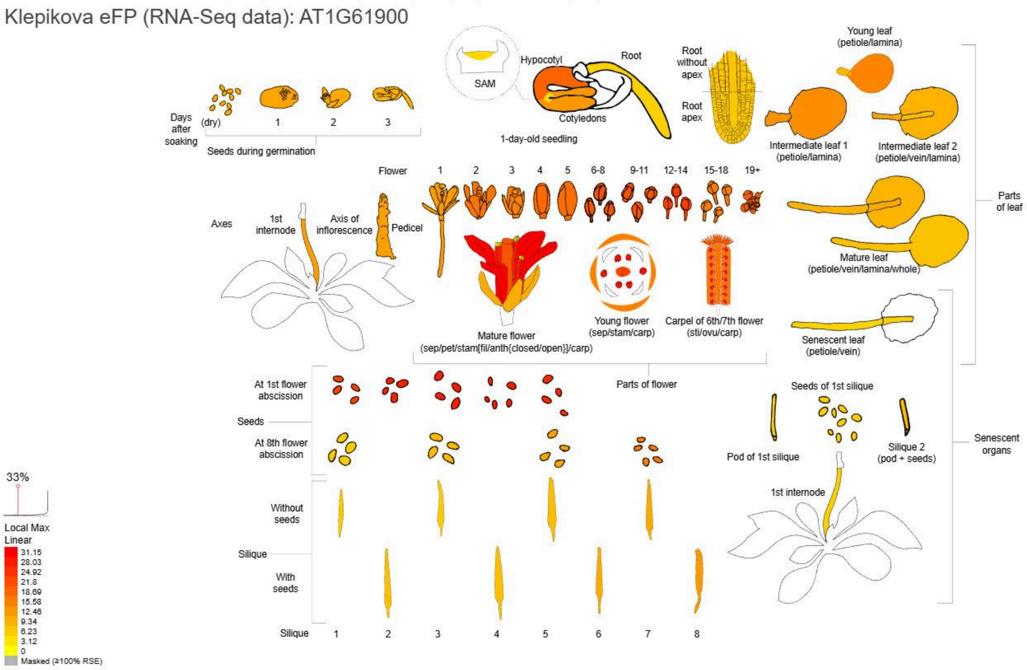

**Supplementary Figure 25. Relative expression level of the *Arabidopsis thaliana* GPI-anchored protein gene At1G61900.** The closest matching Arabidopsis ortholog of *Eucalyptus viminalis* GPI-anchored protein g2295.t1 was identified by a protein BLAST (NCBI) and expression visualized with the tool ePlant (Waese et al., 2017). **(a)** AtGenExpress eFP, data from (Schmid et al., 2005) and (Nakabayashi et al., 2005) **(b)** Klepikova eFP (RNA-Seq data), data from (Klepikova et al., 2016).

**a**

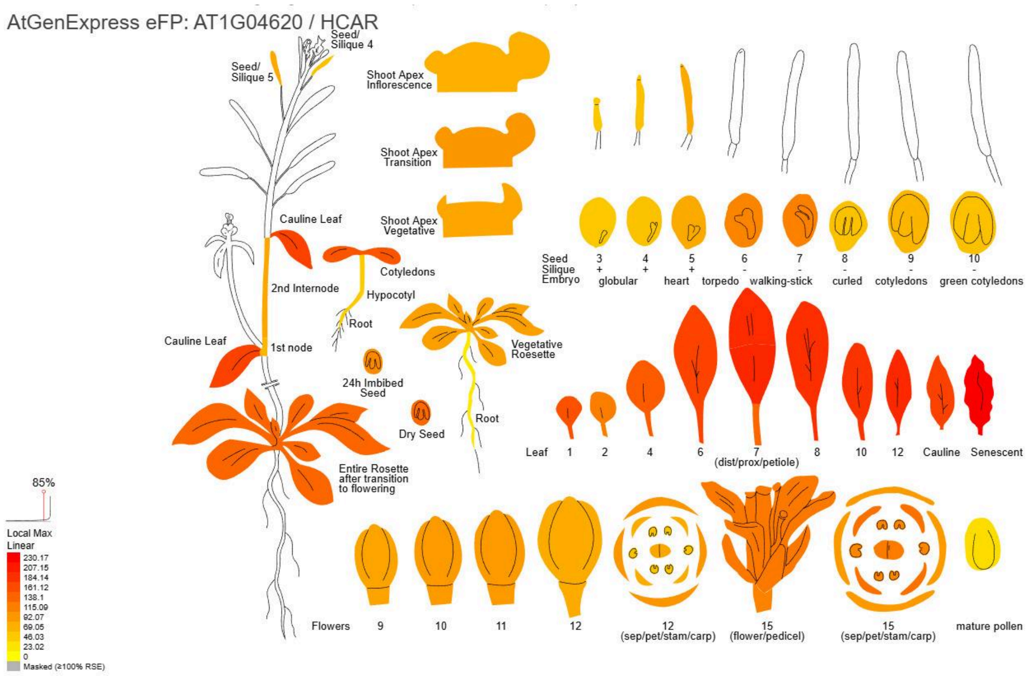

**b**

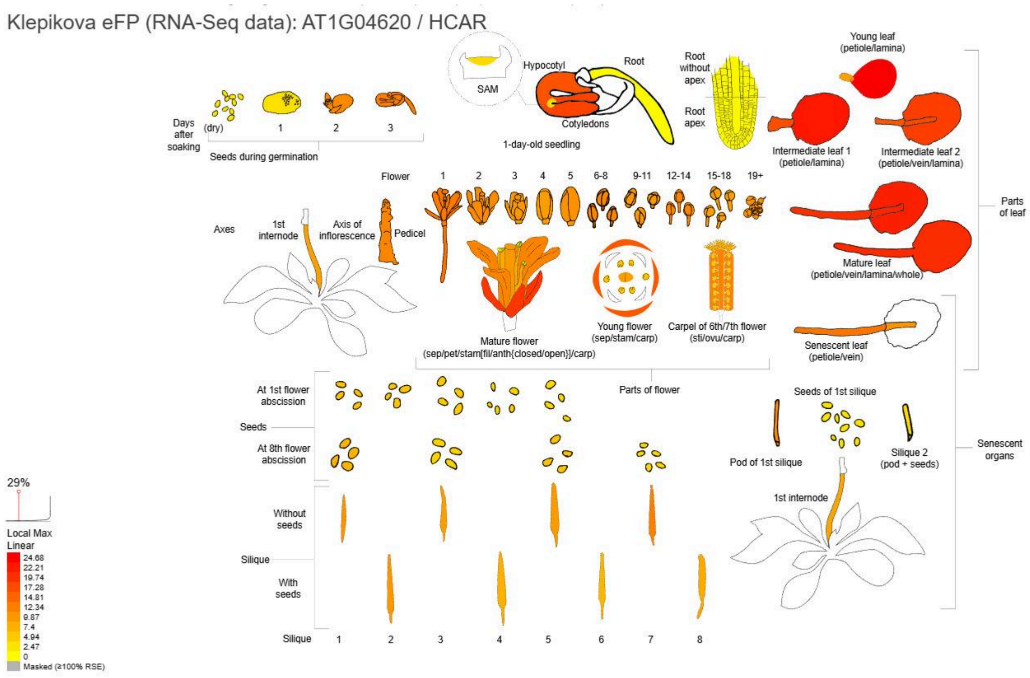

**Supplementary Figure 26. Relative expression level of the *Arabidopsis thaliana* 7-hydroxymethyl chlorophyll a reductase gene AT1G04620.** The closest matching Arabidopsis ortholog of *Eucalyptus viminalis* gene g2297.t1 was identified by a protein BLAST (NCBI) and expression visualized with the tool ePlant (Waese et al., 2017). **(a)** AtGenExpress eFP, data from (Schmid et al., 2005) and (Nakabayashi et al., 2005) **(b)** Klepikova eFP (RNA-Seq data), data from (Klepikova et al., 2016).

#### Note 12: Characterisation of candidate genes at *CHILL8*

Through an SV-GWAS, we identified 10 SVs across four loci conferring climate adaptation (Fig. S27). Nine SVs conferred cold adaptation, while *CHILL-8.1* conferred adaptation to warming temperatures. At *CHILL-8.1*, SVs localized proximal to a phosphate transporter *PHT1;3* (g34497.t1) and a serine/threonine-protein phosphatase *PP1* (g34498.t1) (summarised in Table S12). Phosphorus (P) is essential for plant growth, with most terrestrial plants acquiring phosphorus as inorganic orthophosphate (Pi) from soil (Wang et al., 2025). This adaptation is particularly relevant given that Australian soils exhibit characteristically low total and available P (Eldridge et al., 2018), with P limitation intensifying under tropical climates such as Queensland Australia (Hou et al., 2020). Climate warming further exacerbates these constraints by accelerating soil P depletion and reducing bioavailability, thereby threatening forest ecosystems (Guo et al., 2024; Tian et al., 2023). Microbial competition for P additionally constrains acquisition by *Eucalyptus* species in these P-limited systems (Jiang et al., 2024).

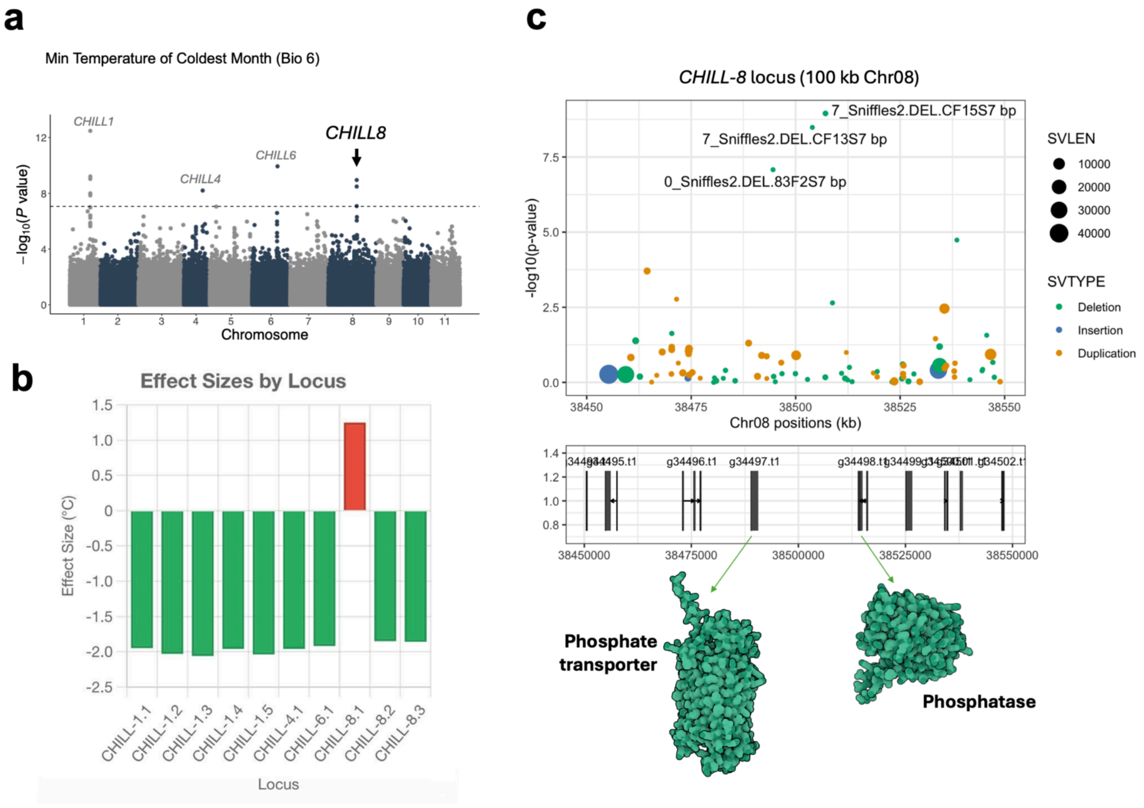

**Supplementary Figure 27. *CHILL8* relating to warm rather than cold adaptation.** (a) Manhattan plot of *CHILL1* and *CHILL8*. (b) Effect size comparison with other *CHILL* loci. (c) Locus zoom-in of *CHILL8* located on chromosome 8.

*E. viminalis EvPHT1;3* showed leaf expression (1,476 TPM). The closest *Arabidopsis thaliana* ortholog *AtPHT1;1* (AT5G43350) exhibited 97% query coverage and 80.50% identity (protein BLAST score 869), with expression enriched in root epidermis plasma membranes (Fig.S28). The *AtPHT1* family comprises high-affinity Pi transporters upregulated under phosphate starvation (Bayle et al., 2011), with overexpression enhancing growth under P limitation (Mitsukawa et al., 1997). Enhanced P uptake elevates root biomass and regulates leaf transpiration to maintain ecosystem productivity (Hou et al., 2020). Functionally, *AtPHT1* transporters interact with plasma membrane intrinsic protein (PIP) aquaporins PIP1;2 and PIP2;1 to co-ordinately regulate water transport (Bellati et al., 2016).

The upstream location of SVs relative to *EvPHT1;3* suggests altered transcriptional regulation. In *A. thaliana*, transcription factors WRKY45 (Wang et al., 2014) and PHR1 (Rubio et al., 2001) bind the *AtPHT1;1* promoter to activate expression under phosphate starvation. SVs may modulate these regulatory interactions, potentially enhancing Pi acquisition capacity under warming-induced P limitation.

The co-localized serine/threonine-protein phosphatase PP1 isozyme 3 isoform X phosphatase (40 TPM) was found to be orthologous to the *Arabidopsis thaliana* type one serine/threonine protein phosphatase 4 (TOPP4, AT2G39840). TOPP proteins are highly expressed in multiple tissues (Fig S29) and may have diverse roles in post-translational regulation of proteins. For instance, phosphatases may further augment P scavenging. Under nutrient starvation, such as warming conditions, the essential degradation process autophagy occurs, which is primarily regulated by autophagy-related (ATG) proteins. TOPP proteins dephosphorylate ATG proteins which initiate autophagy under nutrient starvation (Wang et al., 2022). Alternatively, phosphatase proteins may help influence tissue development in response to climate. In *Arabidopsis*, TOPP4 dephosphorylates PIN-FORMED 1 (PIN1) which regulates interdigitated pavement cell morphogenesis in the leaf (Guo et al., 2015). SVs altering expression levels could provide adaptive capacity while maintaining essential function.

**a**

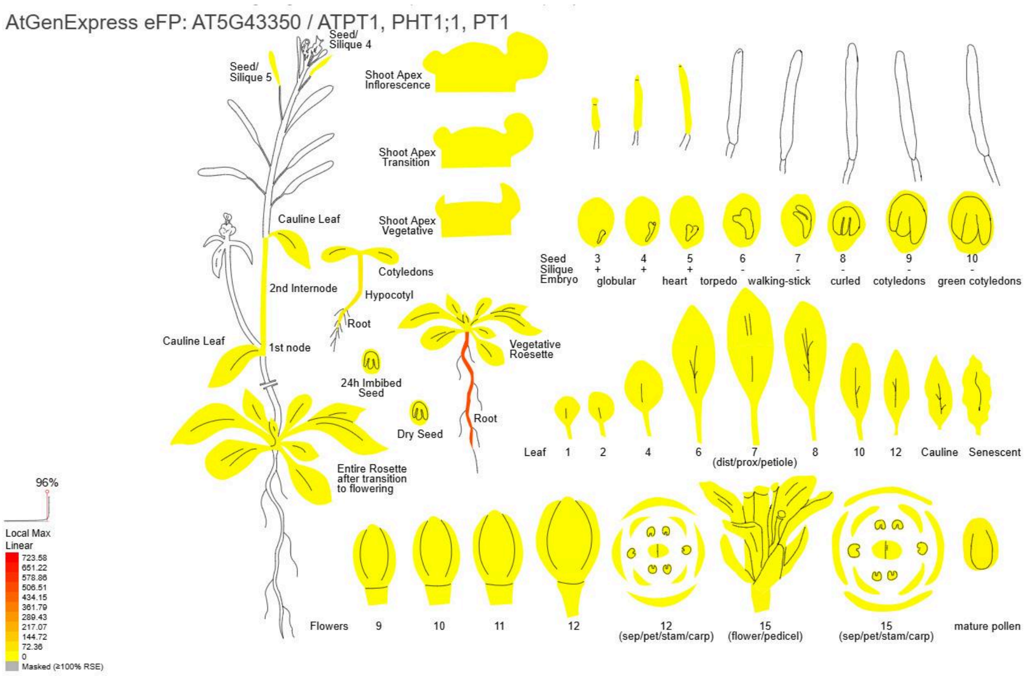

**b**

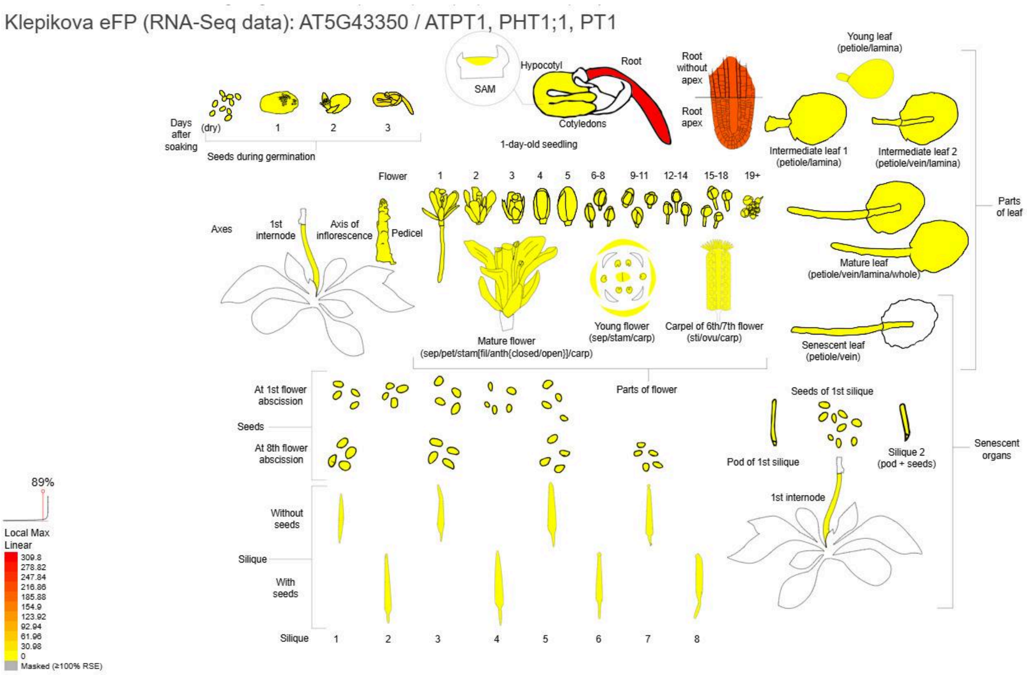

**Supplementary Figure 28. Relative expression level of the *Arabidopsis thaliana* phosphate transporter PHT1;1 gene AT5G43350.** The closest matching Arabidopsis ortholog of *Eucalyptus viminalis* inorganic phosphate transporter 1-3 was identified by a protein BLAST (NCBI) and expression visualized with the tool ePlant (Waese et al., 2017). **(a)** AtGenExpress eFP, data from (Schmid et al., 2005) and (Nakabayashi et al., 2005) **(b)** Klepikova eFP (RNA-Seq data), data from (Klepikova et al., 2016).

**a**

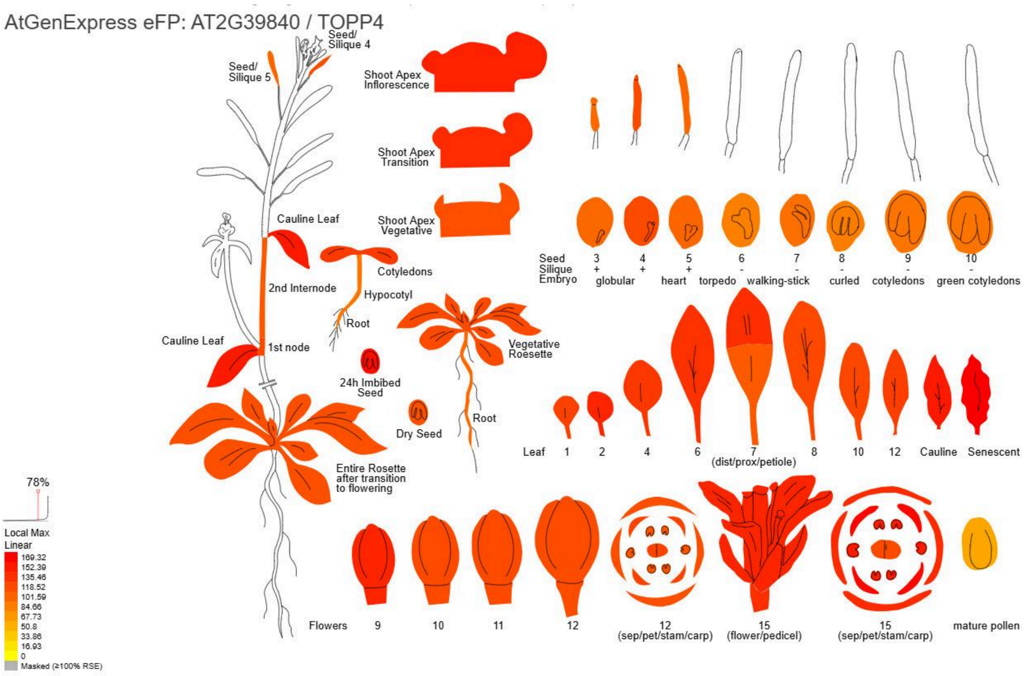

**b**

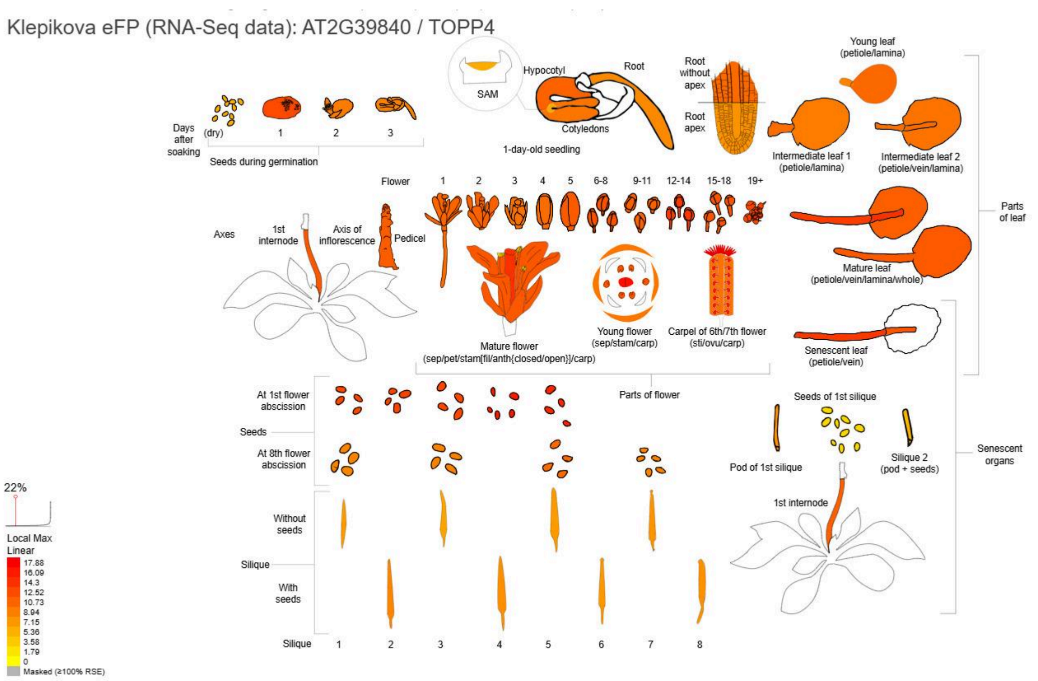

**Supplementary Figure 29. Relative expression level of the *Arabidopsis thaliana* type one serine/threonine protein phosphatase 4 TOPP4 gene AT2G39840.** The closest matching Arabidopsis ortholog of *Eucalyptus viminalis* serine/threonine-protein phosphatase PP1 isozyme 3 isoform X1 was identified by a protein BLAST (NCBI) and expression visualized with the tool ePlant (Waese et al., 2017). **(a)** AtGenExpress eFP, data from (Schmid et al., 2005) and (Nakabayashi et al., 2005) **(b)** Klepikova eFP (RNA-Seq data), data from (Klepikova et al., 2016).

### Note 13: Population genomics with short-read dataset

#### 13.1 Methods

To validate the *CHILL1* locus, the original set of 49 *E. viminalis* samples with long-read data was expanded to include a total of 434 samples of *E. viminalis* (N = 176) and two closely related species, *E. dalrympleana* (N = 138) and *E. rubida* (N = 120) (Supplementary Table 16; Fig. S30b) which were then whole genome sequenced using short-read technology. SNPs were called across species then filtered for quality, minor allele frequency and missingness (see Methods in main paper).

To examine species differentiation and geographic population structure within the 434 samples, principal component analyses (PCA) were applied to data from 247,409 bi-allelic pre-filtered SNPs randomly selected from across the genome at a minimum spacing of 2,000 bp (median of 2,035 bp) using the pcaMethods package in R (https://www.bioconductor.org/packages/release/bioc/html/pcaMethods.html). The same SNP data were used to explore for evidence of recent hybridisation between species using the ADMIXTURE program (https://dalexander.github.io/admixture/).

#### 13.2 Results

The first two principal components based on genome-wide SNP data broadly delineated the three species (*E. viminalis*, *E. rubida* and *E. dalrympleana*) from similar species range (Fig. S30a), explaining 9.5% of the variance (Fig. S30c). Species labels based on morphological observations in the field (leaf, trunk and fruit broad characteristics), and subsequently cross-checked in the laboratory against field records to validate, were notably discrepant to species labels defined by SNP data (12.4% discordance). However, there was no obvious bias from geographic location or species in this discordance. For subsequent analyses, we assigned species identity to samples using SNP-based PC scores rather than morphology-based identification (Fig. S30c).

In addition to species as the primary, albeit weak, driver of genetic structure, latitude also influenced the top four principal components which, together, represent 13% of the total genetic variance (data not shown). Consistent with this, Admixture analyses revealed clear latitudinal structuring in ancestry proportions within each species (Fig. S31). Interestingly, samples exhibiting mixed ancestry tended to draw disproportionately from geographically proximate subpopulations; for example, northern *E. dalrympleana* individuals with *E. rubida* ancestry showed higher contributions from northern *E. rubida* populations than from southern ones (Fig. S31). This spatially structured admixture pattern suggests that interspecific gene flow occurs predominantly at regional scales, rather than uniformly across species ranges. Taken together, genome-wide SNP analyses indicate that the three species exhibit low overall genetic divergence, similar patterns of isolation by distance (latitude) within species, and ongoing hybridisation.

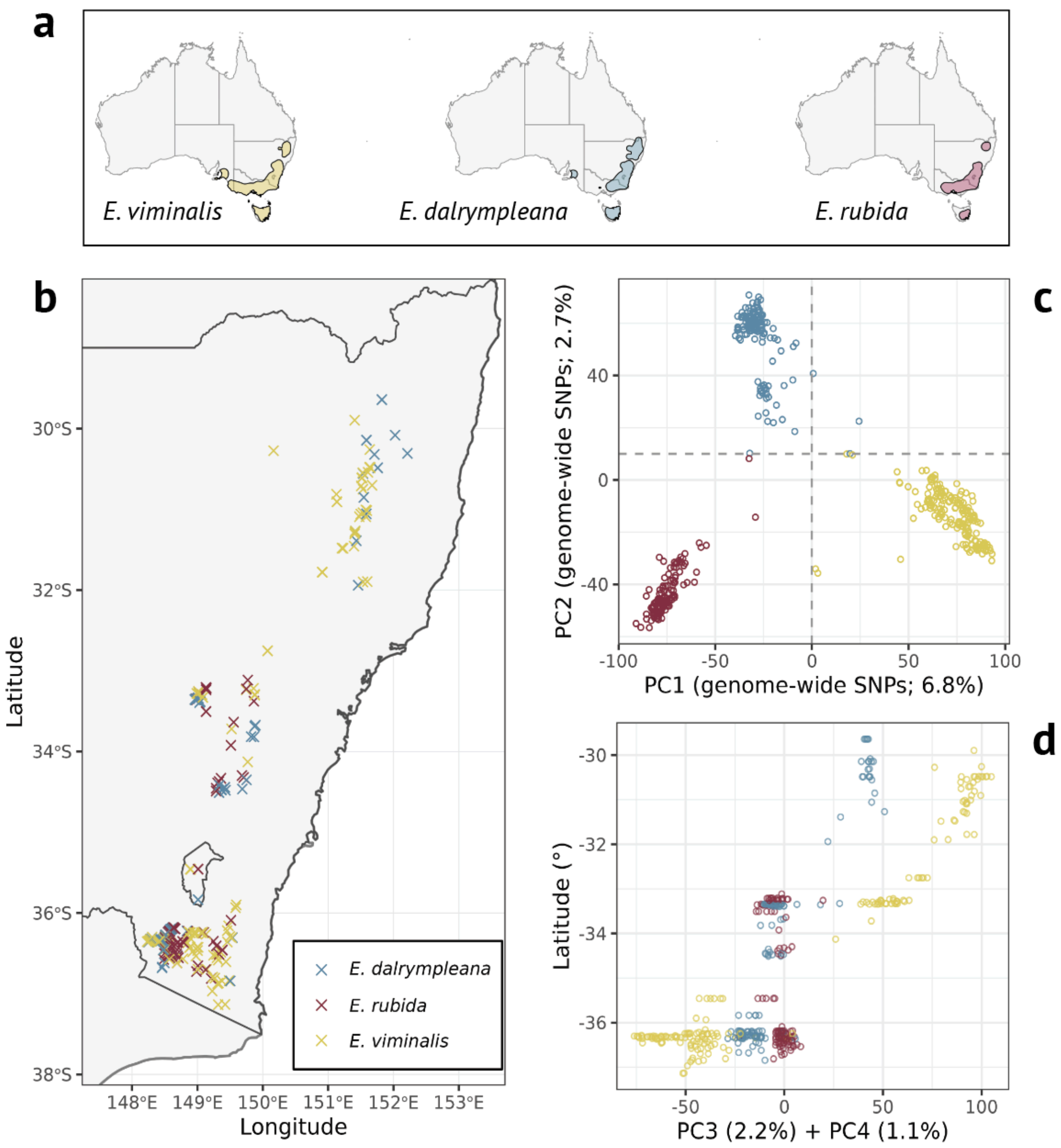

**Supplementary Figure 30. Genome-wide patterns of SNP variation among three closely related species. a.** Geographic distributions of focal species approximated using density kernels based on occurrence records from Atlas of Living Australia, as shown in Fig. 5. **b.** Locations of sample collection points (N = 434) in Southeastern Australia. **b.** Genome-wide SNP variation reflected by the top two PCs; dashed lines show thresholds used to assign species identity, over-riding morphology-based classification in 12.4% of samples. **c**. Sample latitude mapped onto the top two PCs. **d**. Latitude plotted against the sum of the third and fourth PCs.

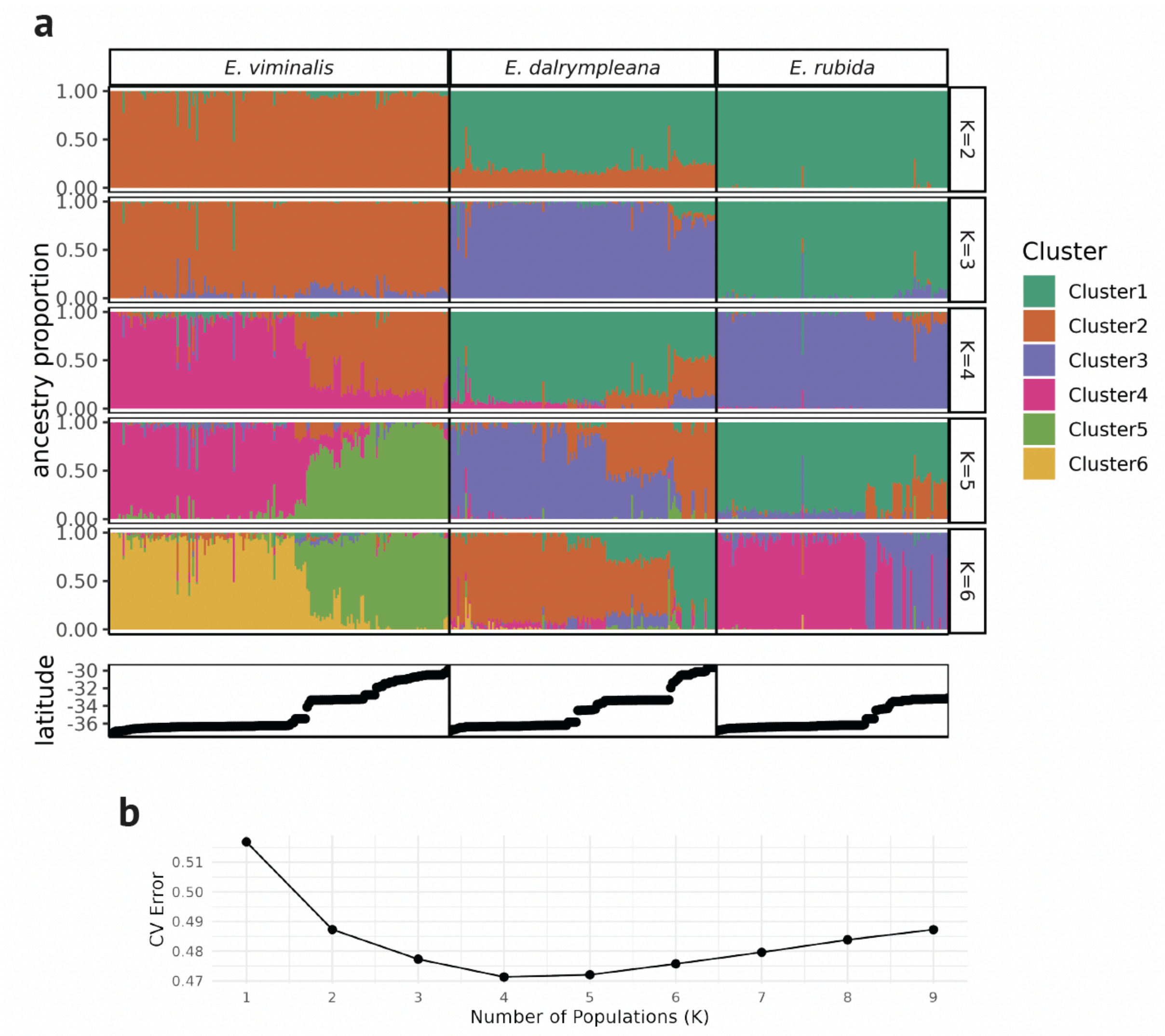

**Supplementary Figure 31. a.** Admixture results using genome-wide SNPs from short-read sequencing of *E. dalrympleana, E. rubida, and E. viminalis* samples, showing estimated ancestry proportions with runs using K=2-6 clusters. Samples in all panels are arranged left-to-right by latitude nested within species. **b.** Cross validation (CV) error scores from Admixture runs from K=1-9.

### Note 14 Short-read and statistical validation of *CHILL1*

#### 14.1 Methods

Since short-read data could not directly genotype SVs, we applied principal component analysis to data on SNPs falling within the 0.1Mbp region that bounds the five *CHILL1* SVs (positions 27,696,962 to 27,799,070, Fig. 4c, N =1,605). After delineating the three distinct haplotypes (*CHILL1/CHILL1*, *CHILL1/chill1*, *chill1/chill1*) based on the first principal component (PC1), we compared these short-read derived haplotypes with those from long-read SV data in the 46 samples that had both types of data.

To validate the relationship found between BioClim6 and *CHILL1* discovered from GWAS applied to SV data (Fig. 4), we fitted a generalised linear model to *CHILL1* haplotype data (coded as 0, 0.5, and 1 for *CHILL1* genotypes *CHILL1/CHILL1*, *CHILL1/chill1* and *chill1/chill1*, respectively) from the larger sample SNP set (n = 434) with fixed effects, in isolation or in combination, for BioClim6, species, latitude, elevation and PCs derived from genome-wide SNPs. The latter act as proxies for background genetic structure and thus perform a similar role to the kinship matrix used in the GEMMA analysis of the SV data on 49 *E. viminalis* (see main text). Models were fitted using the glm() function from the stats package in R using  the 'family=binomial' option to account for the binary nature of the dependent variable (https://stat.ethz.ch/R-manual/R-devel/library/stats/html/00Index.html).

The magnitude of the effect of *CHILL1* on frost risk, and of other variables was estimated by analysing BioClim6 as the dependent variable in linear models fitting fixed effects for latitude (at 2.5 m resolution) and elevation (at 3 s resolution) as linear covariates, *CHILL1* genotype (a factor with three levels) and species (a factor with 3 levels) using the glm() function from the stats package in R (https://stat.ethz.ch/R-manual/R-devel/library/stats/html/00Index.html).  Each explanatory variable was included in the model separately and then simultaneously, including terms for two-way interactions. Choice of final models for presentation were based on the proportion of variance explained by the model, their relative performance and patterns of residuals using the performance package in R (https://rdrr.io/cran/performance/man/performance-package.html).

#### 14.2 Results

Genetic structuring in the *CHILL1* region starkly contrasted to the genome-wide pattern. This was at first evident from the large (400 kbp) block of tightly linked SNPs surrounding the *CHILL1*- associated SVs (Fig. 5b). It was further evident from the fact that the first PC based on SNPs within the *CHILL1* locus explained 65% of the variation (Fig S32c). Individuals clustered according to three *CHILL1* genotype states and these clusters were unrelated to species groups (Fig. S32c). This pattern was concordant between SNPs and SVs from short- and long-read sequencing, respectively, for the subset of *E. viminalis* samples sequenced with both approaches, for all but two samples (Fig S32a). Unsurprisingly, SVs associated with BioClim6 within *CHILL1* were in strong linkage disequilibrium ($R^2$: 0.88-0.97; Fig. S32b). The strong structure around the *CHILL1* region allowed us to use thresholds on SNP PCA axis 1 to assign *CHILL1* genotype states for all samples subjected to short-read sequencing (Fig S32c; and see Fig 5b) and thereby conduct onward analyses between *CHILL1* and BioClim6 in the larger data set. A key finding of this analysis was that *CHILL1* is polymorphic in the three species analysed here, with haplotypic variation of a single 'supergene' allele being more pronounced than variation within species (Fig. S33d).

The significant relationship between *CHILL1* and BioClim6 found within the *E. viminalis* long-read data set was upheld in the short-read dataset. When *CHILL1* haplotype was analysed as the dependent variable in a linear model fitting BioClim6 and background genetic structure (species or, alternatively, the first two PCs that captured genomic background) with highly significant effects ($P<10^{-11}$; Table S11). BioClim6 explained 14% of the variation in *CHILL1* and background genetics explained a further 7-9%. The relatively small amount of variation explained by background genetics, and the independence of the *CHILL1* effect from background genetics indicate that the association between *CHILL1* and BioClim6 was unlikely to be driven by confounding effects between geography and genome-wide population structure. By contrast, when latitude and elevation or their proxies, PCs 3 and 4, were introduced to the above model, a further large amount of variation in *CHILL1* was explained (25%) and the BioClim6 effect was eliminated (Supplementary Table 15). This reflects that BioClim6, latitude and elevation were highly correlated in our data set and therefore substituted for each other as predictors of *CHILL1*.

In order to understand the relative contributions of *CHILL1*, geography (latitude and elevation) and genetic background (species and genome-wide PCs) to the relationship between *CHILL1* and BioClim6, BioClim6 was analysed as the dependent variable in linear models fitting these terms in isolation or in combination. It was found that geography (latitude and elevation) together explained 72.5% of its variation (Supplementary Table 15; Fig. S33), *CHILL1*, on its own explained 15.8% of the variation and species or background genetics explained around 1% (Supplementary Table 15). As for the models above where the dependent variable was *CHILL1*, geographic variables, but not background genetics, traded with *CHILL1* in explaining variation in BioClim6. This covariation between environmental and geographic variables is almost unavoidable in landscape genomic studies and, indeed, is the source of their power.  By ruling out confounding between background genetics and environment as the driver of the relationship between BioClim6 and *CHILL1* we are confident that the strong cline in allele frequency across this environmental gradient reflects historical strong selection for cold tolerance or related traits.

The above models allowed us to estimate the effect size of *CHILL1* on cold tolerance. It was found that the impact of having two vs. zero copies of *CHILL1* would be equivalent to translocating it from a BioClim6 zone of  -2ºC to 0ºC; that is, in effect, transforming it from frost-susceptible to frost-resistant (Supplementary Table 15). This effect size is equivalent to a range extension from latitude -36$^0$ to  latitude -32$^0$ (equivalent to 444 km) or from 1200 m to 600 m elevation (Supplementary Table 15). Thus, effectively, *CHILL1* confers adaptation to the species' full climatic range.

In conclusion, analysis of SNP data from a much larger data set (n = 434) across three species confirms the result from SV data in *E. viminalis* (n = 49). It strongly supports the hypothesis that *CHILL1* confers frost tolerance and has therefore been strongly selected for in the colder southern and high elevation areas and selected against in warmer northern and lower altitude areas. The adaptive value of *CHILL1* to cold tolerance transcends that of species, challenging the commonly held assumption that species drive environment-specific adaptation. Since *CHILL1* appears to have arisen prior to separation of the three species in this study, we propose that this mutation drove expansion of its common ancestor into a much wider climatic and geographic range, after which new species formed. The alternative hypothesis — that *CHILL1* arose in one species, then through introgression via species hybridisation and strong selection (relatively common in *Eucalyptus*) (McLay et al., 2023; Myburg et al., 2004), became established in the other two species - is also plausible.

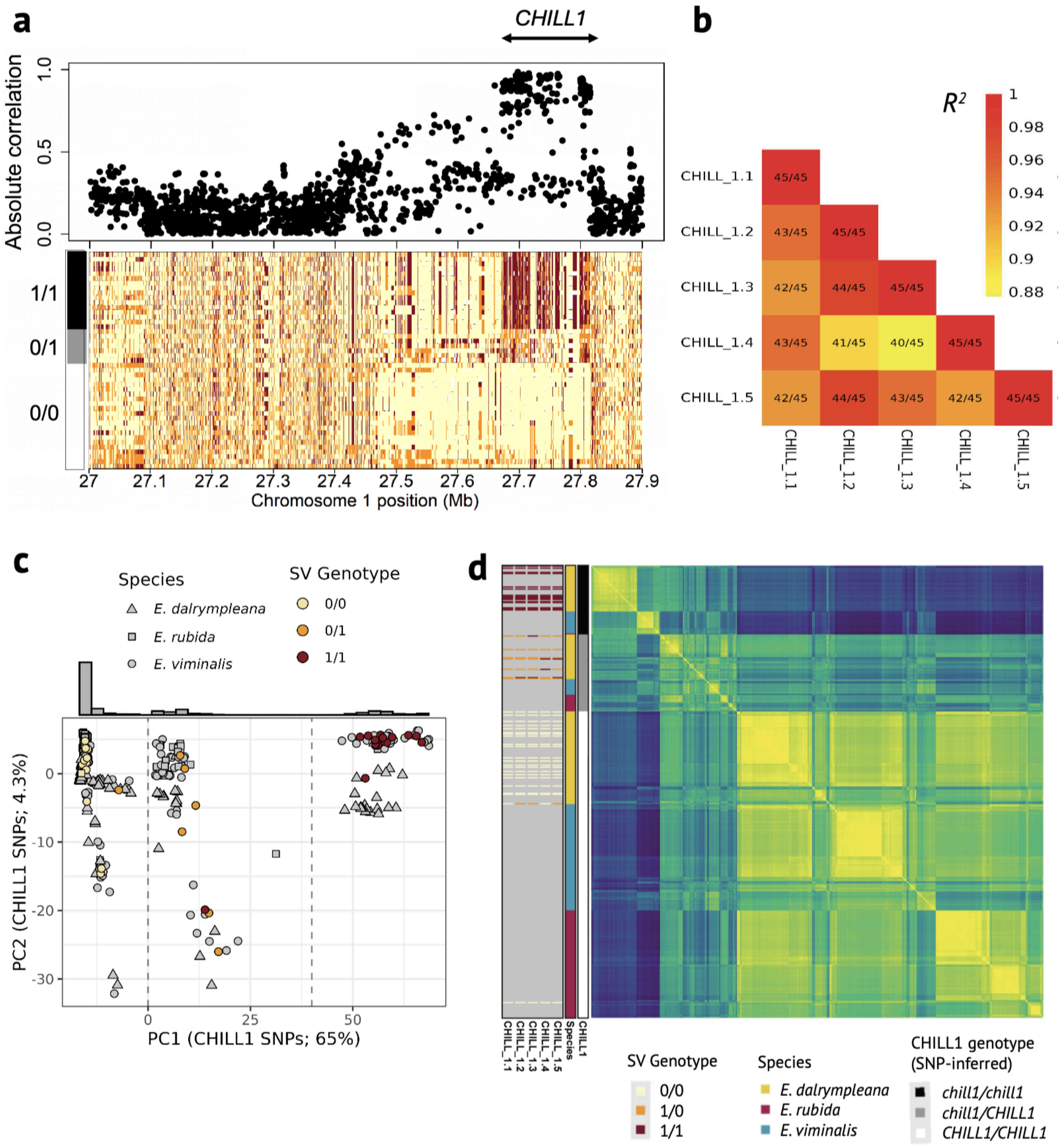

**Supplementary Figure 32. a.** Correlation between the SV *CHILL1.2* and SNPs, for samples treated with long- and short-read sequencing, shows extensive linkage at the CHILL locus that is evident as haplotype blocks in the SNP matrix for those samples. **b.** Pairwise linkage disequilibrium ($R^2$) and genotype concordance (fractions) among the top-5 GWAS hits with BioClim6 of SVs within *CHILL1*. **c.** The first two principal components from SNPs within the *CHILL1* locus (Chr1 27,696,962 to 27,799,070 bp) explain, respectively, 65% and 4.3% of variance. Samples that were sequenced with long-reads (N = 45, main text) are coloured to represent the average SV genotype from the top-5 GWAS hits with BioClim6 within *CHILL1*. Vertical lines represent PC1 thresholds used to assign *CHILL1* genotypes for short-read samples shown as grey; histogram represents the distribution of PC1 scores **c.** Heatmap showing pairwise genetic similarity (high, yellow; low, blue) based on SNPs within the wider *CHILL1* locus (Chr1 25 to 28 Mbp) among 434 samples. Samples are sorted by SNP-based *CHILL1* genotype then species then hierarchical clustering within these.

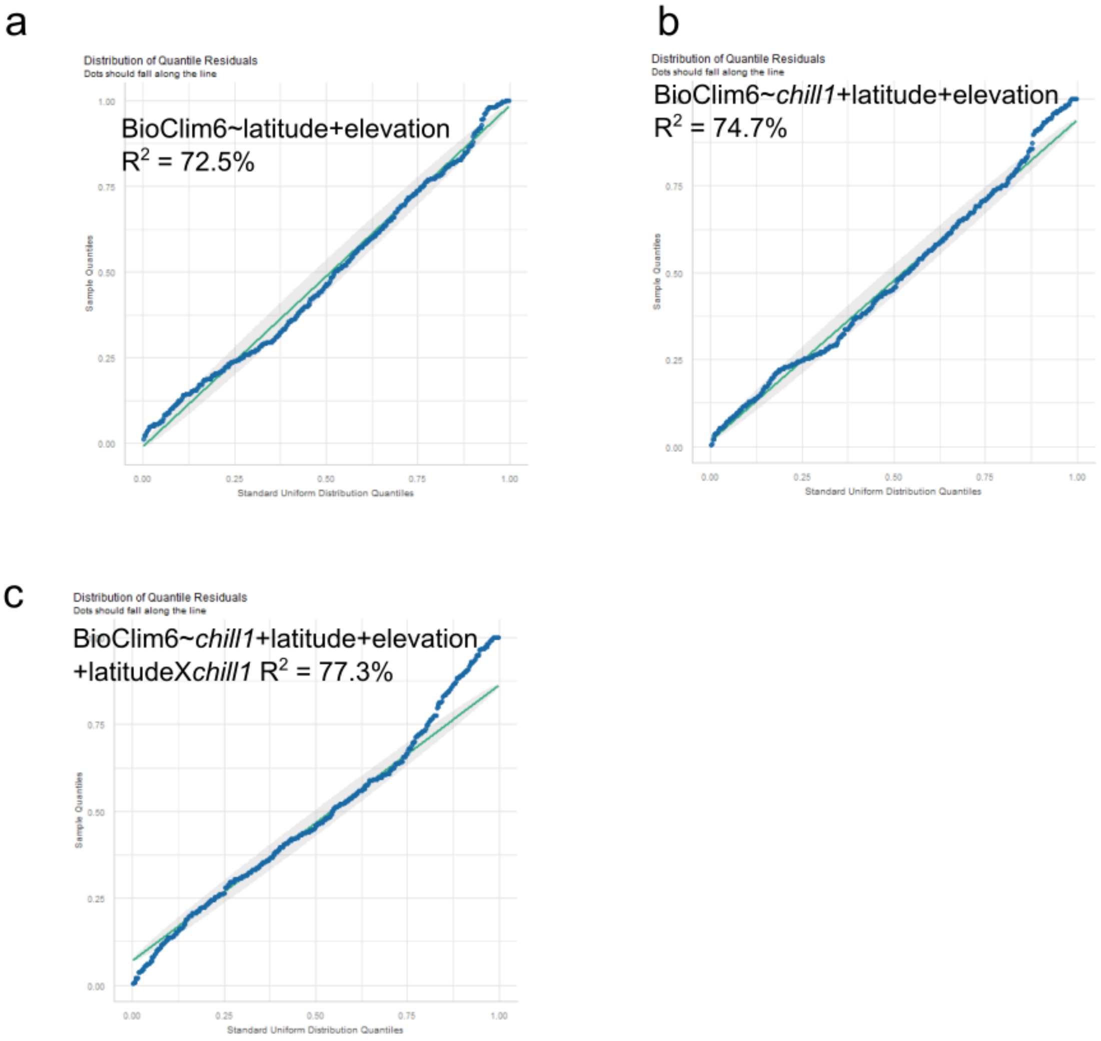

**Supplementary Figure 33. Model fits of BioClim6 to geographic variables (latitude and elevation) and *CHILL1*.** **a**. BioClim6 is well explained by fitting both latitude and elevation as linear covariates. **b** *CHILL1* explains substantial variation in BioClim6 over and above that explained by latitude and elevation but generates a poor fit at the upper end of the latitude spectrum **c**. Due to non-linear relationships between *CHILL1* and latitude, modelled as a two-way interaction, improves overall fit but leads to a strong negative bias to the effect size of *CHILL1* in models that include latitude and its interactions with *CHILL1* (Table S15).

### Note 15: Further characterisation of *CHILL1* - extended results

#### Methods

To examine variation among haplotypes around *CHILL1*, we used both multidimensional dotplots and PGGB. First, to select homologous regions of interest, we used the synteny blocks of GENESPACE to find the best match to *CHILL1* in each of 9 alternative haplotypes (see main Methods). Specifically, we align synteny-resolved orthogroup orders between each haplotype, finding the best match between these haplotype sequences in terms of GENESPACE orthologs. We do this at an orthogroup level for higher sensitivity in the presence of rampant complex structural variants that disrupt sequence-level synteny, in a manner similar to the syntenic neighbourhood definition in Teasdale et al 2025. Then, we use the PGGB pipeline to construct both all-by-all alignments with WFMash, and subsequently a graphical sequence pangenome of the *CHILL1* supergene. All-by-all alignments are plotted as dotplots using custom R code that utilises pafR, and sequence graphs are plotted using odgi vis.

#### Results

Pairwise alignment of 9 haplotypes at the *CHILL1* locus further reinforces the broadly syntenic nature of both *CHILL1* haplotypes (Fig. S34). While there are many large singleton insertions/deletions, overall synteny is preserved both within and between *CHILL1* haplotypes. Several large insertion/deletion SVs are individual-specific. Limited long-read sampling of *chill1/chill1* individuals prohibits detailed examination of the broader structural differences between alternative *CHILL1* haplotypes identified by alignment to a reference (see main manuscript Fig. 5-6)

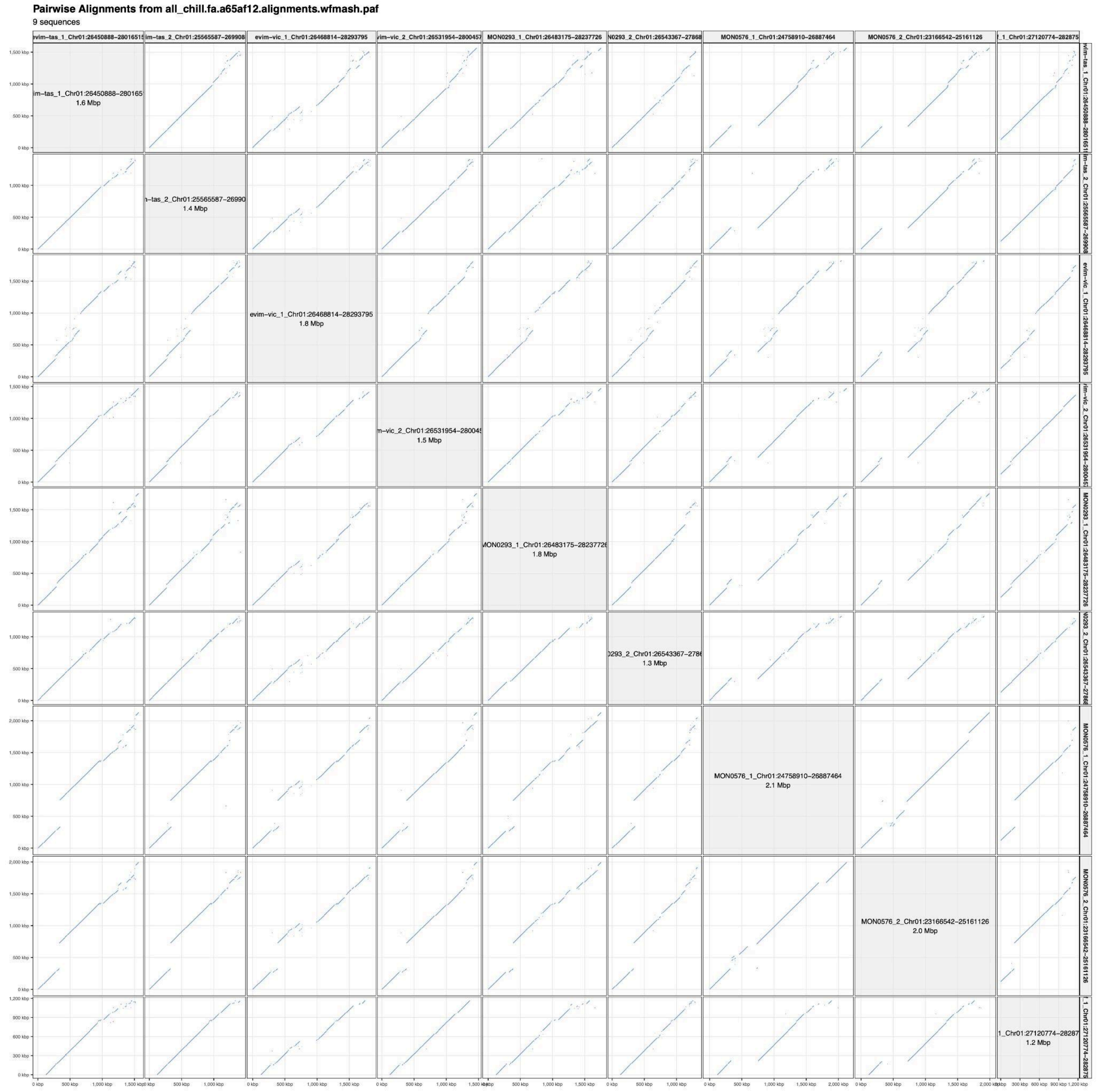

**Supplementary Figure 34. *CHILL1* lacks major inversions and is largely syntenic.** *CHILL1* region defined by shared synteny-resolved orthogroups identified with GENESPACE across 9 haplotype-resolved genomes show the *CHILL1* region is broadly syntenic despite some largely individual-specific complex structural variation.

Liao, W.-W., Asri, M., Ebler, J., Doerr, D., Haukness, M., Hickey, G., Lu, S., Lucas, J.K., Monlong, J., Abel, H.J., Buonaiuto, S., Chang, X.H., Cheng, H., Chu, J., Colonna, V., Eizenga, J.M., Feng, X., Fischer, C., Fulton, R.S., Garg, S., Groza, C., Guarracino, A., Harvey, W.T., Heumos, S., Howe, K., Jain, M., Lu, T.-Y., Markello, C., Martin, F.J., Mitchell, M.W., Munson, K.M., Mwaniki, M.N., Novak, A.M., Olsen, H.E., Pesout, T.,

Porubsky, D., Prins, P., Sibbesen, J.A., Sirén, J., Tomlinson, C., Villani, F., Vollger, M.R., Antonacci-Fulton, L.L., Baid, G., Baker, C.A., Belyaeva, A., Billis, K., Carroll, A., Chang, P.-C., Cody, S., Cook, D.E., Cook-Deegan, R.M., Cornejo, O.E., Diekhans, M., Ebert, P., Fairley, S., Fedrigo, O., Felsenfeld, A.L., Formenti, G., Frankish, A., Gao, Y., Garrison, N.A., Giron, C.G., Green, R.E., Haggerty, L., Hoekzema, K., Hourlier, T., Ji, H.P., Kenny, E.E., Koenig, B.A., Kolesnikov, A., Korbel, J.O., Kordosky, J., Koren, S., Lee, H., Lewis, A.P., Magalhães, H., Marco-Sola, S., Marijon, P., McCartney, A., McDaniel, J., Mountcastle, J., Nattestad, M., Nurk, S., Olson, N.D., Popejoy, A.B., Puiu, D., Rautiainen, M., Regier, A.A., Rhie, A., Sacco, S., Sanders, A.D., Schneider, V.A., Schultz, B.I., Shafin, K., Smith, M.W., Sofia, H.J., Abou Tayoun, A.N., Thibaud-Nissen, F., Tricomi, F.F., Wagner, J., Walenz, B., Wood, J.M.D., Zimin, A.V., Bourque, G., Chaisson, M.J.P., Flicek, P., Phillippy, A.M., Zook, J.M., Eichler, E.E., Haussler, D., Wang, T., Jarvis, E.D., Miga, K.H., Garrison, E., Marschall, T., Hall, I.M., Li, H., Paten, B., 2023. A draft human pangenome reference. Nature 617, 312–324. https://doi.org/10.1038/s41586-023-05896-x

Marcon, H., Domingues, D., Marino, C., 2011. Identification of potential transcriptionally active Copia LTR retrotransposons in Eucalyptus. Presented at the BMC Proceedings, Springer, p. P164.

Marcon, H.S., Domingues, D.S., Silva, J.C., Borges, R.J., Matioli, F.F., de Mattos Fontes, M.R., Marino, C.L., 2015. Transcriptionally active LTR retrotransposons in Eucalyptus genus are differentially expressed and insertionally polymorphic. BMC Plant Biol. 15, 198.

McLay, T.G., Fowler, R.M., Fahey, P.S., Murphy, D.J., Udovicic, F., Cantrill, D.J., Bayly, M.J., 2023. Phylogenomics reveals extreme gene tree discordance in a lineage of dominant trees: hybridization, introgression, and incomplete lineage sorting blur deep evolutionary relationships despite clear species groupings in Eucalyptus subgenus Eudesmia. Mol. Phylogenet. Evol. 187, 107869.

Meguro, M., Ito, H., Takabayashi, A., Tanaka, R., Tanaka, A., 2011. Identification of the 7-hydroxymethyl chlorophyll a reductase of the chlorophyll cycle in Arabidopsis. Plant Cell 23, 3442–3453.

Mitsukawa, N., Okumura, S., Shirano, Y., Sato, S., Kato, T., Harashima, S., Shibata, D., 1997. Overexpression of an Arabidopsis thaliana high-affinity phosphate transporter gene in tobacco cultured cells enhances cell growth under phosphate-limited conditions. Proc. Natl. Acad. Sci. 94, 7098–7102.

Myburg, A.A., Vogl, C., Griffin, A.R., Sederoff, R.R., Whetten, R.W., 2004. Genetics of postzygotic isolation in Eucalyptus: whole-genome analysis of barriers to introgression in a wide interspecific cross of Eucalyptus grandis and E. globulus. Genetics 166, 1405–1418.

Nakabayashi, K., Okamoto, M., Koshiba, T., Kamiya, Y., Nambara, E., 2005. Genome-wide profiling of stored mRNA in Arabidopsis thaliana seed germination: epigenetic and genetic regulation of transcription in seed. Plant J. 41, 697–709.

Pedersen, B.S., Quinlan, A.R., 2018. Mosdepth: quick coverage calculation for genomes and exomes. Bioinformatics 34, 867–868. https://doi.org/10.1093/bioinformatics/btx699

Rubio, V., Linhares, F., Solano, R., Martín, A.C., Iglesias, J., Leyva, A., Paz-Ares, J., 2001. A conserved MYB transcription factor involved in phosphate starvation signaling both in vascular plants and in unicellular algae. Genes Dev. 15, 2122–2133.

Schmid, M., Davison, T.S., Henz, S.R., Pape, U.J., Demar, M., Vingron, M., Schölkopf, B., Weigel, D., Lohmann, J.U., 2005. A gene expression map of Arabidopsis thaliana development. Nat. Genet. 37, 501–506.

Takahashi, D., Kawamura, Y., Uemura, M., 2016. Cold acclimation is accompanied by complex responses of glycosylphosphatidylinositol (GPI)-anchored proteins in Arabidopsis. J. Exp. Bot. 67, 5203–5215.

Tian, Y., Shi, C., Malo, C.U., Kwatcho Kengdo, S., Heinzle, J., Inselsbacher, E., Ottner, F., Borken, W., Michel, K., Schindlbacher, A., 2023. Long-term soil warming decreases

microbial phosphorus utilization by increasing abiotic phosphorus sorption and phosphorus losses. Nat. Commun. 14, 864.
Waese, J., Fan, J., Pasha, A., Yu, H., Fucile, G., Shi, R., Cumming, M., Kelley, L.A., Sternberg, M.J., Krishnakumar, V., 2017. ePlant: visualizing and exploring multiple levels of data for hypothesis generation in plant biology. Plant Cell 29, 1806–1821.
Wang, H., Xu, Q., Kong, Y.-H., Chen, Y., Duan, J.-Y., Wu, W.-H., Chen, Y.-F., 2014. Arabidopsis WRKY45 transcription factor activates PHOSPHATE TRANSPORTER1; 1 expression in response to phosphate starvation. Plant Physiol. 164, 2020–2029.
Wang, Q., Qin, Q., Su, M., Li, N., Zhang, J., Liu, Y., Yan, L., Hou, S., 2022. Type one protein phosphatase regulates fixed-carbon starvation-induced autophagy in Arabidopsis. Plant Cell 34, 4531–4553.
Wang, S., Ciais, P., Reich, P.B., Cescatti, A., Ellsworth, D.S., Janssens, I.A., Sardans, J., Luo, Y., Smith, N.G., Du, E., 2025. Phosphorus constrains global photosynthesis more than nitrogen does. Nat. Ecol. Evol. 1–11.
